# TIR-like NADases act in bacterial immunity and the RNA vault

**DOI:** 10.64898/2026.05.01.722283

**Authors:** Adam Osinski, Benjamin Mayro, Victor A. Lopez, Jason Schrad, Hannah Choi, Diana R. Tomchick, Krzysztof Pawłowski, Kevin Forsberg, Sarah H. Shahmoradian, Vincent S. Tagliabracci

## Abstract

Across all domains of life, organisms exploit NAD^+^ metabolism as a central line of defense against invading pathogens. Here, we show that domain of unknown function 4062 (DUF4062) is a widespread family of TIR-like NADases that hydrolyze NAD^+^ to ADP-ribose and nicotinamide. In bacteria, DUF4062 homologs form a previously unrecognized antiphage defense system, which we name Swarożyc, that assembles with the phage portal into a supramolecular NADase complex to induce abortive infection. In eukaryotes, DUF4062 is found in TEP1, which we demonstrate functions as an active NADase within the RNA vault, an enigmatic organelle-like structure. Single-particle cryo-electron microscopy reveals ADP-ribose bound within the shoulder of both reconstituted and human brain vaults, while cryo-electron tomography positions TEP1 along the central axis at the shoulder. Thus, TEP1, like bacterial Swarożyc, functions by depleting NAD^+^, providing new insight into the long-standing mystery of vault function.

## Introduction

Bioinformatic analyses of the vast protein sequence space have revealed potential functions for previously uncharacterized proteins. By integrating evolutionary relationships, conserved motifs, and sequence and structural similarities to known enzymes, these approaches uncover distant homologies that can be used to predict biochemical activities for experimental validation. The AlphaFold revolution^1,2^, which enables accurate structural modeling for most proteins, has further expanded this capability by revealing deep evolutionary relationships between enzyme families even when their amino acid sequences have diverged beyond the detection limits of the most advanced computational methods.

We have previously used bioinformatic methods to uncover new members of the kinase superfamily^3–6^. We reasoned that a similar strategy could be applied to the Toll/Interleukin Receptor (TIR) domain superfamily to uncover hidden functional diversity, catalytic versatility, or new biology. TIR domains, once thought to function solely as protein-protein interaction modules in innate immune signaling, are now recognized as a family that includes active enzymes^7^. Enzymatically active TIR domains mediate immunity by either depleting NAD^+^ to trigger altruistic cell death or generating small-molecule signals that activate downstream defense pathways^7,8^. TIR domains catalyze diverse reactions that utilize NAD^+^ as a substrate, including NAD^+^ hydrolysis^9–18^, cyclic ADP-ribose formation^19,20^, and production of ADP-ribose derivatives that act as second messengers in bacterial^21–24^ and plant^25–28^ immunity. TIR activation requires oligomerization, which aligns multiple TIR domains in the correct orientation to form the NAD^+^ binding _site10,11,16,19,29,30._

Here, we identify DUF4062 as a conserved family of TIR-like NADases. In bacteria, DUF4062 functions in antiphage defense, acting as the executioner by depleting NAD^+^ to block phage propagation. In humans, DUF4062 is found in TEP1, which we show functions as an active NADase within the RNA vault, a mysterious eukaryotic cellular structure^31^.

## Results

### DUF4062 is a new family of TIR-like domains

To uncover new members of the TIR domain family, we combined sequence and structural similarity searches^32^ and identified DUF4062, which shows distant sequence similarity to known TIR and nucleoside 2-deoxyribosyltransferase (NDT) domains (**Figures 1A and S1A**). Nevertheless, it is predicted to adopt a three-dimensional structure that closely resembles canonical TIR domains, such as SARM1 (**Figure 1B**). Sequence alignments identified a conserved Glu as a likely catalytic residue, corresponding to the known catalytic site of TIR NADases (**Figure S1B, 1C**). DUF4062 often co-occurs with auxiliary domains (**Figures S1D, 1E**), consistent with the modular architecture of TIR-containing proteins, which often rely on accessory domains for signal sensing, oligomerization, and TIR activation^7^.

**Figure 1.**
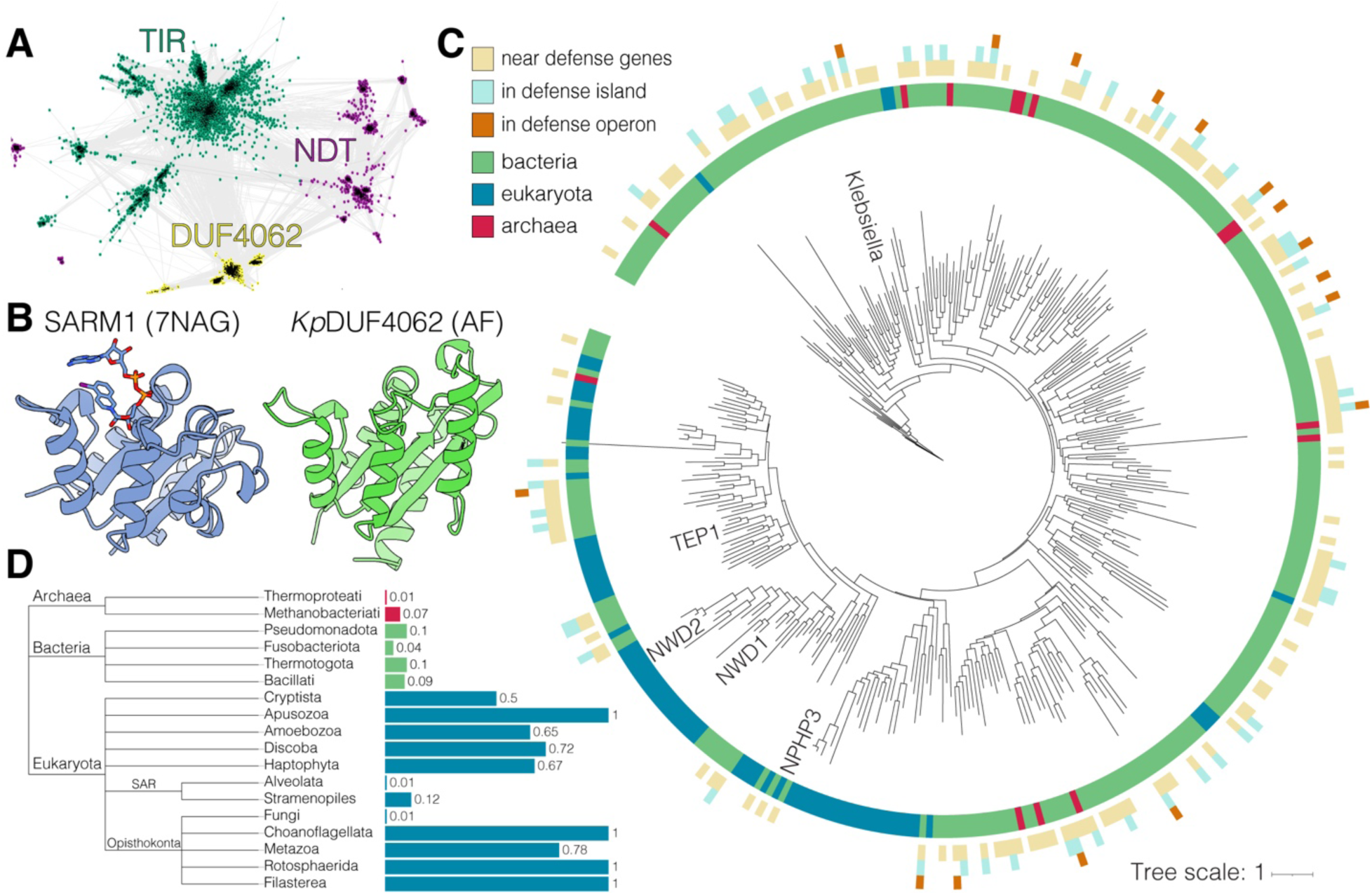
DUF4062 is a new family of TIR-like domains found in eukaryotes and bacterial antiphage defense islands. **(A)** CLANS sequence similarity network of representative TIR-like and NDT superfamily proteins (Pfam clans CL0173/STIR and CL0498/Nribosyltransf; BLAST E ≤ 0.01). Each point represents a single protein sequence, and distances reflect relative sequence similarity. **(B)** Structures of the human SARM1 TIR domain (PDB: 7NAG; left) and *K. pneumonia* DUF4062 domain (*Kp*DUF4062 UniParc: UPI001927123B; AlphaFold; right). SARM1 ligand 1AD is shown as sticks. **(C)** Phylogenetic tree of 345 representative DUF4062 domains. Branches highlight DUF4062 loci that neighbor defense genes (light beige), reside within antiphage defense islands (light teal), or occur in defense operons (orange). Colors denote taxonomic origin: bacteria (green), eukaryotes (blue), and archaea (red). **(D)** Prevalence of DUF4062 domains across analyzed genomes from selected taxa. Archaea (red), Bacteria (green), and Eukaryotes (teal). See also **Figure S1**

### DUF4062 is found in bacterial antiphage defense islands

Prokaryotic DUF4062 homologs display a broad but patchy distribution and are found across 26 bacterial phyla where they occur in ∼1–15% of analyzed reference genomes (**Figures 1C, 1D, and S1F**). In bacteria, ∼20% of DUF4062 homologs reside within predicted antiphage defense islands, and over half neighbor at least one homolog of a known defense gene (**Figure 1C**). Some are also found in operons together with homologs of characterized defense genes (**Figure S1G**). Notably, identical DUF4062 protein sequences can appear in diverse genera, often neighboring different phage defense systems. These observations suggest that DUF4062-containing proteins spread by horizontal gene transfer and may function in bacterial immunity.

### Bacterial DUF4062 mediates antiphage defense

We selected five DUF4062-containing proteins, that have similar C-terminal auxiliary domains (**Figure 2A**) and lie within bacterial defense islands (**Figure 2B**), expressed them in *Escherichia coli* BW25113 cells and challenged them with a subset of the BASEL phage library^33^. We observed strong and broad defense against the tested phages, most notably when DUF4062 proteins from *Klebsiella pneumoniae* and *Enterobacteriaceae* were expressed (**Figures 2C, 2D and S2A**). Phage defense required the active site E97 of *K. pneumoniae* DUF4062 and its auxiliary domain (**Figures 2E and S2B**). We named this newly discovered defense system Swarożyc (Swz), after a Slavic god of fire.

**Figure 2.**
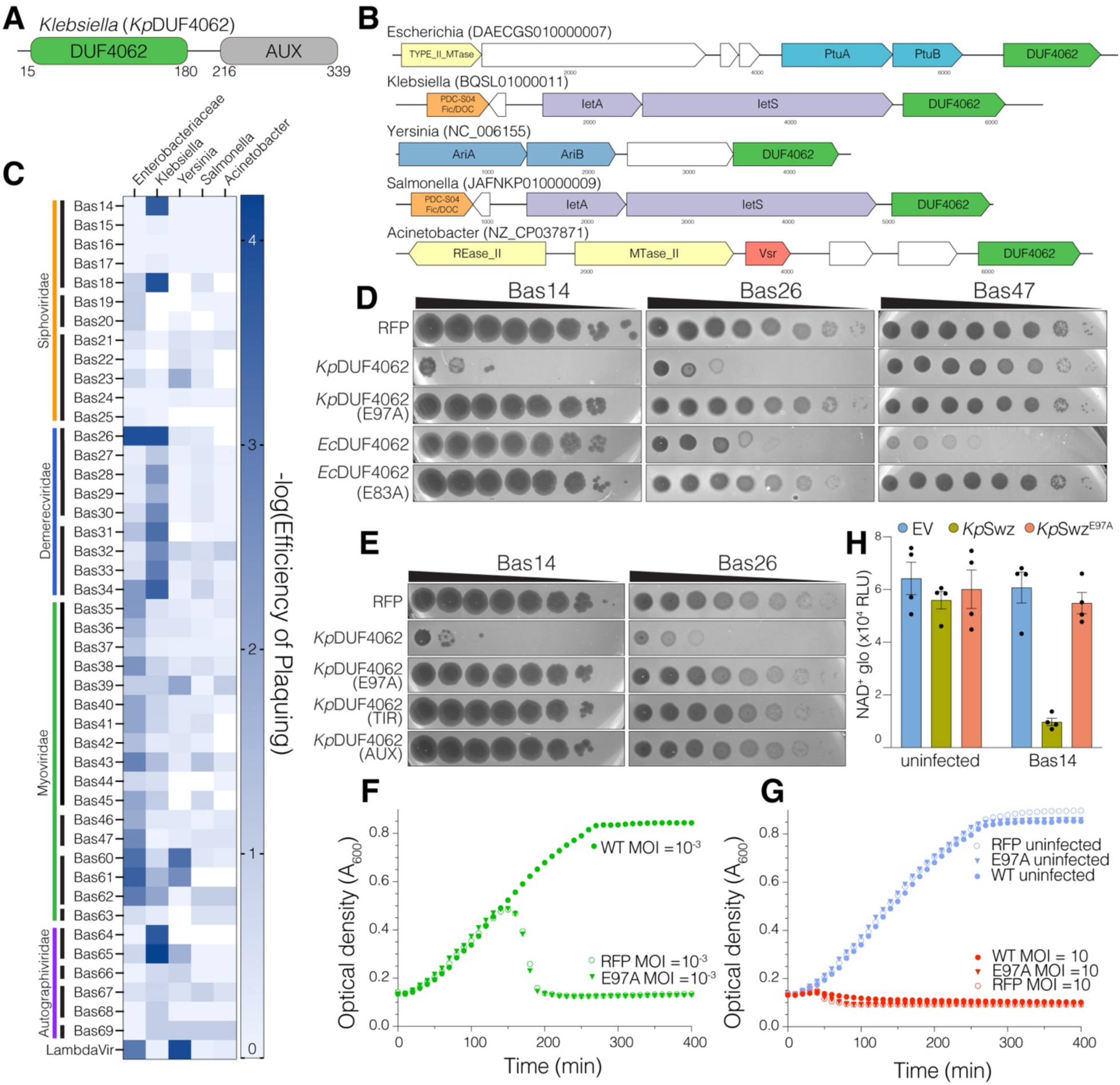
DUF4062 mediates antiphage defense by depleting NAD^+^ and blocking phage propagation. **(A)** Domain organization of a DUF4062-containing protein from *K. pneumoniae* with DUF4062 in green and auxiliary domain (AUX) in grey. **(B)** Genomic neighbourhood of 5 selected DUF4062 proteins highlighting known or predicted defense genes. DUF4062 is in green and each of the other colors denotes a different characterized defense system (all protein sequence accessions are located in the Supplemental Materials). **(C)** Heatmap depicting the efficiency of plaquing (EOP) of DUF4062-containing proteins expressed in *E. coli* BW25113 against a subset of the BASEL library ^33^. **(D)** Phage plaques from ten-fold serial dilutions of Bas14, Bas26 and Bas47 on lawns of BW25113 cells expressing RFP or the indicated *Kp*DUF4062^16-C^ and *Ec*DUF4062 (*Enterobacteriaceae/E. coli* DUF4062) constructs. Images are representative of 3 biological replicates. **(E)** Phage plaques from ten-fold serial dilutions of Bas14 or Bas26 on lawns of BW25113 cells expressing RFP or the indicated *Kp*DUF4062^16-C^ constructs. Images are representative of 3 biological replicates. **(F, G)** Growth of *E. coli* BW25113 expressing RFP or the indicated *Kp*DUF4062 constructs (renamed *Klebsiella pneumoniae* Swarożyc: *Kp*Swz) after infection with Bas14 at MOI 10^-3^ **(F)** or 10 **(G**), alongside uninfected controls. Curves are representative of 3 biological replicates. **(H)** Graph showing NAD^+^ levels in uninfected or Bas14-infected *E. coli* expressing RFP control, *Kp*Swz^16-C^ or the inactive E97A mutant. See also **Figure S2**

TIR domain–containing defense systems often act through abortive infection, in which phage-infected cells undergo altruistic cell death to protect the bacterial population^34,35^. To determine whether *K. pneumoniae* Swarożyc (*Kp*Swz) induces an abortive defense, we infected liquid cultures expressing *Kp*Swz with phage Bas14 at high (10) and low (10^-3^) multiplicities of infection (MOI). *Kp*Swz rescued growth at low MOI (**Figure 2F)**, whereas high MOI caused culture collapse (**Figure 2G**) and reduced phage titers (**Figures S2C, S2D**). Moreover, NAD^+^ levels were reduced upon phage infection in cells expressing WT *Kp*Swz, but not the E97A mutant (**Figure 2H**). Thus, *Kp*Swz protects the bacterial population by depleting NAD^+^ to trigger abortive infection, thereby blocking phage propagation.

### Structural insights into the *Kp*Swz TIR-like domain

We determined a cryo-EM structure of *Kp*Swz^15-C,E97A^, revealing a symmetric TIR-like core dimer resembling a pair of flowers with intertwined stems (**Figures 3A, S3 and Table S1**). These dimers assemble into tetrameric and hexameric bouquets forming a short antiparallel fiber-like structure with a helical turn of ∼53° and a rise of ∼17 Å (**Figures 3B and S3**). The TIR-like domain consists of a five-stranded, parallel β-sheet flanked by five α-helices in a β-α-β-α repeating topology **(Figure S3C).** The core dimer interface is formed between the αA and αB helices (AB interface) (**Figure 3C**), which is distinct from the AE interface seen in other TIR domains, such as SARM1 (**Figure 3D)**^36^. Interactions at the AB interface push the regulatory BB loops, which link βB to αB, into the TIR-like active sites, where they occlude the catalytic pocket and lock the enzyme into an inactive conformation (**Figure 3C**). Beneath the TIR-like domains, strands from both αE helices form a small antiparallel β-sheet that intertwines with their partner (**Figure 3E**) before extending into a pair of helical stalks.

**Figure 3.**
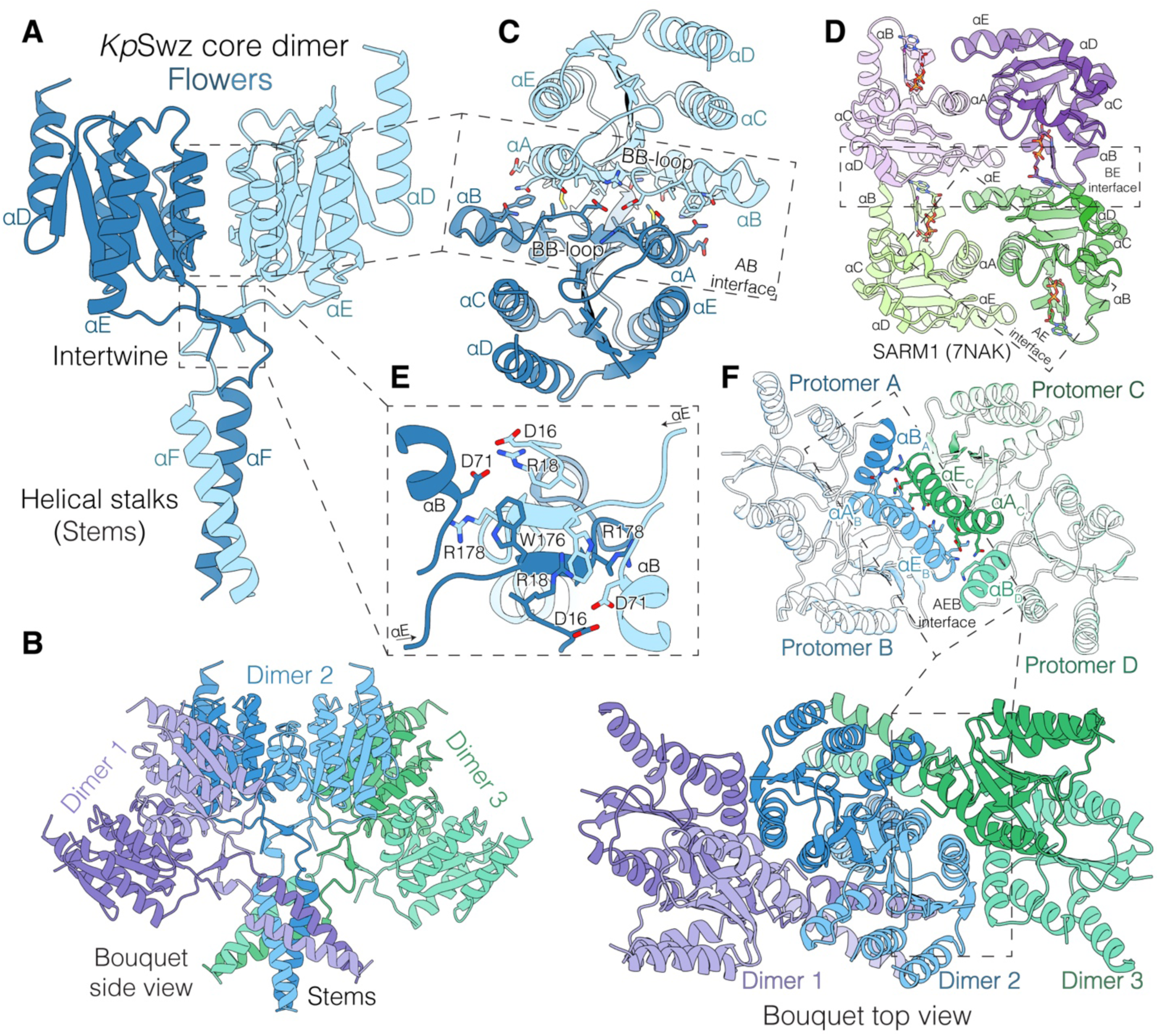
The *Kp*Swz TIR-like domain adopts a distinct, bouquet-like higher-order assembly. **(A)** Model of the *Kp*Swz^15-C, E97A^ core dimer (modeled residues 15–205, with gaps) shown as a cartoon. Shades of blue represent the two monomers. The TIR-like domains, shaped like flowers, intertwine through a small β-sheet into two helical stalks that resemble flower stems. **(B)** Side (left) and top (right) orthographic projections of the hexameric *Kp*Swz mini “fiber”. Each laterally assembled core dimer is shown in two different shades of violet, blue, or green. The side view is perpendicular to the “fiber” axis, whereas the top view represents an arbitrary rotation. **(C)** Top view depicting the AB interface of the *Kp*Swz core dimer. α-helical elements are labeled, and interface residues are shown as sticks. **(D)** Cartoon depiction of the active TIR domain of SARM1 bound to the inhibitor 1AD (PDB: 7NAK; ^69^). The symmetric dimerization interface (AE) and the asymmetric oligomerization interface (BE) are indicated. Molecules of 1AD are shown as sticks. **(E)** Zoomed-in view of the core dimer showing the two chains intertwining beneath the TIR-like domains, stabilizing the dimer and exchanging the αF helices of the stalk region. The interaction is asymmetric, with R18 participating either in salt bridges with D71/D16 or in a cation–π interaction with W176. **(F)** Top view depicting the AEB oligomerization interface of the *Kp*Swz tetramer. Interacting helices are highlighted, and interface residues are shown as sticks. See also **Figure S3**

The *Kp*Swz core dimers further assemble into tetrameric and hexameric bouquets (dimer/trimer of dimers) (**Figure 3B**). The αA and αE helices from two core dimers come together (αAE_B_ and αAE_C_; subscripts indicate protomers), while αB_A/D_ interact with αE_C/B_ to form the AEB oligomerization interface (**Figure 3F**). The helical stalks also mediate inter-dimer contacts, with stem helices (αF) from protomers A and D forming the core of this interaction (**Figure 3B**). In the tetramer, each chain contacts all others, generating a distinct higher-order assembly in which the AB and AEB interfaces enforce a head-to-head, autoinhibited architecture.

### *Kp*Swz is a portal-activated NADase

Bacterial immune systems are typically maintained in an inactive state until phage infection is detected. In response, phages frequently evolve resistance by mutating the proteins recognized by these systems, thereby evading detection and preventing immune activation^37^. When challenged with *Kp*Swz, Bas14 phage consistently formed plaques that evaded immune defense. To identify the phage-encoded triggers of *Kp*Swz, we isolated these immune escapers and sequenced their genomes (**Figures 4A and S4A**). All escapers carried mutations in *bas14_0003*, which encodes the portal protein (portal^Bas^^14^). Phage portal proteins mediate capsid assembly and genome transfer in tailed dsDNA bacteriophages^38^ and are known triggers of bacterial immune defense systems^39–41^. Therefore, we monitored the growth of *E. coli* BW25113 cultures expressing portal^Bas14^ and *Kp*Swz. Expression of WT portal^Bas14^, but not 3 of the 4 escaper mutants led to *Kp*Swz-mediated cell death (**Figures 4B, 4C and S4B).**

**Figure 4.**
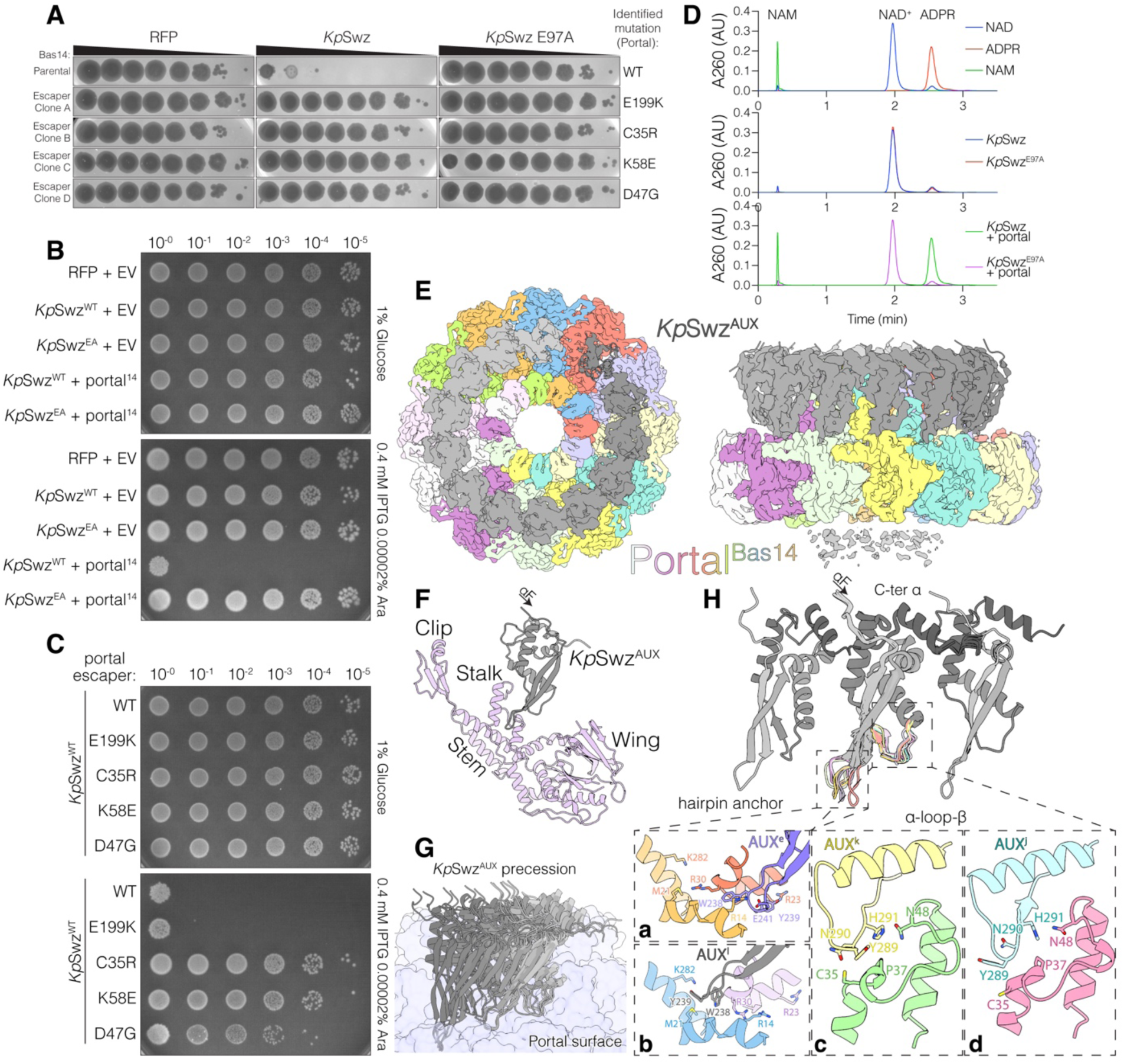
*Kp*Swz is a portal activated NADase. **(A)** Phage plaques from ten-fold serial dilutions of Bas14 (parental) or escaper mutants (Clones A-D) on lawns of BW25113 cells expressing RFP, *Kp*Swz^16-C^ (*Kp*Swz) or the inactive E97A mutant. Images are representative of 3 biological replicates. The mutations are shown on the right. **(B, C)** Growth assays of ten-fold serial dilutions of *E. coli* BW25113 expressing RFP, *Kp*Swz^16-C^, or the inactive E97A mutant (*Kp*Swz^EA^) in combination with empty vector (EV) or Bas14 portal (portal^14^) **(B)**, or with Bas14 portal escaper mutants **(C)**. *Kp*Swz expression was induced with IPTG, and portal expression was induced with arabinose (Ara). (D) HPLC-UV_260_ traces depicting nicotinamide (NAM), NAD^+^ and ADP-ribose standards (top), and reaction products generated after incubation of NAD^+^ with *Kp*Swz^15-C^ (*Kp*Swz, 0.05 mg/mL) alone (middle) or with *Kp*Swz in the presence of excess portal^Bas14^ (bottom). (E) Top (upper) and side (lower) orthographic projections of the symmetry-relaxed *Kp*Swz–portal^Bas14^ complex. Coulombic potential maps are overlayed with the atomic model shown as a cartoon, visible in regions of poor density. The 12 portal^Bas14^ monomers are uniquely colored, and the 13 *Kp*Swz^AUX^ domains in the crescent are shown in a grey gradient. Residual density for the unresolved crown region of the portal is visible as grey “dust” at the bottom of the panel. (F) Cartoon depiction of a single portal (light purple)–*Kp*Swz^AUX^ (grey) heterodimer. The portal wing, stem, and clip regions are indicated. (G) Cartoon depiction of the heterogeneous Portal^Bas14^–*Kp*Swz^AUX^ interface. Each *Kp*Swz^AUX^ domain interacts differently with the portal. The precession of *Kp*Swz^AUX^ around the portal ring (violet surface) is illustrated by overlaying *Kp*Swz^AUX^ monomers from all 13 portal–*Kp*Swz^AUX^ pairs. (H) Cartoon representation of three neighboring *Kp*Swz^AUX^ domains, highlighting examples of their interactions with portal^Bas14^ within the hairpin anchor (**panels a, b**) and the α-loop-β module (**panels c, d**). Each chain is shown as a gray gradient, darkening toward the C-terminus, all chains are uniquely colored in the example panels. The side-by-side assembly is mediated by the rigid HTH/winged helical core, with the C-terminus tucked beneath the N-terminal portion of the domain. All 13 *Kp*Swz^AUX^ monomers were overlaid on the middle monomer. See also **Figures S4-S6**

To determine whether portal^Bas14^ directly activates *Kp*Swz, we incubated *Kp*Swz with portal^Bas14^ and NAD^+^ and analyzed reaction products by HPLC-UV. NAD⁺ was converted to ADP-ribose and nicotinamide, and this activity required the active-site residue E97 and portal^Bas14^ (**Figures 4D and S4C**). Thus, portal^Bas14^ is sufficient to trigger *Kp*Swz-induced cell death by activating *Kp*Swz NADase activity.

### *Kp*Swz and portal^Bas14^ form a supramolecular NADase complex

Portal^Bas14^ formed a stable complex with *Kp*Swz on size-exclusion chromatography (**Figures S4D**). We determined the cryo-EM structure of *Kp*Swz^15-C,E97A^ bound to portal^Bas14^ (**Figures 4E, S5 and Table S1**). Although the TIR-like domain was not resolved, we observed strong density for the *Kp*Swz auxiliary domain (*Kp*Swz^AUX^). The portal assembles into a dodecameric ring, with *Kp*Swz^AUX^ engaging the stalk that projects from the portal stem (**Figures 4E and 4F**). In asymmetric reconstructions of the complex, obtained by relaxing the C12 symmetry, 13 *Kp*Swz^AUX^ subunits assemble into an incomplete ring, forming a crescent around the portal stem and clip (**Figures 4E and 4F**). Each *Kp*Swz^AUX^ monomer bound portal^Bas14^ in a position slightly offset from its neighbors, creating a heterogeneous interface around the ring (**Figure 4G**). *Kp*Swz^AUX^ monomers adopt a helix-turn-helix (HTH)/winged fold and associate side by side through contacts in the α-helical core and interactions between their N- and C-termini (**Figure 4H, upper**).

*Kp*Swz^AUX^ monomers engage the portal through flexible portal-recognition modules that include a hairpin anchor and an α-loop-β motif (**Figure 4H, lower**). The conformations of both modules adjust to the portal surface and change from one unit to the next. A subset of monomers function as anchors, with W238 and Y239 at the hairpin tip buried in a pocket between the wing and stalk domains. Depending on their position within the crescent, W238 and Y239 insert between portal residues R14 and R30 (**Figure 4H, panel a)**, extend toward M21 and K282 (**Figure 4H, panel b)**, or are disordered. Similarly, the *Kp*Swz^AUX^ α-loop-β motif forms heterogeneous interactions with the helical stalk of the portal surrounding C35 (**Figure 4H, panels c, d**). Notably, the C35R Bas14 escaper mutation (**Figure 4A**) fails to activate *Kp*Swz when co-expressed in *E. coli* (**Figure 4C**).

Sequencing of the portal gene from 17 Bas14 escaper plaques revealed that all identified escaper mutations localize at or near the *Kp*Swz^AUX^-binding interface (**Figure S6A**). While their exact effects are unclear, they likely alter the position or conformation of the portal^Bas14^ stalk to evade detection by *Kp*Swz. However, the E199K portal^Bas14^ mutation is not at the *Kp*Swz^AUX^-binding interface and likely reflects a distinct escape mechanism, as it remains capable of triggering bacterial cell death when co-expressed with *Kp*Swz (**Figure 4C**).

Our structural analysis suggests that the flexibility of the *Kp*Swz^AUX^-portal^Bas14^ interaction allows up to 13 *Kp*Swz^AUX^ protomers to bind a dodecameric portal^Bas14^, forming a supramolecular complex and activating *Kp*Swz NADase activity. Although density for the TIR domain in the *Kp*Swz-portal^Bas14^ assemblies was not observed, we propose an activation mechanism whereby portal^Bas14^ engagement with *Kp*Swz^AUX^ pulls on the helical stalks, aligns them, and ultimately repositions the TIR-like domains into an active conformation ^42^. AlphaFold modeling supports this proposed activation mechanism (**Figures S6B and S6C**).

Together, our results suggest that bacterial DUF4062-containing proteins function in antiphage defense by sensing phage infection, thereby triggering structural rearrangements, activating DUF4062 NADase activity, and inducing abortive infection.

### Human TEP1 is an active NADase

Unlike in bacteria, DUF4062 is widely conserved across diverse eukaryotic lineages, including Metazoa (**Figures 1C and 1D**). Building on our findings in bacteria, we sought to determine whether DUF4062 NADase activity is conserved in eukaryotes.

In human NWD1, NWD2, TEP1, and NPHP3, the DUF4062 domain lies adjacent to a NACHT ATPase (**Figure 5A**), a domain that facilitates oligomerization and is common to innate immune proteins across all domains of life^15^. These proteins contain an apoptotic protease-activating factor 1 (APAF-1)–like helical domain (α), and several also have multiple WD40 repeats. A similar domain organization is present in APAF-1 (**Figure 5A**), which senses cytochrome c release from mitochondria to induce apoptosis^43^.

**Figure 5.**
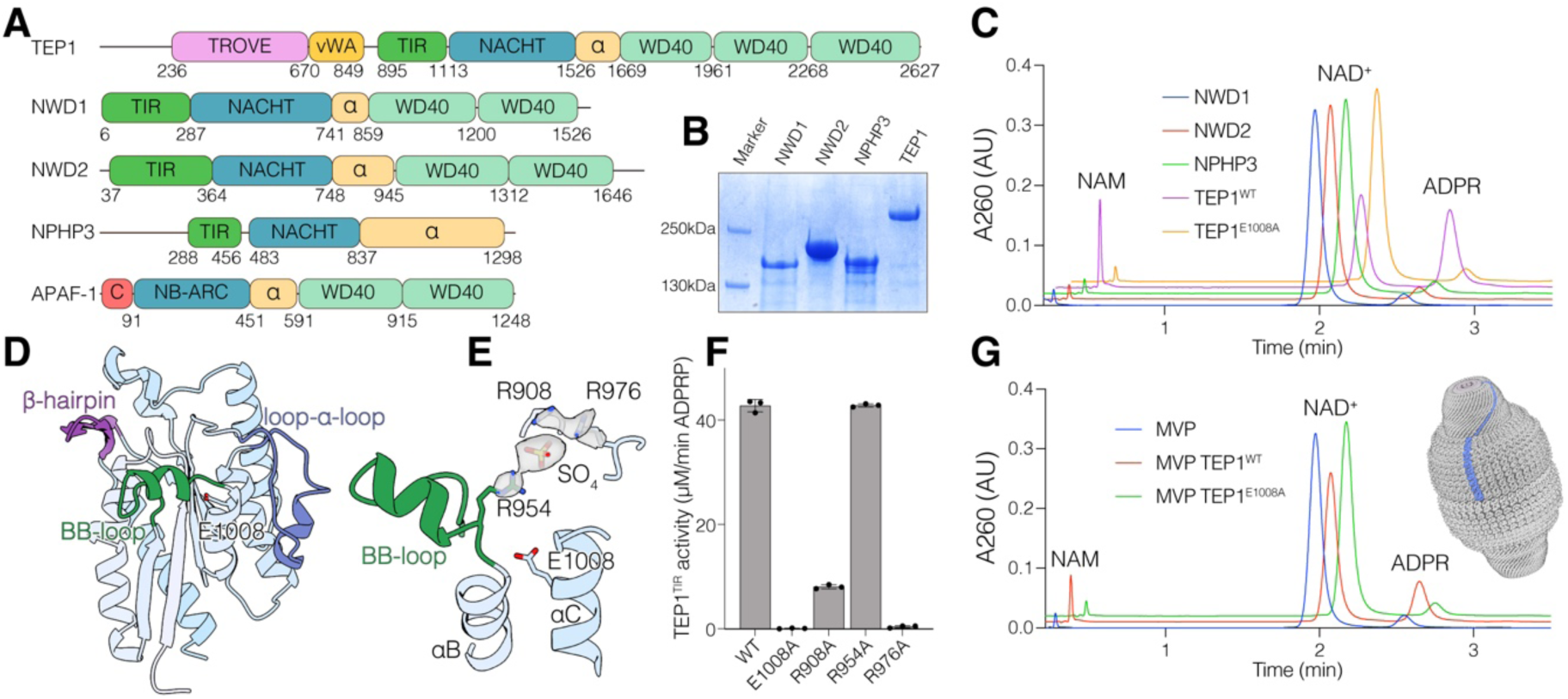
Human TEP1 is an active NADase. **(A)** Domain architecture of human DUF4062-containing proteins TEP1, NWD1, NWD2, and NPHP3. APAF1 is shown for comparison. (Telomerase, Ro and vault, TROVE; von Willebrand A, vWA; CARD, C; alpha helical, α). **(B)** SDS-PAGE and Coomassie-stained gel showing purified NWD1, NWD2, NPHP3, and TEP1. **(C)** HPLC traces depicting the reaction products generated after incubation of NAD^+^ with TEP1, NWD1, NWD2 and NPHP3. Protein preparations were used at 0.5 mg/mL. **(D)** Cartoon depiction of the TEP1^TIR^ domain, highlighting unique structural features, including the extended BB loop with an embedded α-helix (green), a loop–helix–loop motif extending from αC (loop–α–loop; blue), and a β-hairpin preceding αE (purple). **(E)** Zoomed-in view of the TEP1^TIR^ active site, highlighting the BB loop (green) and three Arg residues pointing toward a putative sulphate ion. The catalytic E1008 is shown, and sulphate-interacting residues are depicted as sticks. 2Fo-Fc map is presented as a surface. **(F)** Graph depicting NADase activity of TEP1^TIR^ and select active site mutants (1 μM, 15 min, ambient temperature). Reaction products were analyzed as in **Figure S4C**. **(G)** HPLC traces depicting the reaction products generated after incubation of NAD^+^ with the RNA vault containing either TEP1 (red) or the E1008A mutant (green). Vault preparations were used at 0.75 mg/mL. See also **Figures S7 and S8**

We purified full-length human NWD1, NWD2, NPHP3, and TEP1 (**Figure 5B**), incubated them with NAD^+^ and NADP^+^, and analyzed reaction products by HPLC. Only TEP1 hydrolyzed NAD^+^ and NADP^+^, producing ADP-ribose or ADP-ribose phosphate (ADPRP) and nicotinamide (NAM) with activity dependent on the active site E1008 (**Figures 5C and S7A**). The isolated DUF4062 domain (TEP1^886–1113^, hereafter TEP1^TIR^) was sufficient to hydrolyze NAD^+^ and NADP^+^ (**Figures S7B and S7C)**. Unlike other TIR-domain proteins such as SARM1, AbTIR, or HopAM1, TEP1^TIR^ did not generate additional ADP-ribose derivatives (**Figure S7D**). When using 1 μM TEP^TIR^, the reaction was slow (*k*_cat_ = 173.4 min^-1^), with an apparent K_m_ of ∼6.9 mM for NADP^+^ and undeterminable for NAD^+^ (**Figures S7E and S7F**). However, the specific activity of TEP1^TIR^ increased proportionally with its concentration (**Figure S7G**), suggesting activation via concentration-dependent oligomerization. This is consistent with oligomerization-driven activation observed for other TIR NADases^11^. After SARM1^9^, TEP1 represents the second human TIR domain known to have catalytic activity.

We determined the crystal structure of human TEP1^TIR^, which adopted a canonical TIR-like fold (**Figure 5D and Table S2**). Electron density quality varied throughout the asymmetric unit, with refined B-factors ranging from ∼20-150 Å² and progressively worsening from αA/αE toward αB (**Figure S8**). The BB loop folds into the active site, suggesting an inactive conformation (**Figure 5D**), and no relevant oligomerization was observed in the crystal lattice. Compared to the *Kp*Swz TIR-like domain, TEP1^TIR^ contains an extended BB loop with an embedded, highly charged α-helix, a flexible loop-helix-loop motif extending from αC, and a β-hairpin preceding αE (**Figure 5D**). Within the BB loop, R954 projects into the active site and, along with R908 and R976, coordinates electron density consistent with a sulfate ion present in the crystallization buffer (**Figure 5E**). Alanine substitutions of R908 and R976 reduce TEP1^TIR^ activity (**Figure 5F**). These unique structural inserts distinguish TEP1^TIR^ from the *Kp*Swz TIR-like domain.

### TEP1 is active in the RNA vault

TEP1 was originally identified as a telomerase-interacting protein, suggesting a role in telomerase activity^44^; however, genetic studies show that TEP1 is dispensable for telomerase function and telomere length maintenance in vivo^45^. Instead, TEP1 is thought to function within the RNA vault, a ∼65 nm capsid-like ribonucleoprotein particle first identified in rat liver microsomal fractions^46,47^. The vault consists of two dome-shaped halves, each assembled from 39 copies of the major vault protein (MVP)^48^, and commonly encloses TEP1, the ADP-ribosyltransferase PARP4 and small non-coding vault RNAs ^49,50^. Despite extensive study, its function remains unknown^31^.

To test whether TEP1 is catalytically active within the vault, we reconstituted human vaults with MVP, and TEP1 in HEK293F cells. Purified vaults containing TEP1, but not the E1008A mutant, displayed NADase and NADPase activities (**Figures 5G and S7H**). Thus, NAD(P)ase activity within the vault is conferred by the TEP1 TIR-like domain.

### Cryo-EM analysis of reconstituted and native RNA vaults reveals a bound ADP-ribose

To investigate how TEP1 catalyzes NAD(P)⁺ hydrolysis in the vault, we performed single-particle cryo-EM analysis (SPA) of reconstituted human vaults containing TEP1. We added the NAD^+^ analog benzamide adenine dinucleotide (BAD) to occupy the TEP1 TIR-like domain, and AMP-PNP to occupy the TEP1 NACHT domain. Although this approach yielded high resolution RNA vault maps (**Figure S9 and Table S3**), the TEP1 density could not be resolved.

Similarly, SPA of reconstituted vaults containing TEP1 and PARP4 (**Figures S10 and S11 and Table S4**) in the presence of NADP^+^ failed to resolve the internal proteins. However, sub-particle extraction yielded a ∼2.9 Å reconstruction of the PARP4 MVP-interacting (MINT) domain bound to the vault wall, complementing a recent report^51^ (**Figures S10 and S11C**). Exhaustive 3D classification indicated an average of ∼20 PARP4 molecules per reconstituted vault (**Figure S10B**). Further sub-particle analysis resolved the symmetry-mismatched cap to ∼1.9 Å (C13 symmetry) (**Figure S10 and S11D**), in agreement with recent reports^52,53^.

To assess whether the reconstituted vaults had captured the added nucleotides, we performed focused refinement of the shoulder region, recently reported to bind adenine nucleotides: NAD⁺, ADP and ADP-ribose^51^. In vaults containing TEP1, BAD and AMP-PNP, BAD occupied the shoulder nucleotide-binding pocket (**Figure 6A, panel a**). Interestingly, in vaults containing TEP1 and PARP4 supplemented with NADP⁺, we observed density corresponding to ADP-ribose rather than the expected NADP⁺ or ADP-ribose phosphate (**Figure 6A, panel b**). The source of this ADP-ribose is unclear; it may result from TEP1-mediated hydrolysis of contaminating NAD⁺ in the sample or co-purified from the cells. In any event, its presence despite excess NADP⁺ suggests a preference for ADP-ribose binding to the shoulder of the vault.

**Figure 6.**
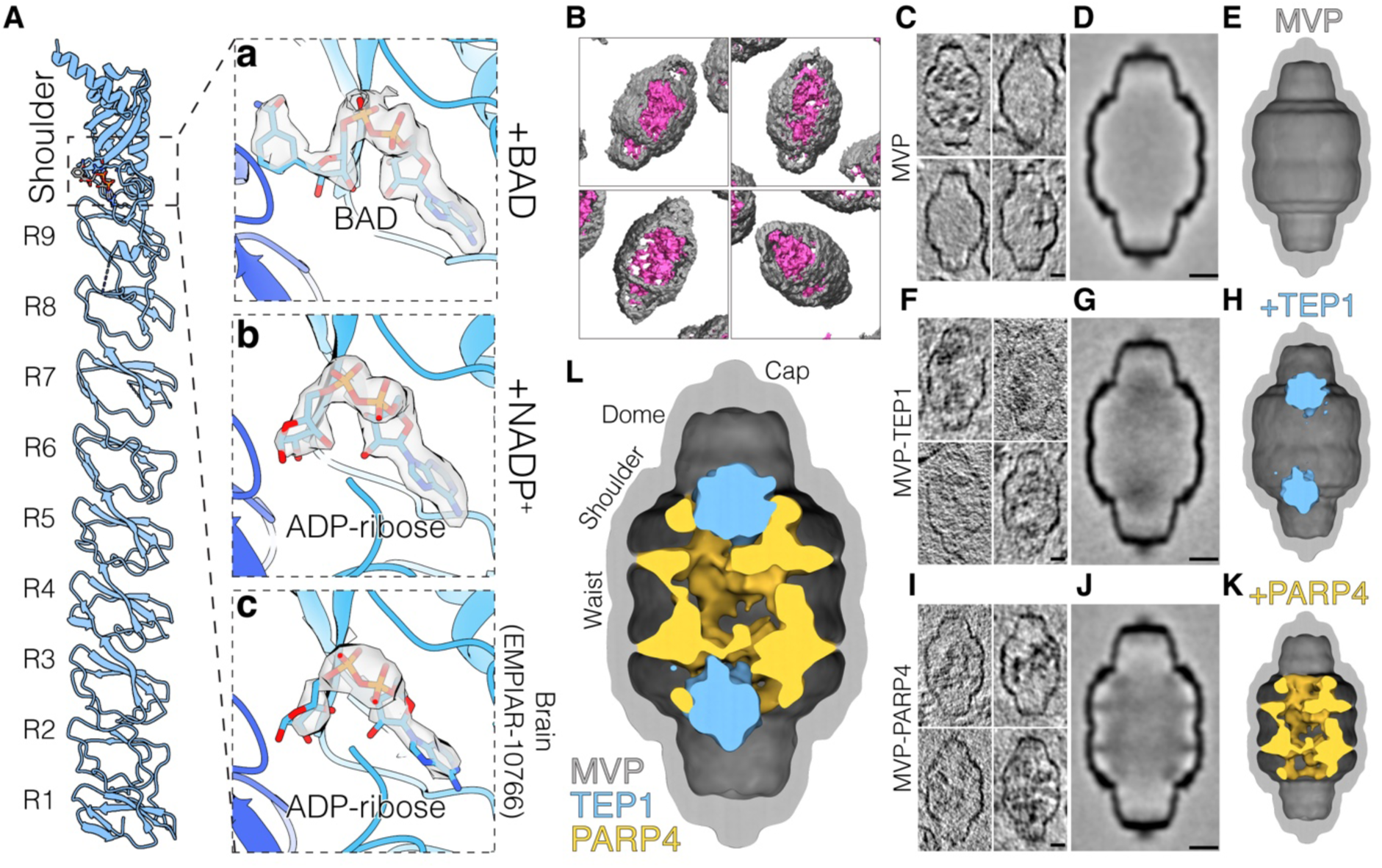
Structural analysis reveals ADP-ribose binding proximal to TEP1 within the vault. (A) Cartoon representation of a single MVP monomer, with repeats R1–R9 and the shoulder region labeled. Panels **a-c** shows the contents of the nucleotide-binding pocket in the three analyzed datasets (**a,** MVP/TEP1/BAD/AMP-PNP; **b,** MVP/TEP1/PARP4/NADP^+^; **c,** human brain EMPIAR-10766): BAD and ADP-ribose nucleotides are shown as sticks, along with the corresponding Coulombic densities. (B) Isosurface renderings of four reconstituted vault particles from a single tomogram. Segmentation was performed using Dragonfly with individual models trained for the vault exterior (grey) and interior (magenta). (C) 5nm-thick central slices of reconstituted, MVP-only vault particles extracted from high contrast tomograms and aligned using PEET. **(D-E)** Central slice (5 nm, **D**) and isosurface rendering (**E**) of the MVP subtomogram average, split along the central axis. Note that the material that appears in some of the particles in **(C)** is averaged out during averaging. **(F-H)** 5-nm thick central slices of MVP-TEP1 vault particles (**F**) and the subsequent subtomogram average (**G**) with the isosurface rendering depicted in (**H**). The MVP shell is depicted in grey and the interior density corresponding to TEP1 is colored blue. **(I-K)** 5-nm thick central slices of MVP-PARP4 vault particles (**I**) and the subsequent subtomogram average (**J**) with the isosurface rendering depicted in (**K**). The density corresponding to PARP4 is colored yellow. **(L)** Composite image with isosurface renderings of the MVP-TEP1 subtomogram average overlaid onto the MVP-PARP4 average. Note the minimal overlap between the densities corresponding to TEP1 and PARP4 in the overlay. Scale bars represent 10 nm. See also **Figures S9-S13**

A recent cryo-EM study determined structures of the RNA vault identified as a contaminant in the sarkosyl-insoluble fraction of frontal cortex from a human brain with Type I globular glial tauopathy^52,54^ (Dataset EMPIAR-10766). We reprocessed this dataset focusing on the shoulder region and observed an ADP-ribose molecule, despite, to the best of our knowledge, no nucleotides having been added (**Figures 6A, panel c, S12, and Table S4**). These results indicate that NAD⁺ hydrolysis products are natively present in human RNA vaults.

### Cryo-electron tomography reveals the internal protein architecture of reconstituted vaults

Given that vaults containing TEP1 exhibit NADase activity and that we observe its product, ADP-ribose, bound at the vault shoulder, we sought a direct structural readout of TEP1 localization within the vault and its proximity to the nucleotide-binding pocket.

Early studies proposed that TEP1 localizes to the cap^55^, but the supporting evidence is limited and the assignment remains uncertain. Therefore, we used cryo-electron tomography (cryo-ET) to visualize reconstituted human vaults. While tomograms of vaults containing MVP, TEP1, and PARP4 readily resolved the external shell, internal features showed limited contrast until Volta phase plate (VPP) imaging revealed reproducible internal densities distinct from the MVP shell (**Figures S13A and S13B**). Segmentation of individual particles identified recurring ultrastructural motifs within the vault body, consistent with a highly ordered internal organization rather than heterogeneous or freely diffusing cargo (**Figures 6B and S13C**).

To assess whether these internal features reflect component-specific localization, we performed subtomogram averaging on vaults reconstituted with MVP alone, MVP and TEP1, or MVP and PARP4. Vaults containing only MVP showed minimal internal density (**Figures 6C-E and S13D**). In contrast, TEP1-containing vaults displayed a distinct density along the central axis in the shoulder region (**Figure 6F-H and S13E**), while PARP4-containing vaults showed a broader peripheral density near the vault wall extending into the shoulder (**Figure 6I-K and S13F**), consistent with a recent report^51^. Comparison of these independent reconstructions showed that the TEP1-associated density occupies a cavity within the PARP4-associated region (**Figure 6L**). Because TEP1- and PARP4-containing vaults were reconstituted and analyzed separately, their complementary localization suggests that TEP1 and PARP4 occupy self-defined positions within the vault.

Previous cryo-ET analyses of intracellular vaults concluded that lumenal densities are not ordered and average out during subtomogram averaging^56^. Those studies analyzed native vaults containing heterogeneous cargo and endogenous cellular components. In contrast, our reconstituted systems isolate defined vault compositions and minimize combinatorial heterogeneity. Under these conditions, and using high-contrast cryo-ET, we observe reproducible, component-specific internal densities. These findings suggest that lumenal organization is context- and occupancy-dependent rather than intrinsically random.

Together, these data support a model in which MVP enforces a highly ordered internal architecture, positioning TEP1 near the central axis at the shoulder while restricting PARP4 to belts close to the shoulder and above the waist. The shoulder region of the vault concentrates the NADase activity of TEP1 and the ADP-ribosyltransferase activity of PARP4 near the MVP nucleotide-binding pocket. This spatial arrangement provides a structural framework to guide future studies aimed at elucidating the function of the vault.

## Discussion

In this work, we identify DUF4062 as a widespread family of TIR-like NADases present in both prokaryotes, where it functions as the executioner in an antiphage defense system, and in eukaryotes, where it hydrolyzes NAD⁺ to ADP-ribose and nicotinamide in the RNA vault.

Immune mechanisms are frequently shared between bacteria and humans; for example, homologs of eukaryotic immunity- and cell death-related genes have been identified as key mediators of bacterial antiphage defense. These include components of the cGAS–STING pathway^16,57^, caspases^24^, gasdermins^58,59^, viperins^60^, signal-transducing ATPases with numerous domains (STAND)^15^, SIR2^21,61,62^ and TIR domains^21,24,61^. Interestingly, vaults appear to be involved in innate immunity, as they are highly expressed in macrophages and dendritic cells^63^ and promote host resistance to *Pseudomonas aeruginosa*^63^ and influenza A virus^64^. Thus, we propose that the TIR-like domain in TEP1, like bacterial Swarożyc, may function as an executioner in a cell death pathway linked to innate immunity.

NWD1, NWD2, NPHP3 and TEP1 share a domain architecture reminiscent of other STAND ATPases, such as APAF-1^65^. APAF-1 contains an N-terminal caspase recruitment domain (CARD), an NB-ARC ATPase/oligomerization domain, and two WD40 propellers that fold back onto the ATPase to prevent oligomerization and subsequent activation. Cytochrome c binding to the WD40 domains relieves this inhibition, promoting nucleotide exchange and assembly of a heptameric APAF-1 ring that recruits and activates procaspase-9^66^. In NWD1, NWD2, and TEP1, DUF4062 occupies the equivalent position of the APAF-1 CARD domain (**Figure 5A)**. We speculate that the TEP1 WD40 domains respond to an unknown signal, which promotes NACHT ATPase oligomerization, leading to activation of the TIR-like domain.

While DUF4062 proteins are far more prevalent in eukaryotes, the broad yet patchy taxonomic distribution of prokaryotic DUF4062 homologs is consistent with the paradigm of defense systems functioning as a “community resource” within a broader pan-immune system, whose components are unevenly distributed across strains and propagated by horizontal gene transfer^67^. Similar to the diversity seen in Thoeris antiphage defense systems that use canonical TIR domains^68^, some DUF4062-containing proteins harbor additional enzymatic domains, whereas others occur in operons alongside diverse enzymes (**Figure S1**). Thus, the full spectrum of antiphage activities mediated by DUF4062 may involve mechanisms more complex than the NADase activity described here.

In summary, we identify DUF4062 as a previously unrecognized family of TIR-like NADases that function in bacterial antiphage defense. Its presence and activity in human TEP1 suggest that RNA vaults may employ a conserved NAD^+^-depleting mechanism in innate immunity. Further, our findings reinforce the idea that bacterial antiphage defense systems are powerful discovery systems for uncovering biochemical activities of human proteins.

## Acknowledgements

We thank members of the Tagliabracci laboratory for helpful discussions, Emma Choo for cryo-ET help, Alexander Harms for the BASEL phages and the BW strains, Helen Aronovich, James Chen, Yang Li, Dan Stoddard (UTSW Structural Biology Laboratory and the CEMF), Nicholas Spellmon, and Rui Yan (HHMI Janelia CryoEM Facility) for help with cryo-EM data collection. This work was funded by NIH Grants DP2GM137419, R35GM158265 (V.S.T.), DP2-AI154402 (K.J.F.) and K00CA264162 (B.M.), Welch Foundation Grants I-1911 (V.S.T.), the Endowed Scholars Program at UT Southwestern Medical Center (K.J.F. and V.S.T.), the Department of Defense Congressionally Directed Medical Research Programs (CDMRP; S.H.S.) Peer Reviewed Medical Research Program (PRMRP; S.H.S), Investigator-Initiated Research Award PR231692P1 (S.H.S.) and the Howard Hughes Medical Institute (HHMI, V.S.T). V.A.L. is supported by the HHMI Hanna Gray fellowship. V.S.T. is a Michael L. Rosenberg Scholar in Medical Research, a CPRIT Scholar (RR150033), a Searle Scholar and an investigator of the HHMI. Use of the Stanford Synchrotron Radiation Lightsource, SLAC National Accelerator Laboratory, is supported by the U.S. Department of Energy, Office of Science, Office of Basic Energy Sciences under Contract No. DE-AC02-76SF00515. The SSRL Structural Molecular Biology Program is supported by the DOE Office of Biological and Environmental Research, and by the National Institutes of Health, National Institute of General Medical Sciences (P30GM133894). The contents of this publication are solely the responsibility of the authors and do not necessarily represent the official views of NIGMS or NIH.

## Author contributions

A.O., B.M., performed experiments; J.S., H.C., and S.S., performed the cryo-ET; A.O. performed the single particle cryo-EM analysis and x-ray crystallography; K.P. performed the bioinformatics, B.M. performed the phage experiments with guidance from K.J.F.; A.O., B.M., V.A.L., and V.S.T., performed protein purification, molecular cloning, and strain construction. D.R.T. collected and processed diffraction data. A.O., B.M., K.P., S.S., and V.S.T. wrote the manuscript with input from all authors.

## Data and materials availability

All materials developed in this study will be made available upon request.

## Supplemental Figures

**Figure S1.**
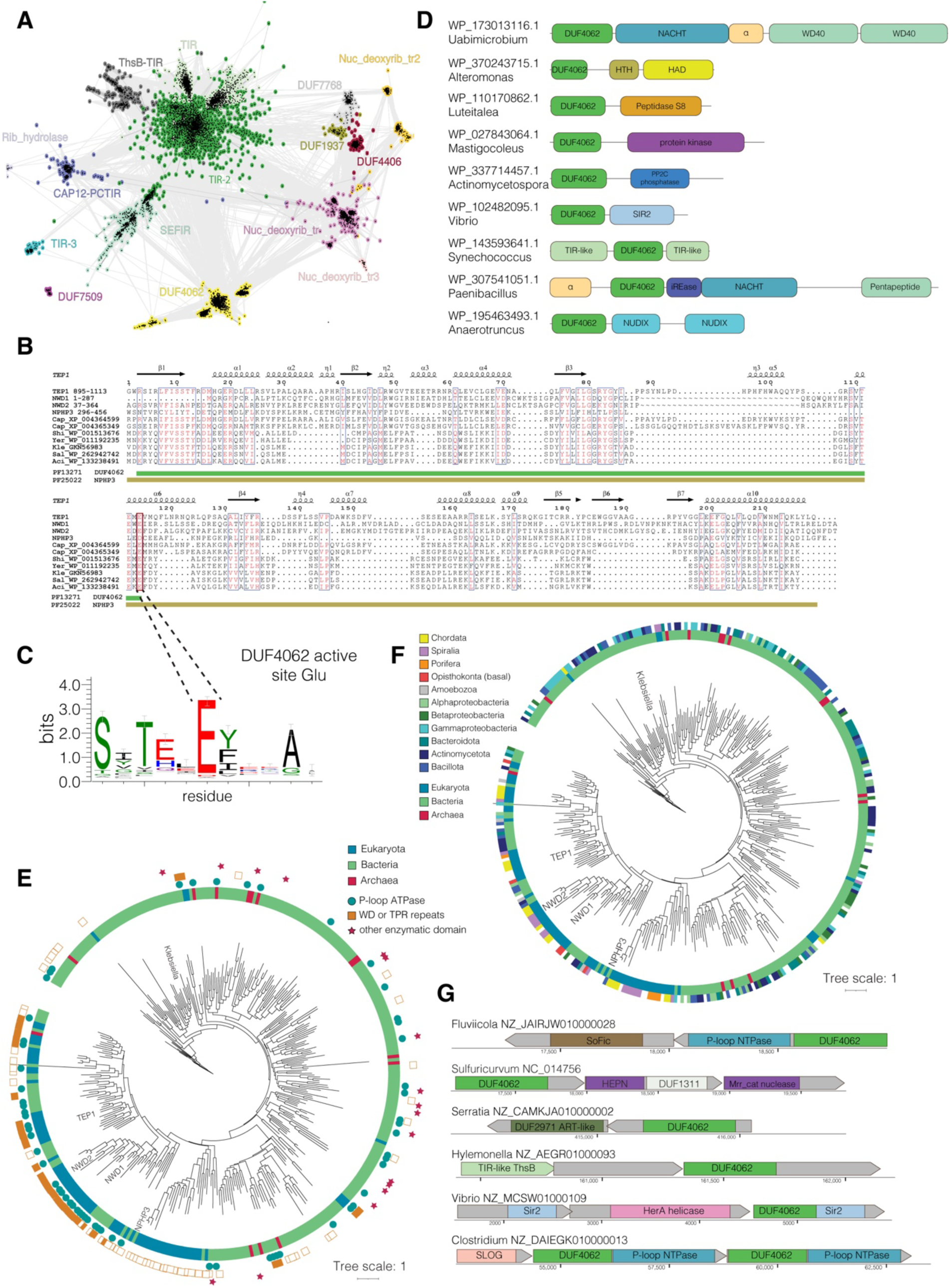
DUF4062 is a TIR-like domain family found in eukaryotes and bacterial anti-phage defense islands. **(A)** CLANS sequence similarity network of representative TIR-like and NDT superfamily proteins (Pfam clans CL0173/STIR and CL0498/Nribosyltransf; BLAST E ≤ 0.01). Each point represents a single protein sequence, and distances reflect relative sequence similarity. As in Figure 1A, with nodes colored by family. **(B)** Multiple sequence alignment of selected DUF4062 domains. Shown are the four human DUF4062 domains, the *Capsaspora* (Cap) DUF4062 domain, and bacterial DUF4062-containing proteins used in the BASEL screen in Figure 1C: *Shigella* (Shi), *Yersinia* (Yer), *Klebsiella* (Kle), *Salmonella* (Sal), and *Acinetobacter* (Aci). NCBI sequence identifiers are shown. A schematic of secondary structure based on the human TEP1 AlphaFold model is shown at the top, whereas the extent of domain annotations in the UniProt database, DUF4062/PF13271 in human TEP1 (green) and NPHP3/PF25022 (olive), is shown at the bottom. **(C)** Sequence logo depicting the conservation of the DUF4062 active site. **(D)** Phylogenetic tree of DUF4062 domains, as in Figure 1C, annotated with auxiliary domains present in the full-length proteins. **(E)** Domain architectures of selected DUF4062 proteins containing auxiliary enzymatic domains. **(F)** Phylogenetic tree of DUF4062 domains, as in Figure 1C, annotated with selected eukaryotic and bacterial taxa. **(G)** Selected operons comprising DUF4062 proteins and associated proteins containing domains implicated in anti-phage defense. See also Figure 1

**Figure S2.**
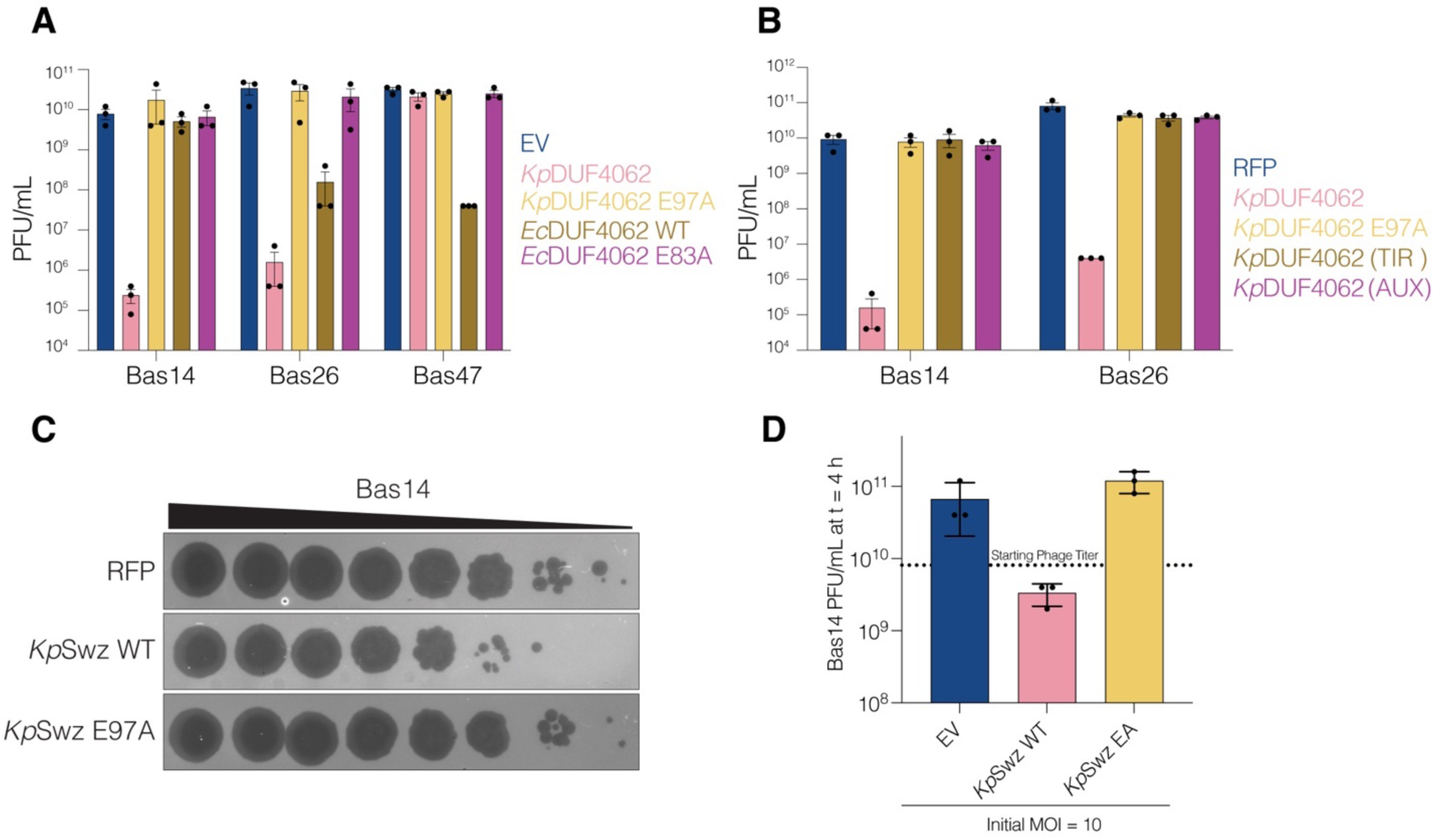
DUF4062 homologs form Swarożyc (Swz), a previously unrecognized antiphage defense system. **(A)** Quantification of phage plaque assays on BW25113 cells expressing RFP or the indicated *Kp*DUF4062^16-C^ (*Kp*DUF4062) and *Ec*DUF4062 constructs. Phages were ten-fold serially diluted. Data are presented as mean ± SEM of n = 3 biological replicates. **(B)** Quantification of phage plaque assays on BW25113 cells expressing RFP or indicated *Kp*DUF4062 constructs. Phages were ten-fold serially diluted. Data are presented as mean ± SEM of n = 3 biological replicates. **(C, D)** Free Bas14 phage collected 200 min post-infection (MOI = 10, Figure 2G). Ten-fold serial dilutions of supernatants were plated on BW25113 lawns expressing RFP **(C)** and quantified in **(D)**. See also Figure 2

**Figure S3.**
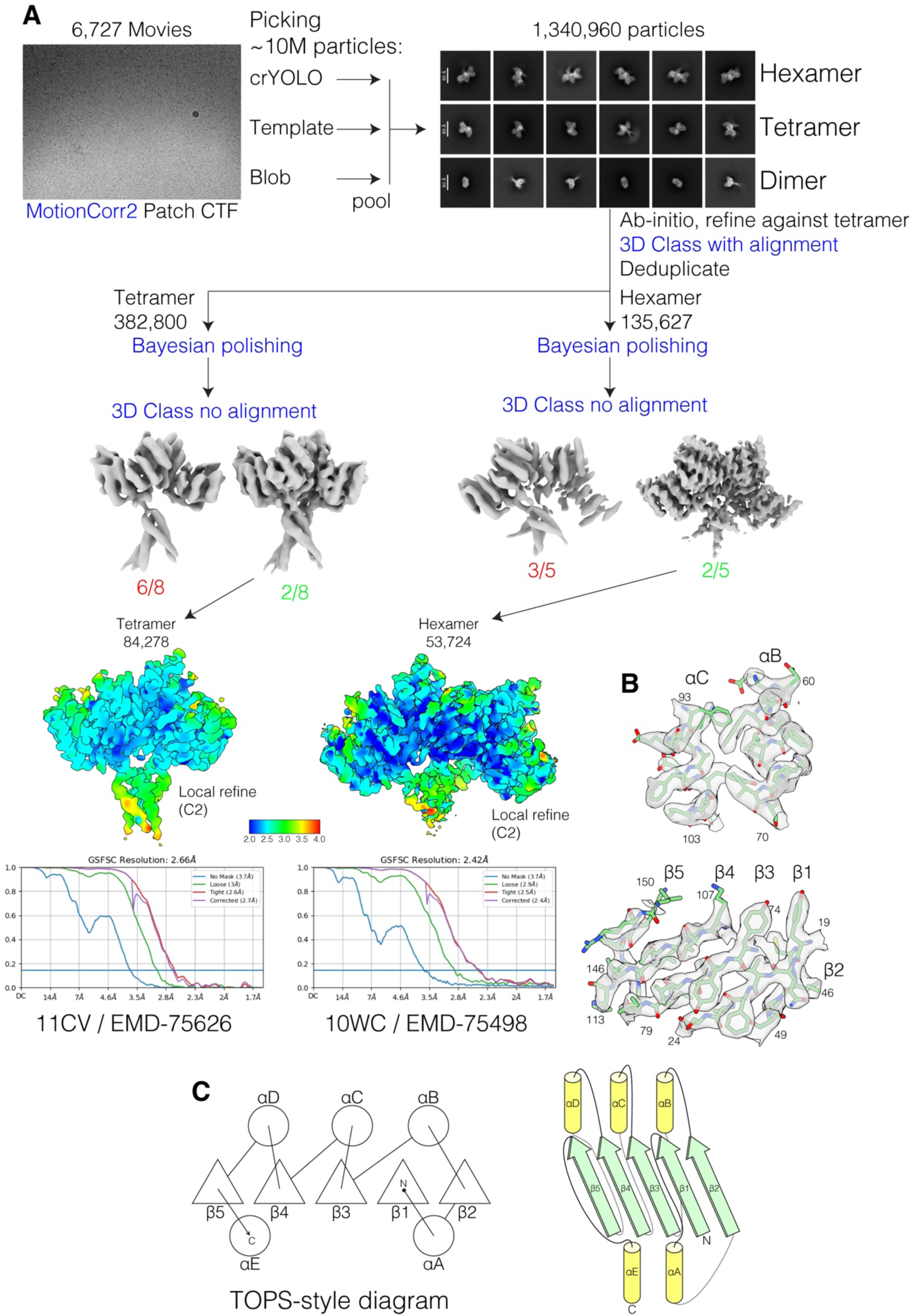
Structural insights into the *Kp*Swz TIR-like domain. **(A)** Simplified Cryo-EM processing schematic of the KpSwz^15-C, E97A^ dataset. **(B)** Density fit of the indicated model portions. **(C)** Topology diagrams depicting the secondary structural elements of *Kp*Swz DUF4062 domain. See also Figure 3

**Figure S4.**
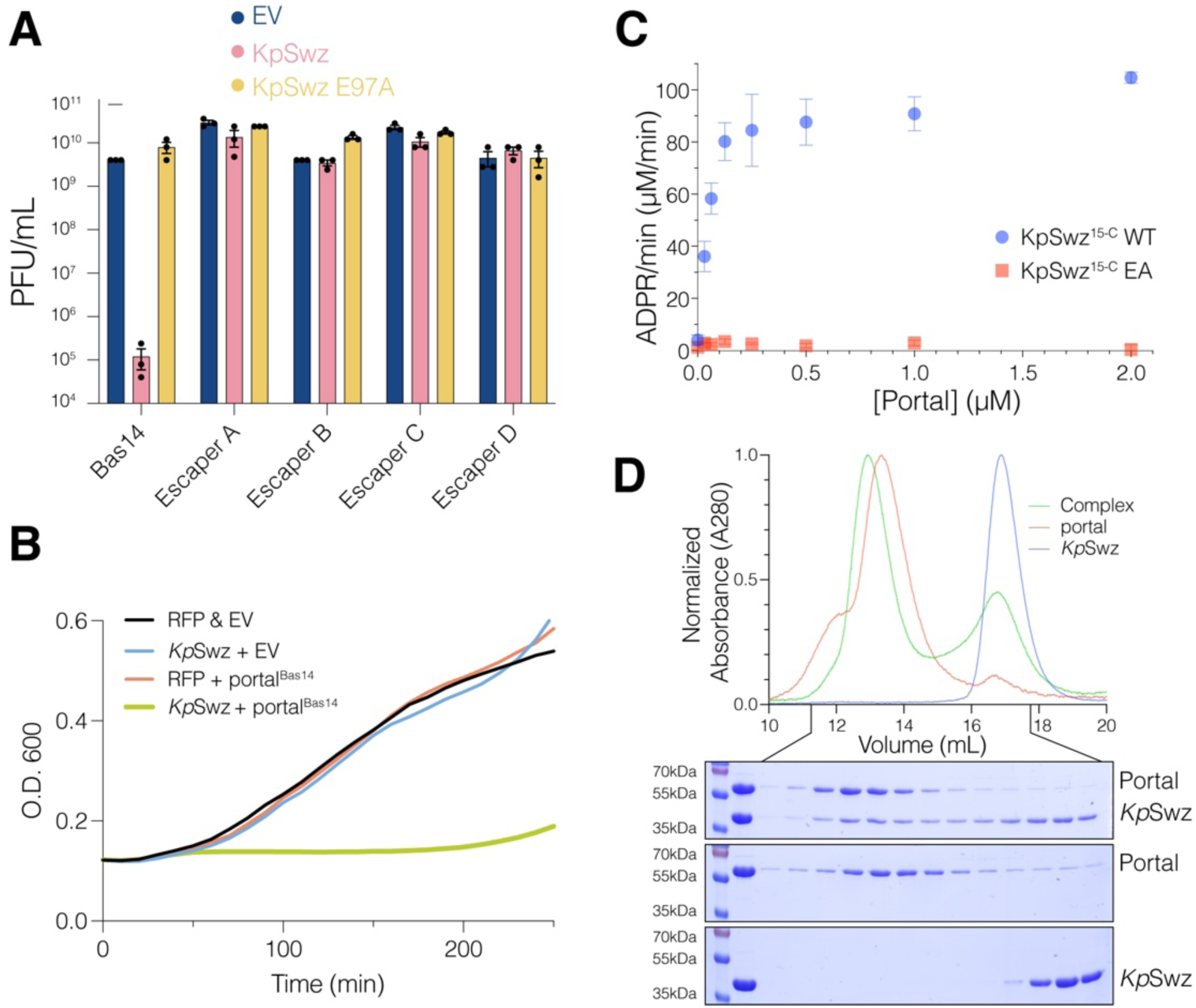
Portal^Bas14^ triggers *Kp*Swz NADase activity and cell death. (A) Quantification of phage plaque assays on BW25113 cells expressing RFP or indicated *Kp*Swz^16-C^ (*Kp*Swz) constructs. Phages were ten-fold serially diluted. Data are presented as mean ± SEM of n = 3 biological replicates. (B) Growth of *E. coli* BW25113 cells expressing the indicated *Kp*Swz and portal proteins. (C) Graph depicting *Kp*Swz^15-C^ NADase activity in the presence of increasing concentrations of portal^Bas14^. Reaction products were separated by HPLC-UV_260_ and quantified by the area under the curve using a standard of ADP-ribose. 0.15 μM of KpSwz^15-C^ was used for 30 minutes at ambient temperature. (D) FPLC-UV_280_ chromatograms of *Kp*Swz^15-C, E97A^ (blue), Portal^Bas14^ (red), and their complex (green). Proteins were separated on a Superose 6 column, and the indicated fractions were analyzed by SDS-PAGE followed by Coomassie staining. ∼150 μg of each protein was used. See also Figure 4

**Figure S5:**
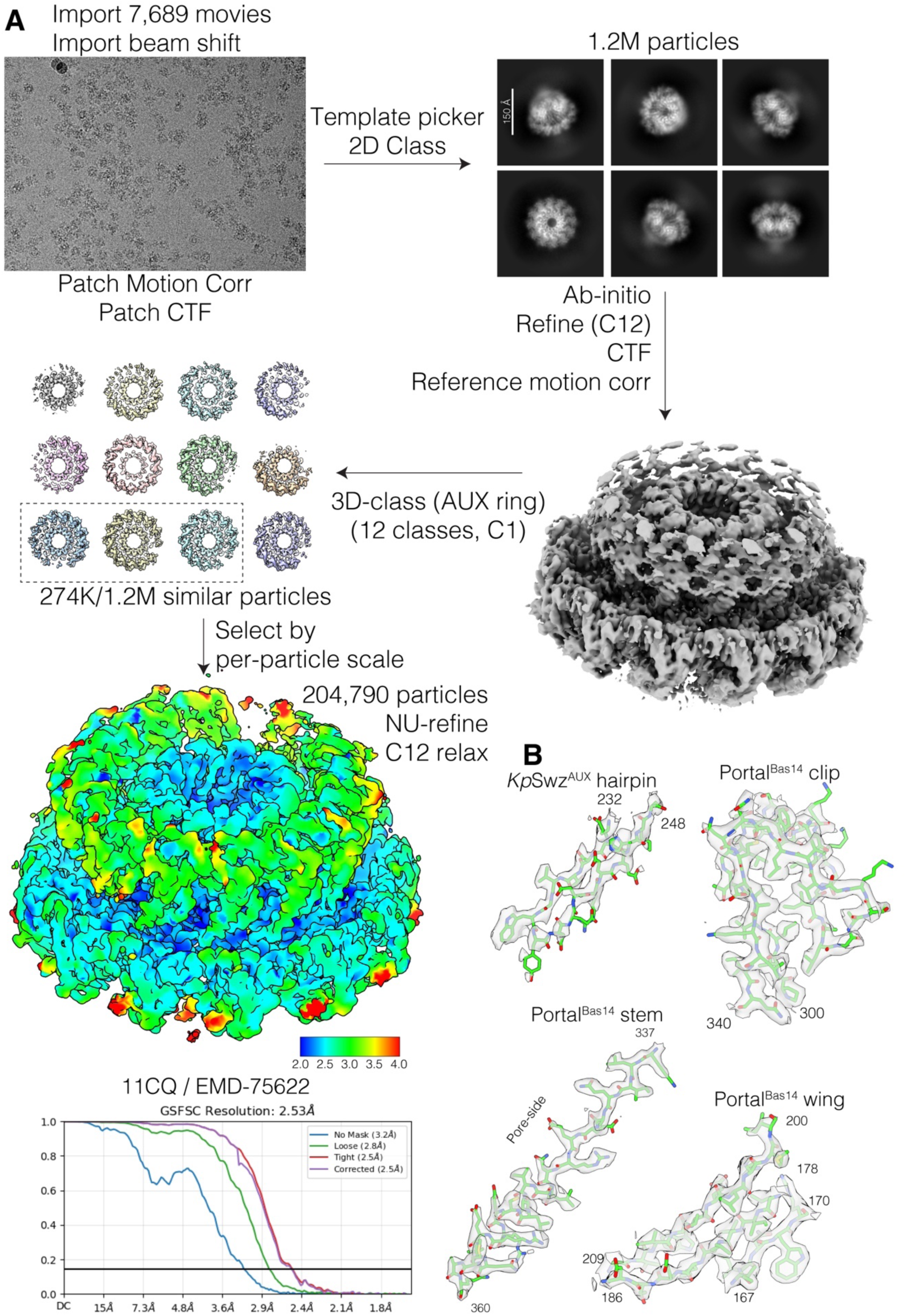
Cryo-EM structure of *Kp*Swz-portal^Bas14^ complex. **(A)** Simplified Cryo-EM processing schematic of the KpSwz^15-C, E97A^ + portal^Bas14^ dataset. **(B)** Density fit of the indicated model portions. See also Figure 4

**Figure S6.**
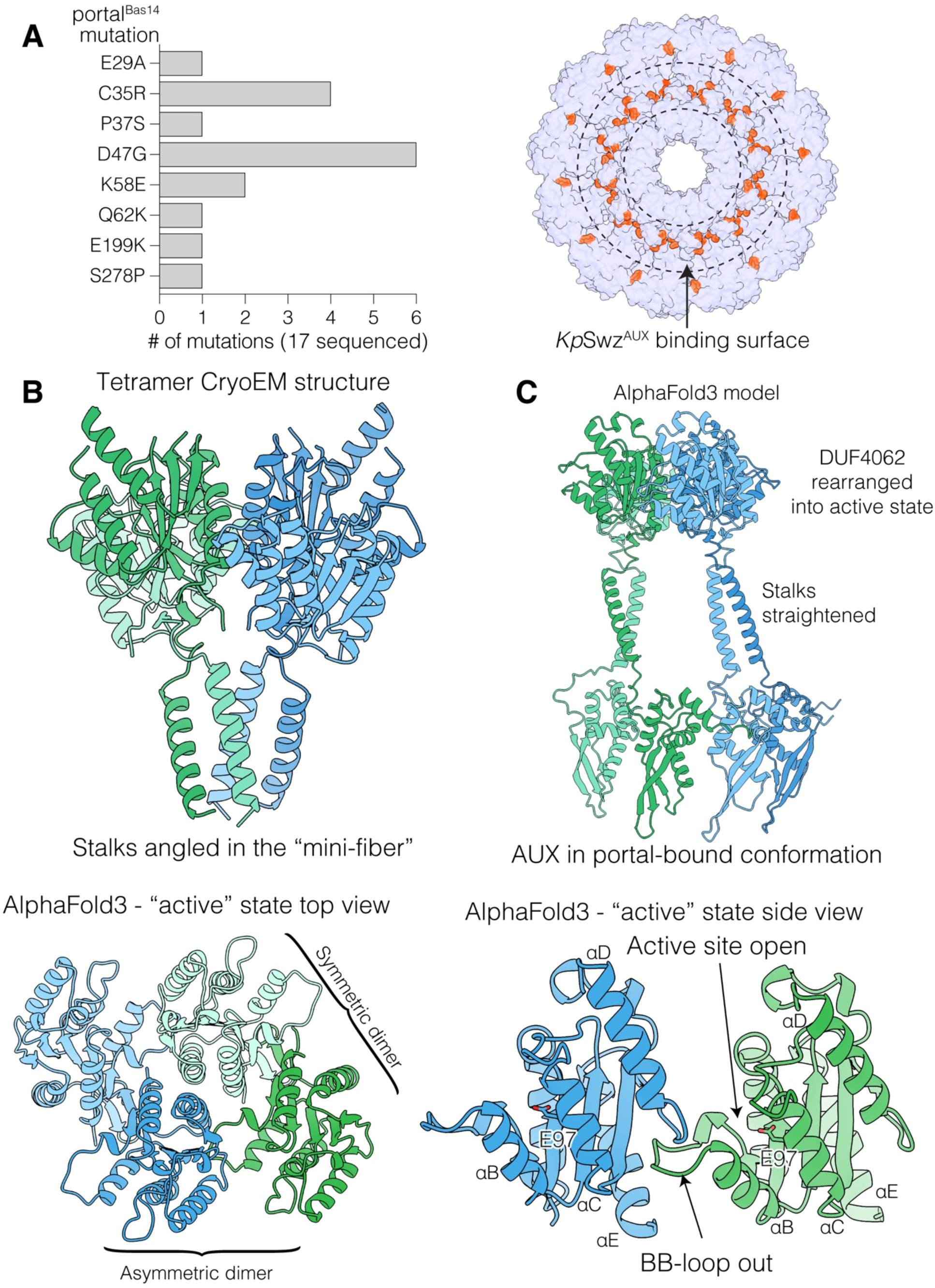
Structural insights into portal^Bas14^ escaper mutations and AlphaFold modeling of the activated *Kp*Swz–portal^Bas14^ complex. **(A)** Chart showing the number of mutations identified for the indicated genotype (left) and surface representation of the *Kp*Swz–portal^Bas14^ complex highlighting the positions of the portal^Bas14^ escaper mutations (right). **(B)** Cartoon representation of the *Kp*Swz tetramer, depicting the intertwined helical stalk (upper) and a zoomed in view of the active site (lower). **(C)** AlphaFold3 modeling of the full-length *Kp*Swz–portal^Bas14^ complex suggests an activation mechanism in which portal^Bas14^ engagement with *Kp*Swz^AUX^ pulls on the helical stalks, aligns them (upper), and ultimately repositions the TIR domains into an active conformation (lower). See also Figure 4

**Figure S7.**
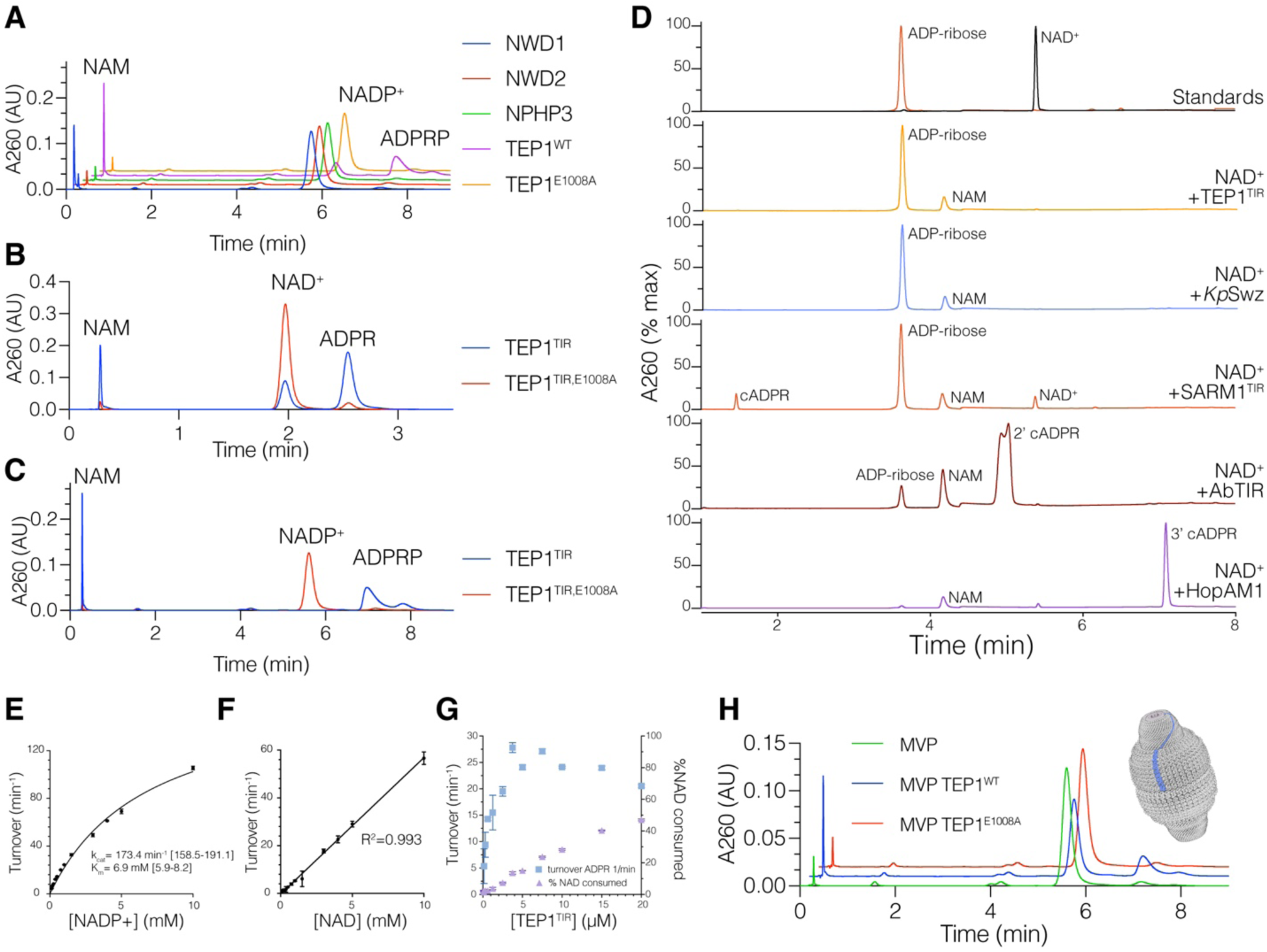
Characterization of TEP1^TIR^ activity. **(A)** HPLC traces depicting the reaction products generated after incubation of NADP^+^ with TEP1, NWD1, NWD2 and NPHP3. Full-length protein preparations were used at 0.5 mg/mL. (**B, C**) HPLC traces depicting the reaction products generated after incubation of NAD^+^ **(B)** or NADP^+^ **(C)** with TEP1^TIR^ or the E1008A mutant. (**D**) HPLC traces depicting the reaction products generated after incubation of NAD^+^ for 1 h at 37 °C with 0.3 mg/mL of each TEP1^TIR^, *Kp*Swz^15-C^ (*Kp*Swz), SARM1^TIR^, AbTIR, or HopAM1 (cyclic ADP-ribose; cADPR. (**E, F**) Kinetic analysis of NADP^+^ **(E)** or NAD^+^ **(F)** hydrolysis by 1 μM TEP1^TIR^, depicting the concentration dependence of NAD(P)^+^ on reaction rate. The inset shows the corresponding apparent K_m_ and k_cat_ values. Reaction products were analyzed and quantified as described in **Figure S4C**. **(G)** Graph showing the specific activity of TEP1^TIR^ at increasing concentrations of TEP1^TIR^. Percentage of consumed NAD⁺ is plotted on the right. Reaction products were analyzed as in **Figure S4C.** Reactions were conducted for 10 minutes with 10 mM NAD^+^. **(H)** HPLC traces depicting the reaction products generated after incubation of NADP^+^ with the RNA vault (0.75 mg/mL) containing either TEP1 (red) or the E1008A mutant (green). See also Figure 5

**Figure S8.**
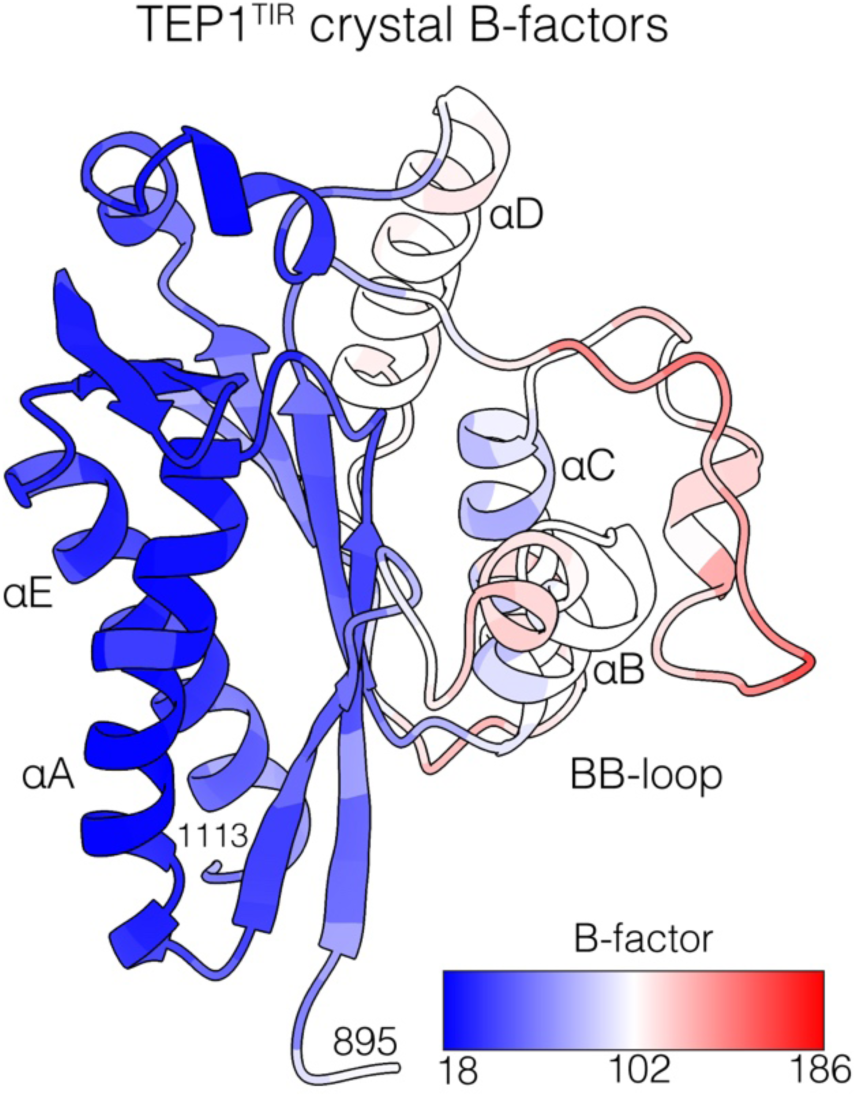
Structure of the TEP1^TIR^ domain. Crystal structure of the TEP1^TIR^ (895-1113) is shown as a cartoon representation and colored by B-factors at Cα atoms. Note the quality degradation towards the flexible regions of the protein around the active site. See also Figure 5

**Figure S9.**
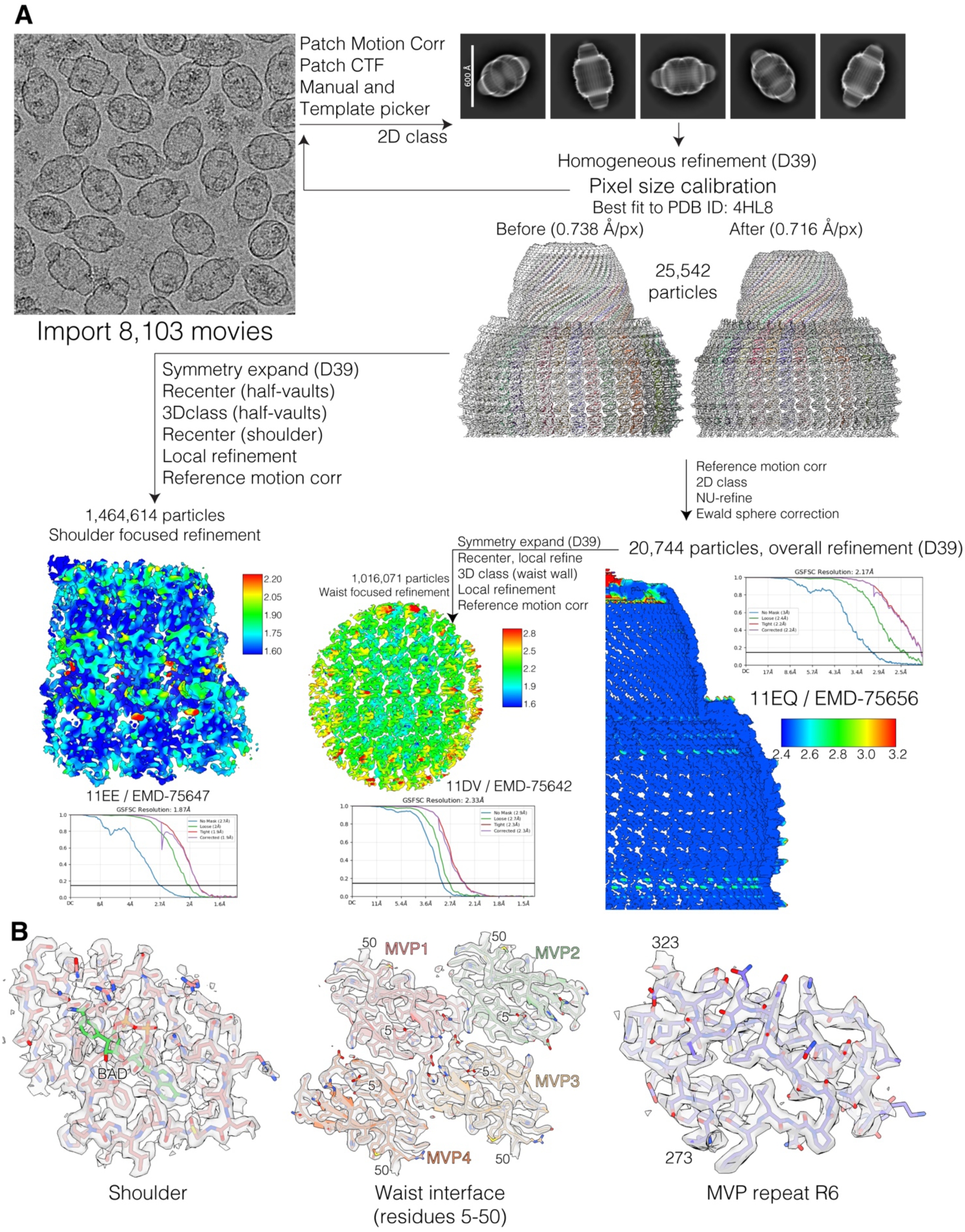
Structure of the RNA vault with TEP1, BAD and AMP-PNP. **(A)** Simplified data processing schematic for the dataset. **(B)** Example density fit for arbitrarily chosen protein regions. See also Figure 6

**Figure S10.**
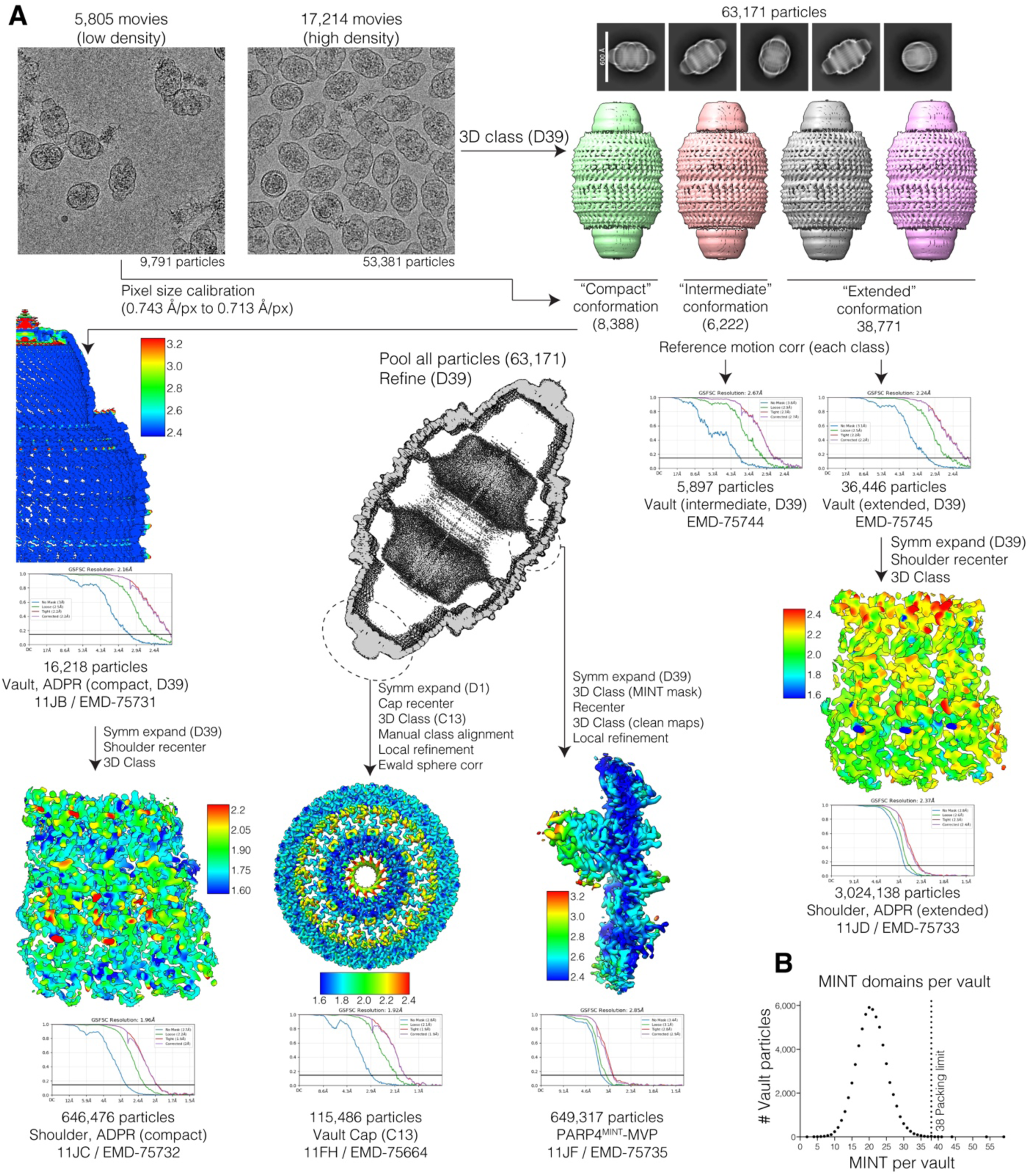
Structure of the RNA vault with PARP4, TEP1 and NADP^+^. **(A)** Simplified data processing schematic for the dataset. **(B)** The MINT domains in each vault shell from the initial MINT-focused 3D classification were counted and plotted as a histogram. See also Figure 6

**Figure S11.**
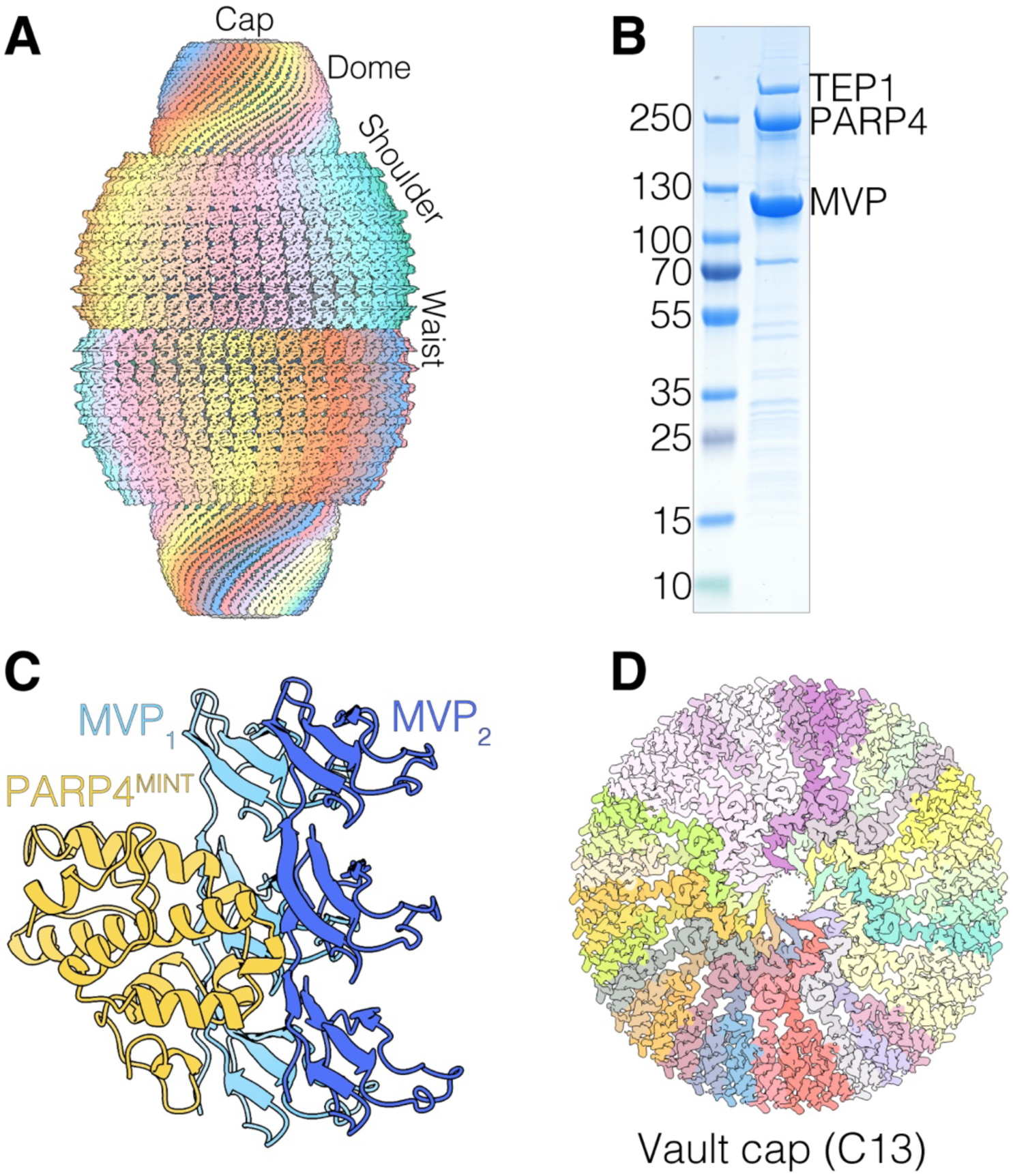
Cryo-EM analysis of reconstituted RNA vaults. **(A)** Orthographic projection of the human RNA vault shell. The unsharpened Coulombic map of a D39-symmetric vault is shown, colored by MVP monomer. **(B)** Coomassie stained SDS-PAGE analysis of the purified reconstituted RNA vault. Bands corresponding to MVP, TEP1, and PARP4 are indicated. **(C)** Cartoon depiction of the PARP4^MINT^ domain (yellow) bound between two MVP monomers (blue/light blue). **(D)** Coulombic map of the human RNA vault cap, refined with C13 symmetry. See also Figure 6

**Figure S12.**
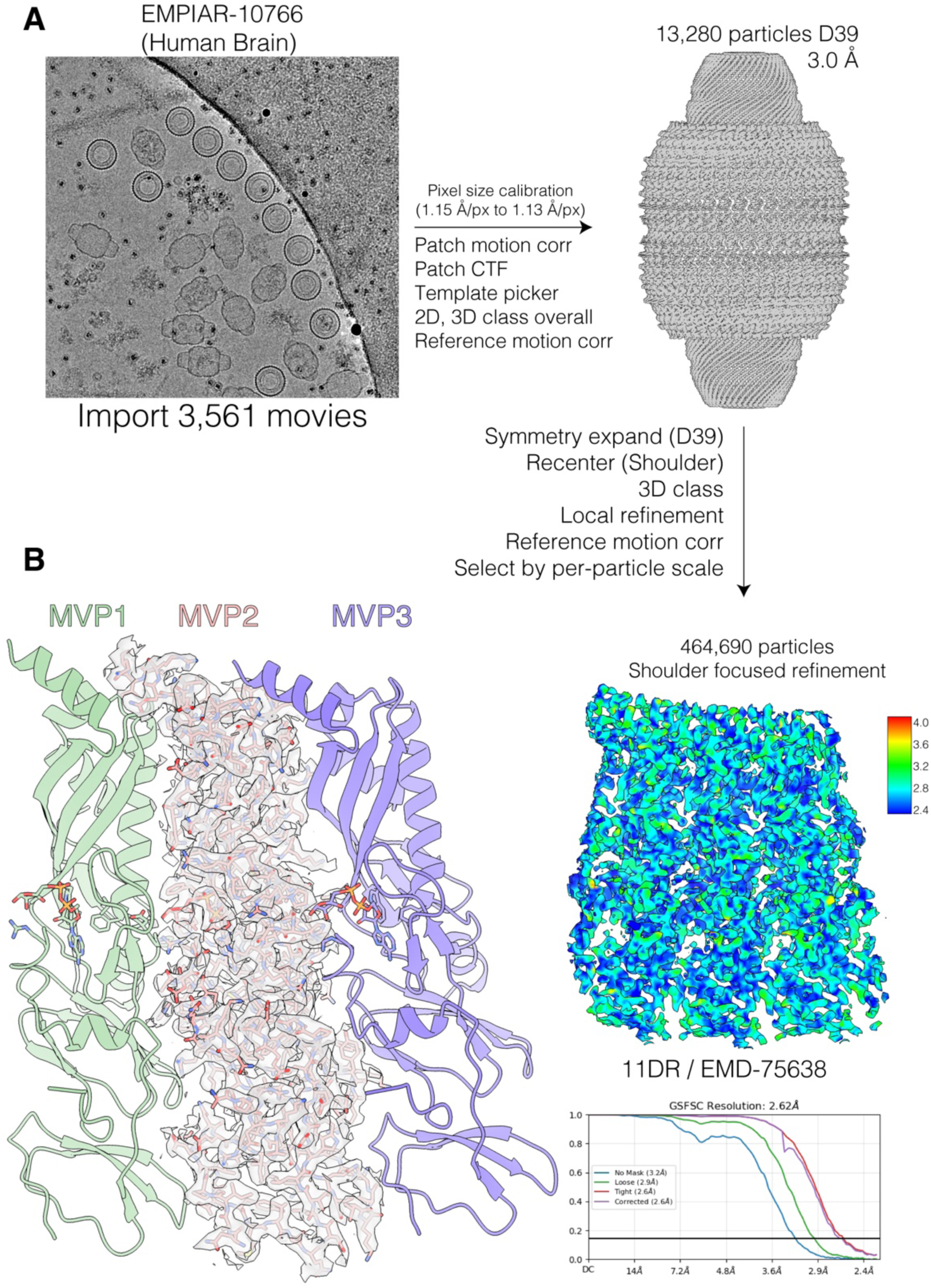
Reprocessing of vaults from human brain reveals a bound ADP-ribose. **(A)** Simplified Cryo-EM processing schematic of the EMPIAR-10766 dataset. **(B)** Density fit of the model. See also Figure 6

**Figure S13.**
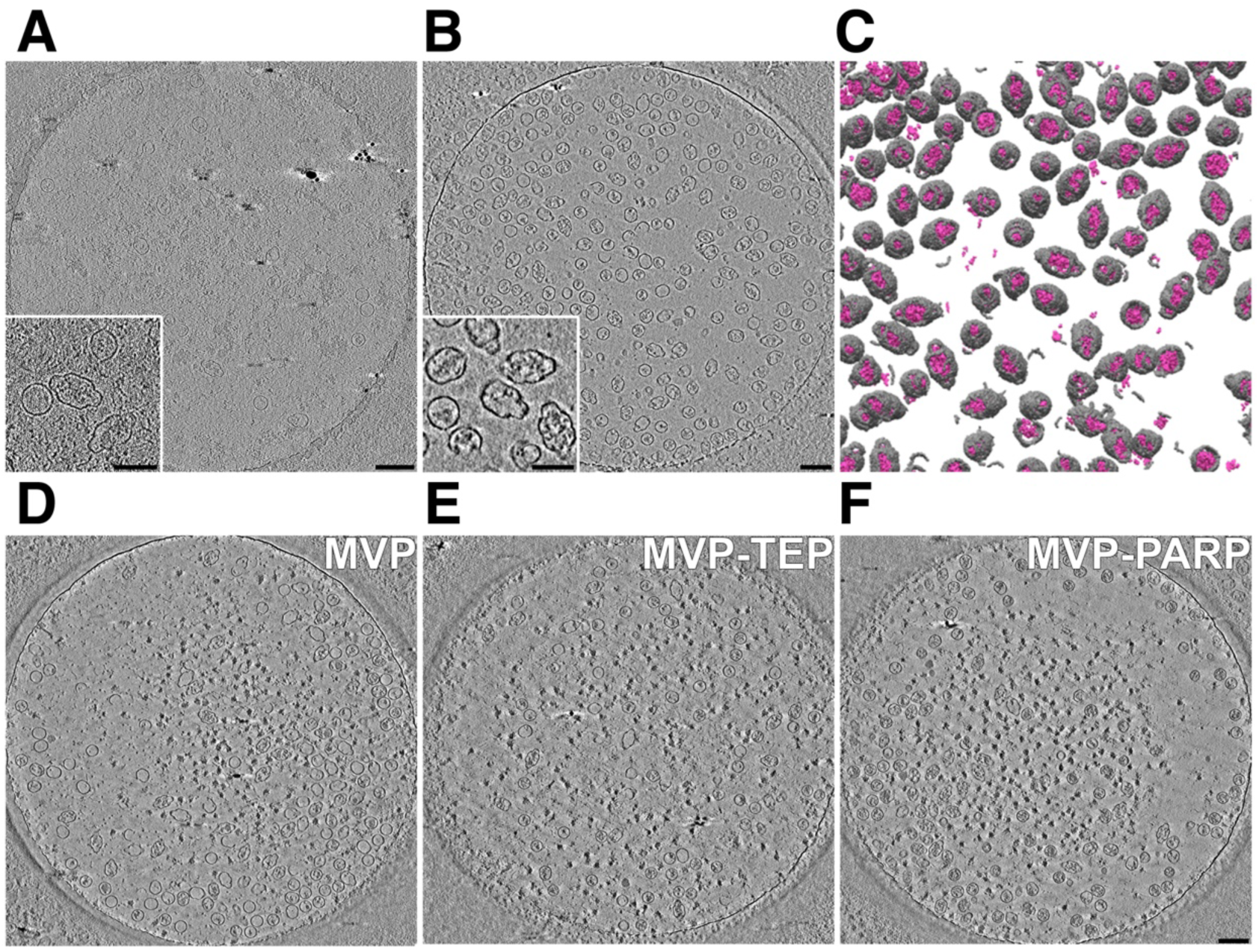
Cryo-ET analysis of reconstituted RNA vaults. **(A–B)** Central slices from reconstituted vault tomograms acquired using either a conventional dose-symmetric tilt scheme **(A)** or the high-contrast tilt scheme used in this study **(B)**. Insets show magnified views of representative particles from each tomogram. The high-contrast acquisition enhances visualization of lumenal features relative to conventional dose-symmetric collection. **(C)** Isosurface renderings of reconstituted vault particles segmented from a high-contrast tomogram. Segmentation was performed in Dragonfly using independently trained models for the vault exterior and interior. Higher-magnification views of representative particles from this dataset are shown in Figure 6B. **(D–F)** Central slices (5 nm thick) from high-contrast tomograms of reconstituted MVP-only **(D)**, MVP–TEP1 **(E)**, and MVP–PARP4 **(F)** vault particles, illustrating composition-dependent differences in lumenal density under identical imaging conditions. Scale bars represent 10 nm. See also Figure 6

## References

1. Abramson, J., Adler, J., Dunger, J., Evans, R., Green, T., Pritzel, A., Ronneberger, O., Willmore, L., Ballard, A.J., Bambrick, J., et al. (2024). Accurate structure prediction of biomolecular interactions with AlphaFold 3. Nature. 10.1038/s41586-024-07487-w.

2. Jumper, J., Evans, R., Pritzel, A., Green, T., Figurnov, M., Ronneberger, O., Tunyasuvunakool, K., Bates, R., Žídek, A., Potapenko, A., et al. (2021). Highly accurate protein structure prediction with AlphaFold. Nature 596, 583–589. 10.1038/s41586-021-03819-2.

3. Sreelatha, A., Yee, S.S., Lopez, V.A., Park, B.C., Kinch, L.N., Pilch, S., Servage, K.A., Zhang, J., Jiou, J., Karasiewicz-Urbanska, M., et al. (2018). Protein AMPylation by an Evolutionarily Conserved Pseudokinase. Cell 175, 809–821 e819. 10.1016/j.cell.2018.08.046.

4. Black, M.H., Osinski, A., Gradowski, M., Servage, K.A., Pawlowski, K., Tomchick, D.R., and Tagliabracci, V.S. (2019). Bacterial pseudokinase catalyzes protein polyglutamylation to inhibit the SidE-family ubiquitin ligases. Science 364, 787–792. 10.1126/science.aaw7446.

5. Park, G.J., Osinski, A., Hernandez, G., Eitson, J.L., Majumdar, A., Tonelli, M., Henzler-Wildman, K., Pawlowski, K., Chen, Z., Li, Y., et al. (2022). The mechanism of RNA capping by SARS-CoV-2. Nature 609, 793–800. 10.1038/s41586-022-05185-z.

6. Osinski, A., Black, M.H., Pawlowski, K., Chen, Z., Li, Y., and Tagliabracci, V.S. (2021). Structural and mechanistic basis for protein glutamylation by the kinase fold. Mol Cell 81, 4527–4539 e4528. 10.1016/j.molcel.2021.08.007.

7. Essuman, K., Milbrandt, J., Dangl, J.L., and Nishimura, M.T. (2022). Shared TIR enzymatic functions regulate cell death and immunity across the tree of life. Science 377, eabo0001. 10.1126/science.abo0001.

8. Wang, M., Ji, Q., Liu, P., and Liu, Y. (2023). NAD(+) depletion and defense in bacteria. Trends Microbiol 31, 435–438. 10.1016/j.tim.2022.06.002.

9. Essuman, K., Summers, D.W., Sasaki, Y., Mao, X., DiAntonio, A., and Milbrandt, J. (2017). The SARM1 Toll/Interleukin-1 Receptor Domain Possesses Intrinsic NAD(+) Cleavage Activity that Promotes Pathological Axonal Degeneration. Neuron 93, 1334–1343 e1335. 10.1016/j.neuron.2017.02.022.

10. Wan, L., Essuman, K., Anderson, R.G., Sasaki, Y., Monteiro, F., Chung, E.H., Osborne Nishimura, E., DiAntonio, A., Milbrandt, J., Dangl, J.L., and Nishimura, M.T. (2019). TIR domains of plant immune receptors are NAD(+)-cleaving enzymes that promote cell death. Science 365, 799–803. 10.1126/science.aax1771.

11. Horsefield, S., Burdett, H., Zhang, X., Manik, M.K., Shi, Y., Chen, J., Qi, T., Gilley, J., Lai, J.S., Rank, M.X., et al. (2019). NAD(+) cleavage activity by animal and plant TIR domains in cell death pathways. Science 365, 793–799. 10.1126/science.aax1911.

12. Hogrel, G., Guild, A., Graham, S., Rickman, H., Gruschow, S., Bertrand, Q., Spagnolo, L., and White, M.F. (2022). Cyclic nucleotide-induced helical structure activates a TIR immune effector. Nature 608, 808–812. 10.1038/s41586-022-05070-9.

13. Koopal, B., Potocnik, A., Mutte, S.K., Aparicio-Maldonado, C., Lindhoud, S., Vervoort, J.J.M., Brouns, S.J.J., and Swarts, D.C. (2022). Short prokaryotic Argonaute systems trigger cell death upon detection of invading DNA. Cell 185, 1471–1486 e1419. 10.1016/j.cell.2022.03.012.

14. Baca, C.F., Majumder, P., Hickling, J.H., Patel, D.J., and Marraffini, L.A. (2025). Cat1 forms filament networks to degrade NAD(+) during the type III CRISPR-Cas antiviral response. Science 388, eadv9045. 10.1126/science.adv9045.

15. Kibby, E.M., Conte, A.N., Burroughs, A.M., Nagy, T.A., Vargas, J.A., Whalen, L.A., Aravind, L., and Whiteley, A.T. (2023). Bacterial NLR-related proteins protect against phage. Cell 186, 2410–2424 e2418. 10.1016/j.cell.2023.04.015.

16. Morehouse, B.R., Govande, A.A., Millman, A., Keszei, A.F.A., Lowey, B., Ofir, G., Shao, S., Sorek, R., and Kranzusch, P.J. (2020). STING cyclic dinucleotide sensing originated in bacteria. Nature 586, 429–433. 10.1038/s41586-020-2719-5.

17. Zaremba, M., Dakineviciene, D., Golovinas, E., Zagorskaite, E., Stankunas, E., Lopatina, A., Sorek, R., Manakova, E., Ruksenaite, A., Silanskas, A., et al. (2022). Short prokaryotic Argonautes provide defence against incoming mobile genetic elements through NAD(+) depletion. Nat Microbiol 7, 1857–1869. 10.1038/s41564-022-01239-0.

18. Tal, N., Morehouse, B.R., Millman, A., Stokar-Avihail, A., Avraham, C., Fedorenko, T., Yirmiya, E., Herbst, E., Brandis, A., Mehlman, T., et al. (2021). Cyclic CMP and cyclic UMP mediate bacterial immunity against phages. Cell 184, 5728–5739 e5716. 10.1016/j.cell.2021.09.031.

19. Essuman, K., Summers, D.W., Sasaki, Y., Mao, X., Yim, A.K.Y., DiAntonio, A., and Milbrandt, J. (2018). TIR Domain Proteins Are an Ancient Family of NAD(+)-Consuming Enzymes. Curr Biol 28, 421–430 e424. 10.1016/j.cub.2017.12.024.

20. Manik, M.K., Shi, Y., Li, S., Zaydman, M.A., Damaraju, N., Eastman, S., Smith, T.G., Gu, W., Masic, V., Mosaiab, T., et al. (2022). Cyclic ADP ribose isomers: Production, chemical structures, and immune signaling. Science 377, eadc8969. 10.1126/science.adc8969.

21. Ofir, G., Herbst, E., Baroz, M., Cohen, D., Millman, A., Doron, S., Tal, N., Malheiro, D.B.A., Malitsky, S., Amitai, G., and Sorek, R. (2021). Antiviral activity of bacterial TIR domains via immune signalling molecules. Nature 600, 116–120. 10.1038/s41586-021-04098-7.

22. Leavitt, A., Yirmiya, E., Amitai, G., Lu, A., Garb, J., Herbst, E., Morehouse, B.R., Hobbs, S.J., Antine, S.P., Sun, Z.J., et al. (2022). Viruses inhibit TIR gcADPR signalling to overcome bacterial defence. Nature 611, 326–331. 10.1038/s41586-022-05375-9.

23. Sabonis, D., Avraham, C., Chang, R.B., Lu, A., Herbst, E., Silanskas, A., Vilutis, D., Leavitt, A., Yirmiya, E., Toyoda, H.C., et al. (2025). TIR domains produce histidine-ADPR as an immune signal in bacteria. Nature 642, 467–473. 10.1038/s41586-025-08930-2.

24. Rousset, F., Osterman, I., Scherf, T., Falkovich, A.H., Leavitt, A., Amitai, G., Shir, S., Malitsky, S., Itkin, M., Savidor, A., and Sorek, R. (2025). TIR signaling activates caspase-like immunity in bacteria. Science 387, 510–516. 10.1126/science.adu2262.

25. Huang, S., Jia, A., Song, W., Hessler, G., Meng, Y., Sun, Y., Xu, L., Laessle, H., Jirschitzka, J., Ma, S., et al. (2022). Identification and receptor mechanism of TIR-catalyzed small molecules in plant immunity. Science 377, eabq3297. 10.1126/science.abq3297.

26. Jia, A., Huang, S., Song, W., Wang, J., Meng, Y., Sun, Y., Xu, L., Laessle, H., Jirschitzka, J., Hou, J., et al. (2022). TIR-catalyzed ADP-ribosylation reactions produce signaling molecules for plant immunity. Science 377, eabq8180. 10.1126/science.abq8180.

27. Bayless, A.M., Chen, S., Ogden, S.C., Xu, X., Sidda, J.D., Manik, M.K., Li, S., Kobe, B., Ve, T., Song, L., et al. (2023). Plant and prokaryotic TIR domains generate distinct cyclic ADPR NADase products. Sci Adv 9, eade8487. 10.1126/sciadv.ade8487.

28. Wu, Y., Xu, W., Zhao, G., Lei, Z., Li, K., Liu, J., Huang, S., Wang, J., Zhong, X., Yin, X., et al. (2024). A canonical protein complex controls immune homeostasis and multipathogen resistance. Science 386, 1405–1412. 10.1126/science.adr2138.

29. Martin, R., Qi, T., Zhang, H., Liu, F., King, M., Toth, C., Nogales, E., and Staskawicz, B.J. (2020). Structure of the activated ROQ1 resistosome directly recognizing the pathogen effector XopQ. Science 370. 10.1126/science.abd9993.

30. Ma, S., Lapin, D., Liu, L., Sun, Y., Song, W., Zhang, X., Logemann, E., Yu, D., Wang, J., Jirschitzka, J., et al. (2020). Direct pathogen-induced assembly of an NLR immune receptor complex to form a holoenzyme. Science 370. 10.1126/science.abe3069.

31. Travis, J. (2024). The vault guy. Science 384, 1058–1062. 10.1126/science.adq8600.

32. Black, M.H., Gradowski, M., Pawlowski, K., and Tagliabracci, V.S. (2022). Methods for discovering catalytic activities for pseudokinases. Methods Enzymol 667, 575–610. 10.1016/bs.mie.2022.03.047.

33. Maffei, E., Shaidullina, A., Burkolter, M., Heyer, Y., Estermann, F., Druelle, V., Sauer, P., Willi, L., Michaelis, S., Hilbi, H., et al. (2021). Systematic exploration of Escherichia coli phage-host interactions with the BASEL phage collection. PLoS Biol 19, e3001424. 10.1371/journal.pbio.3001424.

34. Lopatina, A., Tal, N., and Sorek, R. (2020). Abortive Infection: Bacterial Suicide as an Antiviral Immune Strategy. Annu Rev Virol 7, 371–384. 10.1146/annurev-virology-011620-040628.

35. Hochhauser, D., and Sorek, R. (2025). Manipulation of the nucleotide pool in human, bacterial and plant immunity. Nat Rev Immunol. 10.1038/s41577-025-01206-w.

36. Nimma, S., Gu, W., Maruta, N., Li, Y., Pan, M., Saikot, F.K., Lim, B.Y.J., McGuinness, H.Y., Zaoti, Z.F., Li, S., et al. (2021). Structural Evolution of TIR-Domain Signalosomes. Front Immunol 12, 784484. 10.3389/fimmu.2021.784484.

37. Stokar-Avihail, A., Fedorenko, T., Hor, J., Garb, J., Leavitt, A., Millman, A., Shulman, G., Wojtania, N., Melamed, S., Amitai, G., and Sorek, R. (2023). Discovery of phage determinants that confer sensitivity to bacterial immune systems. Cell 186, 1863–1876 e1816. 10.1016/j.cell.2023.02.029.

38. Dedeo, C.L., Cingolani, G., and Teschke, C.M. (2019). Portal Protein: The Orchestrator of Capsid Assembly for the dsDNA Tailed Bacteriophages and Herpesviruses. Annu Rev Virol 6, 141–160. 10.1146/annurev-virology-092818-015819.

39. Wang, S., Kuang, S., Song, H., Sun, E., Li, M., Liu, Y., Xia, Z., Zhang, X., Wang, X., Han, J., et al. (2024). The role of TIR domain-containing proteins in bacterial defense against phages. Nat Commun 15, 7384. 10.1038/s41467-024-51738-3.

40. Loyo, C.L., and Grossman, A.D. (2025). A phage-encoded counter-defense inhibits an NAD-degrading anti-phage defense system. PLoS Genet 21, e1011551. 10.1371/journal.pgen.1011551.

41. Zhang, T., Lyu, Y., Beck, C.R., Iqbal, N., Barbosa, R., Ghanbarpour, A., and Laub, M.T. (2026). Bacterial immune activation via supramolecular assembly with phage triggers. Nature. 10.1038/s41586-025-10060-8.

42. Yang, Y., Liu, Y., Ma, X., Zhao, X., Cao, J., Liu, Y., Li, S., Wu, J., Gao, Y., Chen, L., et al. (2025). Structural insights into distinct filamentation states reveal a regulatory mechanism for bacterial STING activation. mBio 16, e00388–00325. doi:10.1128/mbio.00388-25.

43. Li, P., Nijhawan, D., Budihardjo, I., Srinivasula, S.M., Ahmad, M., Alnemri, E.S., and Wang, X. (1997). Cytochrome c and dATP-dependent formation of Apaf-1/caspase-9 complex initiates an apoptotic protease cascade. Cell 91, 479–489. 10.1016/s0092-8674(00)80434-1.

44. Harrington, L., McPhail, T., Mar, V., Zhou, W., Oulton, R., Bass, M.B., Arruda, I., and Robinson, M.O. (1997). A mammalian telomerase-associated protein. Science 275, 973–977. 10.1126/science.275.5302.973.

45. Liu, Y., Snow, B.E., Hande, M.P., Baerlocher, G., Kickhoefer, V.A., Yeung, D., Wakeham, A., Itie, A., Siderovski, D.P., Lansdorp, P.M., et al. (2000). Telomerase-associated protein TEP1 is not essential for telomerase activity or telomere length maintenance in vivo. Mol Cell Biol 20, 8178–8184. 10.1128/MCB.20.21.8178-8184.2000.

46. Kedersha, N.L., and Rome, L.H. (1986). Preparative agarose gel electrophoresis for the purification of small organelles and particles. Anal Biochem 156, 161–170. 10.1016/0003-2697(86)90168-5.

47. Kedersha, N.L., and Rome, L.H. (1986). Isolation and characterization of a novel ribonucleoprotein particle: large structures contain a single species of small RNA. J Cell Biol 103, 699–709. 10.1083/jcb.103.3.699.

48. Tanaka, H., Kato, K., Yamashita, E., Sumizawa, T., Zhou, Y., Yao, M., Iwasaki, K., Yoshimura, M., and Tsukihara, T. (2009). The structure of rat liver vault at 3.5 angstrom resolution. Science 323, 384–388. 10.1126/science.1164975.

49. Kickhoefer, V.A., Stephen, A.G., Harrington, L., Robinson, M.O., and Rome, L.H. (1999). Vaults and telomerase share a common subunit, TEP1. J Biol Chem *274*, 32712-32717. 10.1074/jbc.274.46.32712.

50. Kickhoefer, V.A., Siva, A.C., Kedersha, N.L., Inman, E.M., Ruland, C., Streuli, M., and Rome, L.H. (1999). The 193-kD vault protein, VPARP, is a novel poly(ADP-ribose) polymerase. J Cell Biol 146, 917–928. 10.1083/jcb.146.5.917.

51. Lodwick, J.E., Shen, R., Erramilli, S., Xie, Y., Roganowicz, K., Ritchey, S., Kossiakoff, A.A., and Zhao, M. (2025). Structural insights into the roles of PARP4 and NAD(+) binding in the human vault cage. Nat Commun 16, 6724. 10.1038/s41467-025-61981-x.

52. Lovestam, S., and Scheres, S.H.W. (2025). Cryo-EM structure of the vault from human brain reveals symmetry mismatch at its caps. Structure 33, 1643–1648 e1641. 10.1016/j.str.2025.07.014.

53. Li, H., Vallese, F., and Clarke, O.B. (2025). The vault particle is enclosed by a C13-symmetric cap with a positively charged exterior. bioRxiv, 2025.2006.2006.658390. 10.1101/2025.06.06.658390.

54. Shi, Y., Zhang, W., Yang, Y., Murzin, A.G., Falcon, B., Kotecha, A., van Beers, M., Tarutani, A., Kametani, F., Garringer, H.J., et al. (2021). Structure-based classification of tauopathies. Nature 598, 359–363. 10.1038/s41586-021-03911-7.

55. Kong, L.B., Siva, A.C., Kickhoefer, V.A., Rome, L.H., and Stewart, P.L. (2000). RNA location and modeling of a WD40 repeat domain within the vault. RNA 6, 890–900. 10.1017/s1355838200000157.

56. Woodward, C.L., Mendonca, L.M., and Jensen, G.J. (2015). Direct visualization of vaults within intact cells by electron cryo-tomography. Cell Mol Life Sci 72, 3401–3409. 10.1007/s00018-015-1898-y.

57. Whiteley, A.T., Eaglesham, J.B., de Oliveira Mann, C.C., Morehouse, B.R., Lowey, B., Nieminen, E.A., Danilchanka, O., King, D.S., Lee, A.S.Y., Mekalanos, J.J., and Kranzusch, P.J. (2019). Bacterial cGAS-like enzymes synthesize diverse nucleotide signals. Nature 567, 194–199. 10.1038/s41586-019-0953-5.

58. Johnson, A.G., Wein, T., Mayer, M.L., Duncan-Lowey, B., Yirmiya, E., Oppenheimer-Shaanan, Y., Amitai, G., Sorek, R., and Kranzusch, P.J. (2022). Bacterial gasdermins reveal an ancient mechanism of cell death. Science 375, 221–225. 10.1126/science.abj8432.

59. Wein, T., Millman, A., Lange, K., Yirmiya, E., Hadary, R., Garb, J., Melamed, S., Amitai, G., Dym, O., Steinruecke, F., et al. (2025). CARD domains mediate anti-phage defence in bacterial gasdermin systems. Nature 639, 727–734. 10.1038/s41586-024-08498-3.

60. Bernheim, A., Millman, A., Ofir, G., Meitav, G., Avraham, C., Shomar, H., Rosenberg, M.M., Tal, N., Melamed, S., Amitai, G., and Sorek, R. (2021). Prokaryotic viperins produce diverse antiviral molecules. Nature 589, 120–124. 10.1038/s41586-020-2762-2.

61. Garb, J., Lopatina, A., Bernheim, A., Zaremba, M., Siksnys, V., Melamed, S., Leavitt, A., Millman, A., Amitai, G., and Sorek, R. (2022). Multiple phage resistance systems inhibit infection via SIR2-dependent NAD(+) depletion. Nat Microbiol 7, 1849–1856. 10.1038/s41564-022-01207-8.

62. Bonhomme, D., Vaysset, H., Ednacot, E.M.Q., Rodrigues, V., Salloum, Y., Cury, J., Wang, A., Benchetrit, A., Affaticati, P., Trejo, V.H., et al. (2025). A human homolog of SIR2 antiphage proteins mediates immunity via the Toll-like receptor pathway. Science 389, eadr8536. 10.1126/science.adr8536.

63. Kowalski, M.P., Dubouix-Bourandy, A., Bajmoczi, M., Golan, D.E., Zaidi, T., Coutinho-Sledge, Y.S., Gygi, M.P., Gygi, S.P., Wiemer, E.A., and Pier, G.B. (2007). Host resistance to lung infection mediated by major vault protein in epithelial cells. Science 317, 130–132. 10.1126/science.1142311.

64. Ma, C., Luo, C., Deng, F., Yu, C., Chen, Y., Zhong, G., Zhan, Y., Nie, L., Huang, Y., Xia, Y., et al. (2024). Major vault protein directly enhances adaptive immunity induced by Influenza A virus or indirectly through innate immunity. Biochim Biophys Acta Mol Basis Dis 1870, 167441. 10.1016/j.bbadis.2024.167441.

65. Leipe, D.D., Koonin, E.V., and Aravind, L. (2004). STAND, a class of P-loop NTPases including animal and plant regulators of programmed cell death: multiple, complex domain architectures, unusual phyletic patterns, and evolution by horizontal gene transfer. J Mol Biol 343, 1–28. 10.1016/j.jmb.2004.08.023.

66. Cheng, T.C., Hong, C., Akey, I.V., Yuan, S., and Akey, C.W. (2016). A near atomic structure of the active human apoptosome. eLife 5. 10.7554/eLife.17755.

67. Bernheim, A., and Sorek, R. (2020). The pan-immune system of bacteria: antiviral defence as a community resource. Nat Rev Microbiol 18, 113–119. 10.1038/s41579-019-0278-2.

68. Yirmiya, E., Leavitt, A., Hurieva, B., Falkovich, A.H., Béchon, N., Rousset, F., Osterman, I., and Sorek, R. (2025). Systematic discovery of TIR-based immune signaling systems in bacteria. bioRxiv, 2025.2012.2003.692087. 10.64898/2025.12.03.692087.

69. Shi, Y., Kerry, P.S., Nanson, J.D., Bosanac, T., Sasaki, Y., Krauss, R., Saikot, F.K., Adams, S.E., Mosaiab, T., Masic, V., et al. (2022). Structural basis of SARM1 activation, substrate recognition, and inhibition by small molecules. Mol Cell 82, 1643–1659 e1610. 10.1016/j.molcel.2022.03.007.

