## Supplemental Materials for "TIR-like NADases act in bacterial immunity and the RNA vault"

**The PDF file includes:**

Materials and Methods
Figures S1 to S13
Tables S1 to S5
References

### Materials and Methods

#### Cell lines

FreeStyle™ 293F and ExpiF293 cells were purchased from Thermo Fisher Scientific. Sf21 and Sf9 cells were purchased from Expression Systems. *E. coli* DH5α cells were obtained from Invitrogen. *E. coli* Rosetta (DE3) cells were purchased from Novagen. *E. coli* BL21(DE3) cells were a generous gift from the Orth lab. *E. coli* BW251135 cells were a generous gift from the Harms lab.

#### Reagents

FreeStyle™ 293F (Thermo Fisher Scientific, R79007); Freestyle medium (Gibco 12338026); ExpiF293 cells (Thermo Fisher Scientific, A14527); Expi293 medium (Gibco, A1435102); Opti-MEM (Gibco, 31985070); Puromycin (Goldbio, P-600-100); G418 (Gibco, 10131035); Hygromycin (Corning, MT30240CR); E. *coli* Rosetta (DE3) cells (Novagen, 70954-3); *E. coli* DH5α cells (Invitrogen, 18258012); DH10Bac cells (Gibco, 10361012); Sf9 cells (Expression Systems, 94-001S); High Five Hi5 cells (Expression Systems, 94-002S); Sf21 cells (Expression Systems, 94-003S); ESF921 medium (Expression Systems, 96-001-01); Gibson Assembly (New England Biolabs, E2621); LB Brother, Miller (Fisher scientific, BP9723-500); Agar (Fisher scientific, BP1423-500); Terrific Broth (Research Products International, T15050); Antifoam B solution (Sigma-Aldrich, A5757); Ultra Yield flasks (Thomson Instrument Company, 931136-B); Ampicillin (Goldbio, A-301-100); Chloramphenicol (Sigma-Aldrich, C0378); Kanamycin (Goldbio, K-120-100); IPTG (Goldbio, I2481C50); Arabinose (Sigma-Aldrich, A3256); Imidazole (Sigma, I2399); Benzonase nuclease (Sigma-Aldrich, E1014-5KU); Glycerol (Fisher Scientific, G33-4); 0.22 µm PVDF filter (MilliporeSigma, SCGP00525); Bio Tek LogPhase600 (Agilent); flat-bottom clear 96-well plate (Fisher FB012931); Lysing Matrix B (MP Biomedicals, 116911100); FastPrep-24 instrument (MP Biomedicals, 11-600-5500); Amicon Ultra Centrifugal filter 3 kDA MWCO (Milipore UFC500308); Amicon 10K MWCO spin concentrators (Milipore, UFC901024); Amicon 100K MWCO spin concentrators (Milipore, UFC810024); Clear-bottom black 96-well plate (Greiner 655090); Spark Cyto plate reader (Tecan); cOmplete EDTA-free protease inhibitor (Sigma, 5056489001), DMEM (Fisher Scientific, 11-965-118); Fetal bovine serum (Sigma Aldrich, F-2442-500ML); Penicillin-streptomycin (Sigma Aldrich, P0781-100ML); PEI-MAX (Polysciences 24765); Norgen Biotek Phage DNA Isolation kit (Norgen Biotek 46800); NAD/NADH-Glo Assay kit (Promega #G9071); Strep-Tactin®XT 4Flow® high capacity resin (IBA Lifesciences 2-5030-025); D-Biotin (Goldbio B-950-100); NAD (Roche 10127965001); NADP (Roche 10128040001); BAD (BIOLOG B 301); ADPR (Sigma, A0752); ADPR(P) (BIOLOG, 53595-18-9); AMP-PNP (Sigma-Aldrich, 10102547001); Coomassie blue R-250 (Thermo Fisher Scientific, 20279); PEG 3,350 (Hampton Research, HR2-527); His-Pur™ Ni-NTA Resin (Thermo Scientific 88222); GeneJET Plasmid Miniprep Kit buffers (Thermo Scientific, FERK0503).

#### Discovery of similarity of the DUF4062 family to TIR-like domains.

To search for novel, distant relatives of the TIR-like superfamily, we analyzed the results of all-to-all protein sequence comparisons between all Pfam families (*1*), performed using the sequence profile-based method FFAS (*2*). For several member families of the TIR-like superfamily, the uncharacterized Domain of Unknown Function family DUF4062 stood out as a likely homolog for the TIR_2 family (PF13676; Z-score=-10.5), among others. Reciprocal searches using DUF4062 proteins as queries showed several TIR-like and Nucleoside 2-deoxyribosyltransferase-like families as significantly similar (Z-scores between -7 and -14). Both superfamilies (TIR and Nribosyltransf) are structurally similar, belonging to the flavodoxin-like fold (*3*). The FFAS results were corroborated by the HHpred method (*4*).

FFAS and HHpred sequence alignments allowed for unequivocal assignment of a conserved glutamate as a DUF4062 catalytic residue, corresponding to the known catalytic site of TIR-like NADases (*5*).

Using FFAS, the DUF4062 domains were identified in the multidomain human proteins TEP1, NWD1, NWD2, NPHP3 and in the likely pseudogene TTC41.

**Representative set of DUF4062 proteins and relationship of DUF4062 to other TIR-like families**

The Pfam definition of the DUF4062 domain is inaccurate, missing approximately half of the domain sequence. Thus, a representative set of DUF4062 domains was built from scratch, using own definition of a DUF4062 domain from *K. pneumoniae* as (NCBI ID: WP_341800075; residues 1-164; defined using the cryoEM structure of *Kp*Swz$^{15\text{-C, E97A}}$ obtained from NCBI ID: WP_202404386). This domain served as a query for 3 iterations of PSI-BLAST searches (0.001 E-value threshold) in the RefSeq Select database. Similar searches were performed using similarly redefined DUF4062 domains of human TEP1, NWD2 and NWD1 proteins as queries. Two sets of sequence hits (for *Kp*DUF4062 and for the human proteins) were independently cleared of redundancy with MMseqs clustering (*6*) at 0.8 sequence identity threshold and merged. The resulting sequence set (3907 sequences) was aligned using Mafft with standard parameters (*7*). The alignment was used to build a custom DUF4062 HMM sequence model using hmmbuild from the HMMER 3.4 package (*8*). Analogously, an additional HMM model was built for the divergent DUF4062 subfamily, NPHP3. Hmmsearch searches using these HMM models on the RefSeq protein database were performed and yielded 15814 and 1716 sequences, respectively. The two sets were merged, sequences shorter than 80 residues were discarded. Also discarded were sequences with overlapping higher significance hits to other Pfam domains. MMseqs clustering at 0.8 yielded the representative set of 3405 sequences.

To explore DUF4062 relationship to other flavodoxin fold families, the CLANS visualization and sequence-based clustering method was used (*9*), performing all-to-all BLAST comparisons with E-value threshold 0.01. For this analysis, representative sequences of each family from the STIR (TIR-like, Pfam ID: CL0173) and Nribosyltransf (CL0498) clans from the Pfam database were filtered (lengths > 80 aa and MMseqs clustering at 0.6 identity within each family), while the representative DUF4062 sequence set was further clustered using MMseqs at 0.6 identity yielding 1410 sequences.

#### Phylogenetic analysis of the DUF4062 family

For the phylogenetic analysis, the representative DUF4062 sequence set was further clustered using MMseqs at 0.4 sequence identity yielding 345 sequences. Mafft multiple sequence alignment was trimmed using ClipKIT v.1.4.1 (--mode gappy --gaps 0) (*10*). The IQtree v.2.1.4 phylogenenetic package was used (--alrt 1000 -B 1000) (*11*). The tree was visualised in iToL (*12*).

#### Survey of DUF4062 occurrence across the Tree of Life

The set of 192.8M proteins encoded by 25736 NCBI reference genomes (*13*) was searched by hmmscan (*8*) using the custom DUF4062 domain HMM profile as query. If the region of similarity to DUF4062 overlapped with similarity to another Pfam domain with better score, the protein was discarded. Downloading the sequence data and mapping to taxonomy was performed with the help of NCBI Datasets service (*14*).

#### Analysis of genomic features: defense islands, operons

Anti-phage defense systems in proximity to genes encoding DUF4062 proteins were identified using DefenseFinder v.1.3.0 (*15*). A DUF4062 protein was considered to lie within defense island if a defense system was identified within an arbitrarily selected distance of less than 20 genes which roughly corresponds to twice the reported median of defense island size, 17 genes (*16*). A DUF4062 protein was considered to lie near defense genes if any DefenseFinder gene hit was identified within the above distance, not requiring the presence of a full defense system. Putative DUF4062 defense operons were defined as co-directional sets of genes with intergenic distances below 50bp (a conservative cutoff, (*17*)) and with presence of at least one protein domain implicated in antiphage defense. Selected putative operons were validated using the Operon-mapper server (*18*).

#### Sequence alignment, sequence logo and auxiliary domains in DUF4062 family proteins

The sequence logo of the DUF4062 active site was built using the Weblogo method (*19*) on the multiple alignment of 345 representative DUF4062 sequences.

The DUF4062 domain multiple sequence alignment of selected DUF4062 domains was created by superposing respective AlphaFold 3 structure models (*20*) using FoldMason (*21*).

The auxiliary domains in DUF4062 proteins and in members of DUF4062 operons were identified using hmmsearch and Pfam database (version (37.4). Domain boundaries were verified by inspection of AlphaFold structure models.

#### Molecular cloning

Coding sequences for human NWD1 (isoform 3), NWD2, NPHP3, SARM1, TEP1, MVP and PARP4 were amplified by PCR from cDNA (Sequencing Core, UT Southwestern

Medical Center). Codon-optimization of TEP1 and PARP4 was done in SnapGene for Human codon frequency, manually edited to exclude common cloning sites, and synthesized in 4-5 fragments by Twist Bioscience for direct Gibson Assembly. The DUF4062-containing defense systems were codon-optimized in MacVector for Human codon frequency and synthesized by IDT (*Klebsiella pneumonia* UNIPARC: UPI001927123B, *Escherichia coli* UNIPARC: UPI0002A39D72, Acinetobacter baumannii UNIPARC: UPI0010579173, *Yersinia sp*. UNIPARC: UPI0000426B09, *Salmonella enterica* UNIPARC: UPI0021D4C267). Sequence for the portal protein from Bas14 bacteriophage (Bas14_003 gene, GenBank: MZ501107, Uniprot: A0AAE7VZF8) was amplified by PCR directly from the phage, or its escaper clones (1:50 dilution of phage stock in water was boiled for 10 minutes).

Mutations, tags and sequence modifications were introduced via QuikChange site-directed mutagenesis, KLD mutagenesis, In-Vivo Assembly, or Gibson Assembly.

For bacterial protein purification, *Kp*Swz was cloned into ppSUMO vector, a modified pET28a-based vector containing a 6XHis-Smt3 tag. TEP1 886-1113 and 895-1113 were cloned into ppSUMO in a similar fashion. Portal protein from Bas14 was cloned into pET28a with natural start codon, and a C-terminal 6xHis tag. pET28a/ppSUMO for expression of $SARM1^{TIR}$ and portal mutants were modified in the T7 promoter and ribosome-binding site according to Shilling et al. (*22*).

For anti-phage defense assessment, all constructs were tag-free. The anti-phage defense systems were cloned into pBbE6c vector downstream of the lac operator in place of RFP (Addgene #35287). Portal protein for sufficiency experiments was cloned into the pBAD vector, downstream of the arabinose promoter without tags.

For mammalian cell expression, NWD1 (isoform 3), NWD2, NPHP3 and TEP1 were cloned into pEZT-BM (BacMam) and pSBbiGP (Sleeping Beauty) vectors with N-terminal Twin-strep tag. For vault purification, MVP, TEP1 and PARP4 were cloned into pSBbiBH, pSBbiRN and pSBbiGP without tags, respectively. Tag-free overexpression of PARP4 native sequence was inefficient, however N-terminal TwinStrep-14xHis tag on the native sequence, or codon optimization alleviated the problem. Significantly better expression of MVP and PARP4 (codon optimized) was achieved in pEZT-BM vector series through transient transfection and was used for later expression experiments in Expi293.

**Phage propagation**

*E. coli* BW251135 strain was used for phage and toxicity experiments. For phage experiments, cells were grown in CMB (Calcium Magnesium Broth: LB-Miller Broth + 1.25 mM $CaCl_2$ + 1.25 mM $MgCl_2$) supplemented with antibiotics as indicated: ampicillin (100 µg/mL) or chloramphenicol (34 µg/mL).

Phages were amplified by dissolving three phage plaques in 300 µL of SM buffer (Saline Magnesium buffer; 100 mM NaCl, 8 mM $MgSO_4$, 50 mM Tris, pH 7.5), followed by four tenfold serial dilutions in SM buffer. Overnight cultures of BW251135 cells (50 µL) were mixed 1:1 with each phage dilution and allowed to incubate at 37 °C for 15 min. After phage adsorption, 3 mL of soft agar (CMB, 0.5% agar) was added to each bacterium–phage mixture and poured onto LB agar plates. Plates were allowed to solidify for 15 min and then incubated at 37 °C overnight. Plates displaying a semi-confluent bacterial lawn with a reticulated pattern of partial clearing (lacy lawn) were selected. Phages were harvested by adding 3 mL of SM buffer to the plate surface and gently rocking for 1 h. The resulting phage suspension was collected, centrifuged at 3,000 × g for 10 min to pellet residual bacterial cells, and filtered through a 0.22 µm PVDF filter.

**Phage plaquing assays**

To test phage defense, BW25113 cells transformed with either pBbA6C-RFP or plasmids encoding a putative defense gene were grown overnight in CMB supplemented with 34 µg/mL chloramphenicol. A 400 µL aliquot of each overnight culture was mixed with 3 mL of molten CMB soft agar (CMB, 0.5% agar) supplemented with 34 µg/mL chloramphenicol and 0.4 mM IPTG (isopropyl β-D-1-thiogalactopyranoside). The mixtures were poured onto LB agar plates supplemented with 34 µg/mL chloramphenicol and 0.4 mM IPTG and allowed to solidify at room temperature for 30 min.

Eight tenfold serial dilutions of phage stocks in SM buffer were prepared, and 2.5 µL of each dilution was spotted onto the prepared LB agar plates with bacterial lawns. The spots were allowed to dry for 15 min. Afterwards; the plates were incubated for 16 h at 25 °C. Plaque-forming units (PFUs) were determined by counting plaques after incubation. When individual plaques were not distinguishable, a faint lysis zone across the spotted area was scored as 10 plaques, whereas a dark, large lysis zone was scored as 50 plaques.

**Isolation of mutant phages that evade defense**

To isolate phage escaper mutants, plaque assays of Bas14 on BW25113 cells transformed with pBbA6C-*Kp*Swz$^{16\text{-}C}$ were performed. Individual escaper phage plaques were visually identified, picked with a pipette tip into 300 µL of SM buffer, and tenfold serial dilutions in SM buffer were prepared. Overnight cultures of BW25113 cells transformed with pBbA6C-*Kp*Swz$^{16\text{-}C}$ (50 µL) were mixed 1:1 with each phage dilution and incubated at 37 °C for 15 min. Phage clones were plated and propagated as before. Phage gDNA was extracted using the Norgen Biotek Phage DNA Isolation kit (Norgen Biotek 46800). The extracted gDNA from Bas14 parental strain and escapers were sent for whole bacterial genome sequencing using Illumina sequencing (Performed by SeqCenter, Pittsburgh, PA) and the variants were called by mapping against the wild-type Bas14 genome (MZ501107) using BreSeq (*23*).

**Infection assays in liquid cultures**

Overnight cultures of BW25113 cells transformed with either pBbA6C-RFP or indicated *Kp*Swz plasmids were diluted 1:50 into CMB supplemented with 34 µg/mL chloramphenicol and 0.4 mM IPTG. The cultures were grown to an OD600 of 0.8 and subsequently diluted to an OD600 of 0.2 using the same media. 100 µL of each dilution was transferred into a flat-bottom clear 96-well plate. Bas14 phage dilutions or SM buffer (negative control) were added to each well to achieve final MOIs of 0, 0.001, or 10. Plates were transferred to a Bio Tek LogPhase600 and incubated at 37 °C while orbital shaking at 800 rpm. Bacterial growth was monitored by measuring OD600 every ten minutes.

To assess abortive infection phenotypes, parallel infections were performed at an MOI of 10 in a second 96-well plate. After observation of bacterial culture crash at an MOI of 0.02 (~4 h post-infection), each well was collected and centrifugated at 15,000 RCF for 15 min. The phage containing supernatants were collected and a plaque assay was performed on BW25113 cells transformed with pBbA6C-RFP to determine phage titers.

**NAD level measurement in bacterial culture**

Overnight cultures of BW25113 cells transformed with either pBbA6C-RFP or indicated *Kp*Swz plasmids were diluted 1:50 into 30 mL of CMB supplemented with 34 µg/mL chloramphenicol and 0.4 mM IPTG. Cultures were grown at 37 °C until reaching an OD600 of 0.3 and then infected with Bas14 phage at an MOI of 5. Infection was allowed to proceed while shaking at 37 °C for 30 min. Afterwards, 20 mL of each culture were transferred to a conical tube, centrifuged at 3400 RCF for 10 min, and the supernatant was discarded. The pellets were resuspended in 600 µL of resuspension buffer (50% methanol (v/v), 6 mM $NaH_2PO_4$, and 94 mM $Na_2HPO_4$) and snap frozen in liquid nitrogen.

Frozen samples were thawed at room temperature and 500 µL was transferred to a 2 mL tube containing lysing matrix B (MP Biomedicals). Mechanical lysis was performed using a FastPrep-24 instrument (40 s total time, 6 $ms^{-1}$). Samples were centrifuged at 15,000 RCF for 10 min at 4 °C. The supernatants were transferred to Amicon Ultra Centrifugal filters with a 3 kDa molecular weight cut off (Milipore UFC500308). The samples were centrifugated at 14,000 RCF for 1.5 h at 4 ° C and then the filtrates were snap frozen in liquid nitrogen for future analysis.

Relative NAD levels were determined using the NAD/NADH-Glo Assay kit (Promega #G9071). Samples were thawed at room temperature and 25 µL of each sample transferred to the wells of a clear-bottom black 96-well plate (Greiner 655090). Subsequently, 25 µL of NAD/NADH-Glo reagent was added to each well and incubated at room temperature for 15 min. Luminescence values were measured using a Tecan Spark Cyto plate reader.

**Bacterial protein expression and purification**

*Kp*Swz (His-SUMO-*Kp*Swz$^{15\text{-}C}$ and the E97A mutant) SARM1$^{TIR}$ (561-700), AbTIR, HopAM1 and TEP1$^{TIR}$ (886- and 895-1113) truncations were produced in *E. coli* Rosetta(DE3) or BL21(DE3). Overnight inoculum cultures (5% of the production culture volume) were grown in LB medium with Kanamycin (50 µg/mL; Rosetta(DE3) were additionally supplemented with 34 µg/mL Chloramphenicol). Cultures were used to inoculate Terrific Broth medium, supplemented with 0.5% Glycerol, ~13 drops of Antifoam B solution, 1.5 mM $MgSO_4$, and appropriate antibiotics in 2.5 L Ultra Yield flasks, 1 L per flask. Cultures were grown at 37 °C for ~3-4 h with 220-250 RPM, until OD600 was 1.0 or more.

For *Kp*Swz and TEP1$^{TIR}$, temperature was decreased to 4 °C for ~3 h with constant shaking to allow for continued bacterial growth, after which the temperature of the culture medium was measured. When culture medium reached 12-15 °C, IPTG was added to 0.4 mM, and incubators were set to 12 °C. Expression of the mutant proteins did not require strict temperature control. Yield for the wild-type TEP1$^{TIR}$ protein was especially improved by the low temperature, from ~150 μg/L of TB medium to >2.5 mg/L. Nicotinamide supplementation had no effect. Cultures were harvested by centrifugation (15 minutes, 3,500 RCF), and bacterial pellets were resuspended in 50 mM Tris 8.0, 300 mM NaCl, 4 mM β-ME, 1 mM PMSF, 15 mM imidazole. Lysates were produced by sonication, and cleared by centrifugation (30 min, 35,000 RCF). Ni-NTA resin was added to the lysates and incubated for 0.5-1 h with gentle rocking in 50 mL falcon tubes. Protein-bound resin was spun 1,000 RCF for 5 minutes, resuspended in wash buffer (lysis buffer supplemented with imidazole to final 30 mM) and spun again. Resin was loaded onto gravity column and washed with wash buffer. Additional washes were done with 50 mM Tris 8.0, 1 M NaCl, 4 mM β-ME, and 30 mM imidazole.

For TEP1$^{TIR}$, protein was eluted with 300 mM imidazole in wash buffer, cleaved with 6His-Ulp1 protease, and dialyzed against 50 mM Tris 8.0, 50 mM NaCl, 1 mM DTT crystallization buffer overnight. Protein was passed over HiTrap CaptoQ column equilibrated with crystallization buffer. Flow-through and wash fractions containing TEP1$^{TIR}$ were collected and passed over HiLoad S75 column equilibrated with the crystallization buffer. Fractions containing TEP1$^{TIR}$ were concentrated to 15-40 mg/mL using Amicon 10K MWCO spin concentrators. TEP1$^{TIR}$ E1008A protein was less soluble than the wild-type, with ~15 mg/mL stable concentration after a freeze-thaw. All proteins were stored at -80 °C.

*Kp*Swz protein was eluted on-resin GST-Ulp cleavage to avoid the contaminants. The Ni-NTA resin was washed with TBS-βME (50 mM Tris 8.0, 150 mM NaCl, 4 mM β-ME) to remove the imidazole. Resin was resuspended in equal volume TBS-βME and incubated with ~20-40 ug of GST-Ulp1 protease at room temperature for ~1 h to cleave the SUMO tag and release the untagged protein (protease was not his-tagged to avoid sequestration by Ni-NTA resin). Flow-through was collected, and resin was washed with 5 volumes of TBS-βME + 30 mM imidazole to release the non-specifically bound *Kp*Swz. Pooled fractions were concentrated and separated on S200 increase column. Fractions were concentrated to ~3 mg/mL and stored at -80 °C. Yield for *Kp*Swz$^{15\text{-}C}$ WT was ~150 μg/L.

Interestingly, we were able to remove *Kp*Swz toxicity entirely by using VNp15-14xHis-NeonGreen-3C tag (*24*). However, this protein was not used for any experiments in this manuscript.

AbTIR (Uniprot: A0A009IHW8), Human SARM1$^{TIR}$ (561-700, Uniprot: Q6SZW1) and HopAM1 (Uniprot: Q877R9) were produced in a similar fashion, however induction was performed at 18 °C. Proteins were eluted with buffer supplemented with 300 mM imidazole, cut with 6His-Ulp1 overnight, and passed over S200 increase size exclusion column in 50 mM Tris, 150 mM NaCl, 1 mM DTT.

Portal protein (Bas14, Portal-6His) cultures were inoculated into TB medium as above, and grown at 37 °C. After reaching OD600 ~1, incubator was set to 18 °C, and 1 hour later IPTG was added to 0.4 mM. Next day, cultures were harvested as above, and resuspended in 50 mM Tris 8.0, 400 mM NaCl, 1 mM $MgCl_2$, 4 mM β-ME, 15 mM imidazole, 1 mM PMSF, and supplemented with ~20 µL of Benzonase nuclease. Sonicated lysates were cleared as above. Lysates were incubated with Ni-NTA resin for ~1 h, and washed with 50 mM Tris 8.0, 1 M NaCl, 4 mM β-ME, 30 mM imidazole. Protein was eluted in 50 mM Tris 8.0, 400 mM NaCl, 300 mM imidazole, 1 mM DTT, and passed over Superose 6 column in 50 mM Tris, 150 mM NaCl, 1 mM DTT. Peak fractions were concentrated and stored at -80 °C.

**Stable cell line generation (Freestyle 293F)**

293F cells for purification of NWD1 (isoform 3), NWD2, NPHP3, TEP1, and TEP1/PARP4/MVP were stably integrated with the Sleeping Beauty transposase system, adapting the original protocol to suspension culture (*25*). 293F cells in actively growing cultures were centrifuged at 300 RCF and resuspended in 30-50 mL of fresh Freestyle medium. Transfection was performed by mixing 1 ug of transposon-containing plasmid, 0.l ug of Sleeping Beauty transposase plasmid and 3 ug PEI-MAX (40K, Polysciences) per mL of culture (~50 ug DNA, 150 ug PEI-MAX per 50 mL of culture). Plasmids were added first, and after 5 minutes of incubation PEI-MAX was added. Alternatively, DNA-PEI complexes were pre-formed in Opti-MEM and added to cultures after 15 minutes of incubation. Cultures were returned to the incubator for 3 days, after which medium was replaced and selection antibiotics were added (depending on the transposon cassette, 3 µg/mL puromycin, 400 µg/mL G418 or 400 µg/mL Hygromycin). Cells were passaged under selection until fluorescent markers from the cassettes were abundantly expressed, and culture viability was restored to >95%. For protein purification, cultures were scaled up and harvested upon reaching the stationary phase around 3.5 e6/mL. Stable cell lines were also successfully generated in adherent 293F cultures grown in DMEM +10% FBS, using conditions from Aronovich et al., (*25*).

**BacMam generation and transduction**

Bacmids were generated from pEZT-BM plasmids by transforming DH10Bac cells and blue-white screening, as per pFastBac manufacturer instructions. Bacmids were isolated from 5 mL cultures by lysing the cells with GeneJET Plasmid Miniprep Kit buffers. Bacmids were precipitated directly from the lysates with 50% isopropyl alcohol, spun down, and washed with 70% ethanol. Crude bacmids were dissolved in 100 µL of miliQ $H_2O$, verified by M13/pUC primer PCR, quantified by nanodrop, and frozen for long term storage.

Sf9 cells were seeded in T25 flasks directly from a live suspension culture in ESF921 medium. 8 e6 cells per flask were added to each flask in ~2 mL of medium and incubated for ~20 minutes to allow attachment. Media was replaced with fresh 5 mL of ESF921 to aid transfection. 8 ug of the bacmid DNA were diluted into 200 µL of ESF921. 16 ug of PEI-MAX was diluted in 200 µL of ES921 and added to the diluted bacmid DNA (8 ug of bacmid + 16 ug of PEI-MAX). After 15 minutes, DNA:PEI complexes were added to the

T25 flasks and gently mixed with the medium. Transfection was monitored by GFP marker fluorescence from pEZT-BM GFP cassette. After 5 days, the medium was collected (P1) and added to 50 mL of Sf9 culture at 3 e6/mL. After 5 days the cells were spun down and supernatant (P2) was stored for large-scale virus production. For P3 generation, ~200 mL of Sf9 culture was transduced with 2 mL of P2 supernatant. After 5 days, medium was collected and stored at 4 °C.

293F cells were seeded in fresh Freestyle medium at 3 e6/mL. BacMam virus supernatant was added to ~10% final culture volume. Next day, 5 mM Sodium Butyrate was added. Cells were harvested for protein purification 3-5 days post transduction.

**Human DUF4062-containing protein purification.**

Harvested 293F cell pellets were resuspended in cold 50 mM Tris 8.0, 300 mM NaCl, 1 mM DTT, 5% Glycerol and 1 mL of Biolock solution per liter of culture (crude chicken egg avidin preparation, IBA Lifesciences). Resuspended cells were diluted 1:1 with a 2% NP-40, 2 mM PMSF, 2x cOmplete™ Protease Inhibitor Cocktail (EDTA-free) in the resuspension buffer. Lysate was vortexed, sonicated briefly to reduce viscosity, and centrifuged 35,000 RCF for 30 minutes. Cleared lysate was incubated with 1 mL of Strep-Tactin®XT 4Flow® high-capacity resin (IBA Lifesciences) for 1 hour. Resin was separated from the lysate by centrifugation (5 min, 1000 RCF), resuspended in the lysis buffer and spun again. Resin was applied onto a gravity column, and washed further with 50 mM Tris 8.0, 150 NaCl, 1 mM DTT, 5% glycerol. Proteins were eluted with wash buffer containing 50 mM biotin. TEP1, NWD2 and NPHP3 were further purified over S200 increase size-exclusion column. All proteins were concentrated and stored at -80 °C.

All affinity-purified full-length human DUF4062-containing proteins contained N-terminal TwinStrep (BacMam), or TwinStrep-14xHis tag (Sleeping Beauty). NWD2 yields were in the milligram/L range for both the Sleeping Beauty and BacMam systems. NPHP3 was made from the Sleeping Beauty transduced cells with acceptable yields and could also be made in Hi5 cells from pFastBac system. TEP1 could be purified from both BacMam and Sleeping Beauty, with yields varying ~50-150 µg/L. NWD1 isoform 3 was made using Sleeping Beauty system and expressed poorly.

**NADase experiments**

All NADase experiments were performed in buffer containing 50 mM Tris 8.0, 50 mM NaCl and 1 mM DTT in 20 µL reactions. Proteins purified as described were diluted in the reaction buffer to 50-90% volume of the reaction. Amount of proteins was 10 µg of full-length human protein preparation, 15 µg of the vault preparation, 1 µg of isolated domains, unless indicated otherwise. Reactions were started by addition of $NAD^+$/$NADP^+$ to final 5 mM (or indicated concentration). Reactions were stopped after overnight incubation at 37 °C by addition of 300 µL of HPLC buffer (75% acetonitrile, 20 mM Ammonium Acetate ($NAD^+$) or Ammonium Bicarbonate ($NADP^+$), pH 9). Reactions were spun down at 21,300 RCF to remove precipitated proteins and separated on Waters UHPLC system with a UV detector set to monitor absorbance at λ=260 nm. The column used was Atlantis Premier BEH Z-HILIC Column, 1.7 µm, 2.1 mm X 50 mm (Waters

186009978). Columns with the VanGuard filter were not appropriate for the separation. 3 µL of solution was injected using the autosampler. Isocratic elution was performed at 0.5 mL/min at 30 °C, in 75% acetonitrile with 20 mM Ammonium Acetate ($NAD^+$) or Ammonium Bicarbonate ($NADP^+$).

Assay comparing activity of known TIR domains (AbTIR, HopAM1, and $SARM1^{TIR}$) was performed with 3 µg of each protein in 10 µL reactions with 1 mM $NAD^+$ for 1 h at 37 °C. Assay was stopped by addition of 10% final TCA (~1 uL of 100% solution to stop 10 µL reaction), centrifugated at 21,300 RCF, and diluted with 250 uL of HPLC buffer A: 100 mM Potassium Phosphate pH 6. Separation was performed on Atlantis Premier BEH C18 AX Column, 1.7 µm, 2.1 x 100 mm (Waters 186009368) at 0.5 mL/min, 30 °C. 3 µL of solution was injected using the autosampler. Buffer B was 100 mM Potassium Phosphate pH 6 in 10 % methanol. Separation gradient was: 0-2 min 100% A, 2-6 min linear gradient 0-100% B, 6-7 min 100% B, 7.0 min 100% A for a 12 minute total run time.

$TEP1^{TIR}$ enzyme concentration turnover dependency was measured by incubating $TEP1^{TIR}$ from a two-fold dilution series between 20-0.078 µM with additional points at 15 µM, 7.5 µM and 3.75 µM with 10 mM $NAD(P)^+$ in 10 µL reactions that included 1 mg/mL BSA. After 10 minutes, reactions were stopped with 2 µL TCA and diluted with 250 µL of buffer A. Separation was performed on C18 AX column as before, but pH of the phosphate buffer was 8.0.

For kinetic assays of *Kp*Swz with Portal, *Kp*Swz$^{15\text{-}C}$ at 0.15 µM was pre-incubated with 7 different concentrations of portal$^{Bas14}$ from a 2-fold dilution series starting at 2 µM for 1 h, and a buffer control. Protein dilutions were prepared in buffer containing 0.1 mg/mL BSA. To start the reactions, NAD was added to final 5 mM in the 10 µL reactions. After 30 minutes at ambient temperature, triplicate reactions were stopped by addition of 1 µL of 100% TCA. Reactions were centrifugated for 10 minutes at 21,300 RCF. 9 µL of the reaction was mixed with 191 µL of buffer A. Reactions were separated on C18 AX column as above, pH of the phosphate buffer was 8.0. Area under the curve was quantified using Empower software (Waters) based on fresh $NAD(P)^+$ and ADPR(P) standards, prepared like the reactions. Quantified ADPR(P) amounts were additionally adjusted by subtracting the ADPR(P) contamination quantified within the $NAD(P)^+$ standards.

Km determination with $TEP1^{TIR}$ were performed at 1 µM enzyme in 10 µL reactions. 5 µL of enzyme was mixed with $NAD^+$ or $NADP^+$ solution, incubated for 15 minutes, and stopped with 2 µL of TCA. 250 µL of buffer A was added, and 3 µL of reaction was separated on C18 AX column as above, in phosphate buffer pH 6.0. Concentrations of nucleotide were 10, 5, 4, 3, 1.5, 1, 0.75, 0.5, 0.325, 0.25, 0.1625, 0.125 mM, formed by 2-fold dilution series from appropriate starting concentrations.

**KpSwz-portal$^{Bas14}$ complex formation**

150 µg of the portal$^{Bas14}$ was mixed with 150 µg of *Kp*Swz$^{15\text{-}C,\ E97A}$. Proteins were incubated for ~30 minutes at ambient temperature and separated on Superose 6 increase column equilibrated with TBS-DTT, measuring A280. Fractions from the complex and

control runs were separated on 12% SDS-PAGE gels and stained with Coomassie blue R-250.

### Bacterial spotting assay (Portal sufficiency)

*E. coli* BW25113 were transformed with indicated plasmids (pBbE6c with RFP or *Kp*Swz$^{16\text{-}C}$ and indicated mutants, pBAD empty, or with portal$^{Bas14}$ with indicated mutations). Cultures were grown from single colonies in 5 mL of LB overnight at 37 °C. Bacteria were harvested by centrifugation at 4000 RCF for 5 minutes. Bacteria were resuspended in 2 mL of 150 mM NaCl solution (sterile filtered). Bacteria were diluted 10-fold, and the Optical density at 600 nm was measured using a spectrophotometer. Optical density was adjusted to 0.5, and 10-fold dilution series were prepared by pipetting 50 µL of bacteria to 450 µL of saline. 10 µL of bacterial dilution were manually spotted on LB plates containing 34 µg/mL chloramphenicol, 100 µg/mL ampicillin, and either 1% of glucose, or indicated concentration of IPTG (0.4 mM) and arabinose (0.00002%). Arabinose concentration had to be lowered to avoid toxicity arising from strong portal overexpression.

### RNA vault purification

Human RNA vault purifications were performed with tag-free proteins. Protocols were focused on increasing the amount of PARP4 and TEP1 relative to MVP and decreasing the amount of contaminating ribosomes without using any nucleases. Protocols for each preparation are described separately.

We were unable to successfully purify appreciable amounts of C-terminally His-tagged vaults from human cells, despite their good expression. Similarly, C-terminal FLAG and 2xHA worked poorly. Effect is likely similar to the observations by Hananya et. al. (*26*) N-terminal tags were not attempted to avoid enrichment of half-vaults.

### RNA vault purification with PEG 3,350: CryoEM sample MVP/TEP1 + BAD + AMP-PNP

293F cell line stably overexpressing MVP (pSBbiBH), TEP1 (pSBbiRN, native sequence) and PARP4 (pSBbiGP, native sequence with no tag, no visible overexpression above native PARP4) were resuspended in 300 mL of fresh Freestyle media at a density of 3 e6/mL. To the media, plasmids were added, amounts per mL of culture: pSBbiRN TEP1 (1.25 ug), pSBbiGP PARP4 (1.25 ug), pSBbiBH MVP (0.5 ug) and returned to the incubator. After 5 minutes, 9 ug of PEI-MAX was added into the culture, per mL of media (for 300 mL of culture 900 ug total plasmid and 2700 ug PEI-MAX). Day after transfection, culture was supplemented with 5 mM Sodium Butyrate. Cells were harvested on day 3 after transfection, resuspended in 12 mL MVP buffer: 50 mM Tris 7.5, 75 mM NaCl, 1.5 mM $MgCl_2$, 1 mM DTT, and transferred to a 50 mL conical tube. Resuspended cells were diluted 1:1 with 12 mL of MVP buffer containing 2 % NP-40, 2 mM PMSF and 2x cOmplete protease inhibitor (EDTA-free), vortexed, sonicated briefly and centrifugated for 20 minutes at 20,000 RCF. Supernatant volume was measured with a pipette, and 50% w/v solution of PEG 3,350 was added to a final PEG concentration of 7%, with optimal concentration between 6-8% depending on the batch of lysate. PEG precipitates most ribosomes in this condition, however vaults tend to precipitate at a similar concentration, ~8% and above. Therefore, optimization of the PEG concentration is recommended. Solution was vortexed and kept on ice for 30 minutes, after which centrifugation was repeated for 20 min at 20,000 RCF. Supernatant was collected and overlayed on top of 1.5 mL of 30% sucrose cushion prepared in MVP buffer in Thin Wall Polyallomer Tubes (~12 mL volume, Seton 5030). Samples were spun at 37,000 RPM in TH-641 rotor for 2 h at 4 °C. Supernatant was removed with vacuum, and small volume of the MVP buffer was added to the pellets (total 1.5 mL per culture lysate). Pellets were broken up from the bottom of the tube with disposable inoculation needles and transferred with a cut P1000 tip into a 2 mL glass dounce. Pellets were resuspended by douncing on ice and transferred into two 1.5 mL tubes. Resuspended vaults were cleared by centrifugation in a tabletop centrifuge (~10 minutes, 21,300 RCF). Soluble fraction was transferred to fresh tubes, and pellets were reextracted by vortexing with 200 µL MVP buffer, and centrifugation. Supernatants were overlayed on top of a stepwise sucrose gradient prepared in MVP buffer. Sucrose solutions were pre-chilled and overlayed on top of each other with a cut P1000 in a Ultra-Clear™ thin-wall tube (Beckman Coulter 344059). Gradient composition was 2 mL 55%, 2 mL 50%, 2 mL 45%, 2 mL 40%, 2 mL 35%, 1 mL 30% of sucrose. Phase boundaries were marked with a sharpie, and vault sample was overlayed on top. Samples were centrifugated at 30,000 RPM in TH-641 rotor for 16 h at 4 °C. Fractions were separated one-by-one by submerging uncut P1000 tip to the first marked boundary line. 0-35% fractions were saved as waste (ribosome-enriched). The 40% and 45% fractions were pooled in a 15 mL Amicon concentrator (100K MWCO), diluted to 15 mL with MVP buffer to reduce sample density and concentrated. Sample was loaded onto Superose 6 column equilibrated with MVP buffer. Void fraction was collected, concentrated, quantified by Bradford reagent and stored at 4 °C until use. PARP4 was not observed on the SDS-PAGE gel of this preparation, consistent with negligible expression of PARP4 from the native-sequence plasmid used.

**RNA vault purification with high salt method: CryoEM sample MVP/PARP4/TEP1 + NADP**

Samples were transfected into as for MVP/TEP1 sample, with plasmids for MVP, TEP1 and PARP4. Cells were stably overexpressing codon optimized TEP1 (pSBbiRN) and codon optimized PARP4 (pSBbiGP). Protocol for ribosome removal was adapted from Rivera, et al. (*27*). Cell pellets were resuspended in 50 mM Tris 8.0, 500 mM NaCl, 5 mM $MgCl_2$, 1 mM DTT. Resuspended pellet was diluted 1:1 with resuspension buffer supplemented with 2 % NP-40 and 2x cOmplete protease inhibitor. Puromycin was added to 0.1 mg/mL, and additional $MgCl_2$ was added to 50 mM. Lysate was sonicated and incubated for ~30 minutes on ice. Lysate was clarified by centrifugation for 20 min at 20,000 RCF. Supernatant was overlayed on top of 1.5 mL 30% sucrose cushion prepared in high salt buffer (50 mM Tris 8.0, 500 mM NaCl, 50 mM $MgCl_2$, 1 mM DTT) and centrifugated for 2 h at 37,000 RPM in TH-641 rotor. Pellets were resuspended in high salt buffer with 0.2 mg/mL puromycin using the dounce method. Resuspended vaults were incubated at room temperature for ~15 minutes, vortexed briefly and clarified in a tabletop centrifuge (10 min, 21,300 RCF, 4 °C). Soluble fraction was overlayed on top of a sucrose gradient prepared in MVP buffer, with sucrose percentages as in the PEG method. Equilibrium centrifugation was performed as before, and vaults were polished over Superose 6 column in MVP buffer. Samples were concentrated and stored at 4 °C.

Alternative method for ribosome removal from Rivera, et al. (*27*) used high amount of EDTA instead of $MgCl_2$ to dissociate the ribosomes. It was similarly successful in the pilot vault purifications, however no experiments were performed with vaults prepared with EDTA.

**RNA vault purification: CryoET samples: MVP, MVP/PARP4, MVP/TEP1**

293F cell lines stably overexpressing MVP, MVP/PARP4 and MVP/TEP1 were transfected with additional plasmids, as for MVP/PARP4/TEP1 sample preparation. Samples were purified like for MVP/PARP4/TEP1 sample, using the high-salt method. Instead of the MVP buffer, TBS-DTT was used.

**RNA vault purification: Activity assays**

MVP/TEP1 WT and MVP/TEP1 E1008A were transfected and purified as described for the PEG method. Cell lines were stably expressing TEP1 WT or TEP1 E1008A from integrated pSBbiRN vectors. PARP4 was not appreciably present in those purified vaults. Preparation was initially tested for activity and then stored in -80 °C before use alongside the below described samples.

MVP, MVP/PARP4, MVP/PARP4/TEP1 WT and MVP/PARP4/TEP1 E1008A were purified using the high-salt method from transiently transfected wild-type Expi293 cells. Expi293 cells from live culture at 5-7 e6/mL were spun down and resuspended in 90% of final culture volume by brief vortexing in fresh warm Expi medium (270 mL for 300 mL transfection). Cell density in the final culture volume was ~3 e6/mL. Transfection mixtures were prepared in warmed, room-temperature Opti-MEM medium. 10% of culture volume of room-temperature Opti-MEM was split into two conical tubes (for 300 mL of transfection, 2x 15 mL). DNA was diluted in one half, PEI-MAX in the other, the two were mixed, and incubated for 15 minutes, with occasional mixing. Transfection mixture added to cells contained:

MVP: 2 ug pEZT-BM MVP + 4 ug PEI (per mL of final culture)

MVP/PARP4: 1 ug pEZT-BM MVP + 1 ug pEZT-BM PARP4 (codon optimized) + 4 ug PEI-MAX (per mL of culture)

MVP/PARP4/TEP1WT/E1008A: 0.4 ug pEZT-BM MVP + 0.4 ug pEZT-BM PARP4 (codon optimized) + 2.2 ug pSBbiRN TEP1 (codon optimized) + 6 ug PEI-MAX

Transfections were supplemented with 5 mM Sodium Butyrate 1 day post transfection, harvested on day 3 post transfection, and processed with the high salt method. Instead of MVP buffer, TBS-DTT was used.

### CryoEM grid preparation

*Kp*Swz$^{15\text{-}C\ E97A}$ was purified as above. Vitrification was performed in liquid ethane using Vitrobot MK. IV. Quantifoil Cu R1.2/1.3 M300 grids were glow discharged (PELCO easiGlow™) and 3 µL of 2 mg/mL sample in TBS-DTT buffer were applied. Grids were screened on a TFS Glacios.

To obtain *Kp*Swz- portal$^{Bas14}$ structure, portal$^{Bas14}$ was mixed with excess (1:1.5) of mutant *Kp*Swz$^{15\text{-}C\ E97A}$. Mixture was incubated for ~30 minutes at ambient temperature and separated on Superose 6 size exclusion column in 50 mM Tris 8.0, 150 mM NaCl, 1 mM DTT. Peak fraction was concentrated and stored overnight at 4 °C before grid preparation. Sample at 4 mg/mL was supplemented with 0.025% DDM and vitrified as above.

vaults at ~4 mg/mL were supplemented with 0.025% DDM, 1 mM BAD and 1 mM AMP-PNP (MVP + TEP1 sample) and vitrified as above.

For MVP + PARP4 + TEP1 sample with $NADP^+$, vaults were pre-diluted in 1.5 mL test tubes and kept on ice to avoid $NADP^+$ hydrolysis by TEP1. ~1 minute before freezing, 1 mM NADP and 0.025% DDM was added to the samples with 4.1 mg/mL and 3 mg/mL of the vaults. Sample with 4.1 mg/mL was frozen first, using a double application protocol (3 µL of solution were applied to the grid, manually blotted, another 3 µL were applied onto the grid and blotted with the Vitrobot). The 3 mg/mL sample was frozen ~2 minutes later using single application of 3.5 uL. Data was collected from both grids during a 3-day microscope session (1 day low density, 2 day high density), and different conformations of the RNA vault were observed between the two grids. It is unclear what caused the difference in the vault conformation distribution between the two grids; however it is most likely the vault concentration, the incubation time with NADP/DDM, or NADP:MVP ratio. The more uniform, compact conformation of the vault was observed at the lower concentration of the vault after a longer incubation time with a higher nucleotide ratio. However, the exact reason for the difference requires further investigation.

### CryoEM data acquisition

All data collections were performed on TFS Krios microscopes using SerialEM. The pixel sizes for MVP datasets were recalibrated as described in data processing method section.

*Kp*Swz data was collected on a TFS Krios at UT Southwestern Medical Center Cryo-Electron Microscopy facility. K3 Bioquantum detector was used in CDS mode, with 3 shots-per-hole on a 3x3 matrix (27 movies per stage shift). Magnification was 105,000x with detector pixel size of 0.83 Å. Defocus range was 1-2.4 µm, and approximate dose 52 $e^-/Å^2$ in a 4.5 s exposure. Energy filter was set to 20 eV width.

*Kp*Swz-Portal$^{Bas14}$ data was collected on a TFS Krios at UT Southwestern Medical Center Cryo-Electron Microscopy facility. K3 Bioquantum detector was used in CDS mode, with 3 shots-per-hole on a 3x3 matrix (27 movies per stage shift). Magnification was 105,000x with detector pixel size of 0.826 Å. Defocus range was 0.9-2.2 µm, and approximate dose 50 $e^-/Å^2$ in a 1.8 s exposure. Energy filter was set to 20 eV width.

MVP+TEP1+BAD+AMP-PNP data was collected on a TFS Krios at UT Southwestern Medical Center Cryo-Electron Microscopy facility. Falcon IV detector was used, with 4 shots-per-hole on a 3x3 matrix (36 movies per stage shift). Magnification was 165,000x with detector pixel size of 0.738 Å. Defocus range was 0.9-2.2 µm, and approximate dose 50 $e^{-}/Å^{2}$ in a 3.5 s exposure. Energy filter was set to 20 eV width.

MVP+TEP1+PARP4+$NADP^{+}$ data was collected on a TFS Krios in HHMI Janelia EM Facility. Falcon IV detector was used, with 6 shots-per-hole on a 3x3 matrix (54 movies per stage shift). Magnification was 165,000x with detector pixel size of 0.743 Å. Defocus range was 0.9-2.2 µm, and approximate dose 50 $e^{-}/Å^{2}$. Energy filter was set to 6 eV width.

### CryoEM data processing

#### ***Kp*Swz$^{15\text{-}C,\ E97A}$ apo structure**

Raw movies were aligned using MotionCorr2 in RELION 3.1.3. Movies were imported into Cryosparc 3, where patch CTF estimation was performed. All further reconstruction steps were performed in cryoSPARC with C2 symmetry applied, while 3D classifications were done in RELION. Initial particles were picked using blob picker and extracted. After 2D classification, best 2D classes were used as templates for template picking. Additional particles were picked using crYOLO. All batches of picked particles were subjected to independent 2D classification. Good classes were pooled, and duplicates removed. 2D classes for dimeric, tetrameric and hexameric forms of protein were observed. Dimeric 2D classes were rejected, as they didn't refine to atomic resolutions. Tetrameric and hexameric particles were pooled and separated by 3D classification with alignment in RELION 3.1.3. Tetrameric class was aligned to C2 symmetry in cryoSPARC. Hexameric class was aligned to C2 symmetry using relion_align_symmetry. Symmetry alignment of the hexameric class required increasing orientation sampling, due to incorrect alignment with default parameters. Incorrect alignment resulted in shadowing of the hexameric class into a distorted octameric class upon refinement in C2 symmetry. Both classes were imported into cryoSPARC 3 for reconstruction and CTF refinement, and exported into RELION for 3D-classification without alignment. Final particles were subjected to Bayesian polishing in RELION, and imported into cryoSPARC 4 for non-uniform refinement, CTF refinement, finally followed by local refinement in C2 symmetry. Resolution estimates for the hexamer and tetramer were 2.42 Å and 2.66 Å, respectively.

#### ***KpSwz*$^{15\text{-}C,\ E97A}$ with portal$^{Bas14}$**

All steps were performed in cryoSPARC 4.7.1; 7,689 movies were imported and subjected to patch motion correction and patch CTF. Beamshift was imported from the *.mdoc files recorded by serialEM, after conversion with mdoc_xml.py script by ccgauvin94. Initial 200 movies were used to generate 2D templates for template picking during the data collection. Ab-initio reconstruction with 2 classes was performed from 20K particles. Good class was used to refine all particles in C12 symmetry, and handedness was corrected. Lowpass filter was set to 12 Å for all further refinement steps to avoid hand-flipping. The ~1.9M particles picked were subjected to 2D classification, resulting in 1.2M particles. All 1.2M particles were then refined against the initial map in C12 symmetry. Two masks were generated manually in UCSF Chimera, one encompassing the portal protein and the other the DUF4062 AUX domain. The *Kp*Swz$^{AUX}$ was visible as a distorted ring of density around the C12-refined portal, indicating a symmetry mismatch. 3D classification of the C12-refined structure was performed in C1 symmetry with the AUX mask. Around 300K particles were selected, yielding acceptable density for the AUX domain crescent after homogeneous refinement with relaxed C12 symmetry using an overall mask. To improve reconstruction quality, reference motion correction was performed on all 1.2M particles refined in C12 symmetry with portal-only mask. After another homogeneous refinement in C12, 3D classification without alignment in C1 symmetry was performed with AUX mask, and additional rounds of 2D classification. 1M particles containing *Kp*Swz$^{AUX}$ were kept for further refinement, and vacant portals were rejected. Particles from the initial AUX-domain refinement were substituted with the reference corrected data, yielding 274K particles at an improved resolution (2.8 Å to 2.57 Å). After CTF and non-uniform refinement with relaxed C12 symmetry, particles were separated by per-particle scale. Final non-uniform refinement was performed from ~205K particles in relaxed C12 symmetry to yield a 2.53 Å map.

We observed that the portal protein exhibited a large amount of motion, including relative wing movements in all possible directions, pore contraction and dilation, and torsions between the pore and the wings. As a result, refinements in C12 symmetry were difficult to interpret outside of the pore region, even when resolution was estimated as high as 2.1 Å. Wing domain was modelled using local refinement as a reference. Overall reconstruction was symmetry-expanded and classified with a mask focused on a single wing domain. One class with a single sharp wing density was used in overall local refinement, resulting map used as a reference for wing model building.

**MVP/TEP1 + NAD$^+$ + AMP-PNP sample**

All processing steps were performed in cryoSPARC, with CTF refinements done after reconstructions and reference motion correction. 8,103 movies in EER format with 1112 frames each were fractionated into 48 fractions with dose ~1.03 $e^-/Å^2$ and upsampling factor 2. A small subset of movies was processed first to generate initial model (C1) and a refined 3D map in D39 symmetry from 300 particles. Templates from the 3D volume were generated. After motion correction and patch CTF of the full dataset, templates were used to pick vault particles, which were subjected to 2D classification and homogeneous refinement in D39 symmetry. Attempt to fit the rat crystal structure (PDB: 4HL8) or available EM structures into the initial maps at ~3 Å resolution was unsuccessful and required re-calibration of the pixel size. Provided physical pixel size of 0.738 was adjusted to 0.716 by manual comparisons of model-map fit correlation in UCSF Chimera. The crystal structure (PDB: 4HL8) and available EM structures were a good fit, with only small differences in pixel size evident. Movies were reprocessed with the corrected pixel size, yielding ~25K particles. Particles were extracted with 1200 box size, and downsampled to 800 for pixel size of 1.074 Å, to address memory constraints on GPUs used for processing (A5000 with 24 GB of VRAM was used). Homogeneous refinement in D39 symmetry with positive Ewald sphere correction progressed close to the downsampled Nyquist frequency, yielding 2.23 Å maps. Particles were further subjected to 2D classification, and final map for the overall vault was obtained with 20,744 particles at an estimated 2.17 Å resolution.

RNA vaults were further refined to improve local features, especially the flexible waist region. Vault particles were expanded in D39 symmetry, and re-centered around the upper part of the waist (mask encompassing all visible protein around the waist), and the shoulder regions (tight mask around a trimer of MVP). Particles for either class were re-extracted with full pixel size of 0.716, local refined, and split into random batches for reference motion correction (to overcome GPU memory limits due to a large number of particles per movie). Waist region was additionally subjected to 3D classification, before final local refinements.

**MVP/PARP4/TEP1 + NADP sample**

Data was collected from a low density, and a high-density grid during a single 3-day session on a TFS Krios, as described before. Movies from each grid were pre-processed separately. Using initial 200 movies from the low-density grid, 516 particles were picked to obtain a ~3 Å reconstruction in D39 symmetry. Pixel size was re-calibrated as before, from provided 0.743 Å/px to 0.713 Å/px, bringing data to scale with the crystal, and published EM structures. Data was then processed as before.

Interestingly, attempts to fit the crystal or EM structures to the data from high-density grid initially failed. After 3D-classification, it was discovered that most of the particles in the high-density grid belonged to a larger class of the RNA vault particle, exhibiting an extended shoulder region that dilated in diameter, also resulting in an overall taller structure. Mixed-class refinement thus contained smeared density around the cap and shoulder regions. From total 53,381 particles on the high-density grid, the commonly observed “compact” vaults were a minor class with 8,388 particles, while 38,771 were the extended conformation. Remaining 6,222 particles belonged to an intermediate “medium” conformation (overall models for medium and large conformations were not built). Compact vaults from both grids were pooled (18,171 particles) and processed separately from the medium and extended vault. Reference motion correction was run separately on all classes. Particles were downsampled to pixel size of 1.0695. Final overall resolution for the compact conformation was 2.16 Å, 2.67 Å for the intermediate, and 2.24 Å for the extended conformation.

To improve the resolution around the shoulder region, compact and extended RNA vaults were separately symmetry expanded in D39, re-centered, re-extracted with full pixel size and locally refined. Masks encompassing all visible MVP protein were used for refinement of both the compact and extended vault shoulder conformations, and both models were built. Extended conformation required a large relative repositioning of the R9 and shoulder domains, with only minor internal changes within the domains. Final resolution for the compact shoulder was 1.96 Å, and 2.37 Å for the extended shoulder.

To obtain the $PARP4^{MINT}$-MVP structure, all 63,171 particles from both grids were pooled in a homogeneous refinement, symmetry expanded in D39, and 3D-classified with a mask around a single MINT domain. 1,308,658 particles contained PARP4 density, out of

4,927,416 total for an average of ~20 MINT domains/vault in the reconstituted sample. Particles were re-centered and re-extracted with full pixel size. Wide mask was applied, and 3D classification was performed to select a uniform particle class for a final local refinement to yield a 2.85 Å map.

To obtain the structure of the vault cap, 63,171 pooled particles were expanded in D1 symmetry, recentered around the cap region and re-extracted. Resulting 124,197 particles were aligned by homogeneous refinement in C39 symmetry. 3D-classification in C13 symmetry with 3 classes was performed with a mask encompassing the top of the cap. Classes were found to uniformly distribute into all 3 possible positions and were manually rotated by +/-9.23° to overlay. Particles were pooled in local refinement using C13 symmetry, and reference motion correction was performed. Particles were sorted by per-particle scale, and top 115,486 were local refined and reconstructed with positive Ewald sphere correction to obtain the final 1.92 Å map.

Despite extensive classifications, we were unable to obtain atomic reconstructions of the pore. Pore displayed high flexibility, changing its diameter between classes. Densities above the pore corresponding to the knob region were observed forming different symmetric structures on top of the pore and the cap, and within it. However, no consistent consensus was ever achieved, leading us to believe all the classes were artifacts. Density within the pore also changed appearance between the classes, moving up or down. Similar results were achieved with the other two datasets presented here.

**EMPIAR-10766 Human Brain RNA vault re-processing.**

Movies from EMPIAR-10765 did not contain any RNA vaults. However, based on dataset metadata from Lövestam & Scheres, 2025 (*28*), we found that the RNA vault dataset had the same number of movies as EMPIAR-10766. Like before, pixel size was recalibrated from 1.15 Å/px to 1.13 Å/px, this time using our MVP/PARP4/TEP1 structure for consistent comparison. Overall vaults (19,230 particles) were processed as before to focus on the shoulder region. After 3D classification and rejection of particles with low per-particle scale, remaining 464,690 particles yielded a 2.62 Å map from a local refinement. Model from MVP/PARP4/TEP1 was fit, adjusted in Coot (*29*)and refined in Phenix (*30*).

**Model building**

*Kp*Swz dimer was fit with the AlphaFold3 model into the tetrameric map and manually rebuilt in Coot. Maps for initial building were processed with deepEMhancer (*31*). Model was rebuild in ISOLDE (*32*), and refined in Phenix. Dimer was expanded into a tetramer and process was repeated. Tetrameric model was built into the hexamer map, adjusted in Coot and ISOLDE, and refined in Phenix. Solvent was added with Douse in Phenix, inspected in coot and refined in Phenix real space refinement.

*Kp*Swz$^{AUX}$-portal$^{Bas14}$ structure was based on AlphaFold3 prediction of the portal protein, and the AUX domain tetramer. Portal was rigid body fit into a C12-symmetric map. A trimer was manually fit in Chimera. Wing and stalk regions were fit separately, due to incorrect prediction of the angle. After rebuilding of the trimer in Coot and ISOLDE, the middle monomer was symmetry expanded into a C12 refined map in ChimeraX using commands "volume symmetry" to set C12 symmetry, "measure symmetry" to estimate map symmetry, "sym" to apply symmetry to the model, and "combine" to make a dodecameric PDB file. Model was fit into the C12-relaxed map, trimmed and rebuilt to fit the density in Coot and ISOLDE. *Kp*Swz$^{AUX}$ AlphaFold model was fit into the best-resolved density around the AUX crescent and rebuild in Coot. Model was rigid-body fit into the 13 densities around the portal and regularized in ISOLDE. Each AUX domain was then manually rebuilt one-by-one, focusing on the variable regions. The wing domain of the portal was compared to the density obtained from a local refinement to guide building. Final model was refined in Phenix real space refine with disabled NCS restraints.

The RNA vault (TEP1 + BAD + AMP-PNP) was built using the rat EM structure as the starting model (PDB: 7PKR). Single MVP monomer was fit into the D39 density map in Chimera and mutated to human sequence in Coot. Sharpened map was used for the cap/shoulder region and nearby MVP repeats, and unsharpened for the heterogeneous waist region. Model was fit using real space refinement in Coot and manually expanded into a hexamer in Chimera. After rebuilding, middle monomer was refined in ISOLDE and refined in Phenix Real space refine. Monomer was then expanded into a 78-mer in ChimeraX, using the same commands as for the portal, but with D39 symmetry. Overall model was refined with strict NCS in Phenix real space refine. Ligand BAD was obtained from the CIF library under the code DQV. Restraints were generated in Phenix Elbow.

The shoulder region local refinement was built from the overall model. A trimer of the shoulder was excised and fit into a sharpened map in ChimeraX. Residues were adjusted for the differences in the local refinement, including different conformers. Trimer was refined in ISOLDE and Phenix real space refine. Waters were added in Douse, cleaned up manually and final model was refined in Phenix.

The waist region local refinement was built from the overall model, similar to the waist region. Single MVP repeat from the other half of the vault was included to enable study of the interface.

The RNA vault (PARP4/TEP1 + $NADP^+$) was built from the TEP1-only MVP structure, in a similar fashion. The shoulder region of both conformations was built accordingly, with ADP-ribose built from AR6 molecule in the CIF library. ADP-ribose was observed despite adding $NADP^+$ to the sample.

The MVP cap region was built in Coot de-novo. First, trimeric asymmetric unit was manually built and expanded into C13 symmetry in ChimeraX. Trimer in the context of the full cap was rebuilt in ISOLDE, expanded into C13, and refined in Phenix real space refine.

The MINT-MVP model was predicted using AlphaFold3. Model was fit into the map, and the MVP molecule was substituted with a fragment from the overall MVP model. Model was manually rebuilt in Coot, regularized in ISOLDE and refined in Phenix real space refine. We observed additional extensive density on PARP4 Cys1622. Similar densities are often observed in vitrified CryoEM specimens (*33*).

#### TEP1$^{TIR}$ Crystallization, Data Collection and Structure Determination.

TEP1 895-1113 was in solution with 50 mM Tris 8.0, 50 mM NaCl, 1 mM DTT, 2 mM BAD (benzamide adenine dinucleotide) at 40 mg/mL. Crystals were grown by the hanging drop vapor diffusion method using a 1:1 ratio of protein to reservoir solution containing 0.2 M $MgSO_4$, 17-20% w/v PEG 3350. Crystals were cryoprotected in reservoir solution with 22% PEG 3350 and 20% ethylene glycol. Crystals diffracted to a minimum Bragg spacing ($d_{min}$) of 2.44 Å and exhibited the symmetry of space group $P4_332$ with cell dimensions of a=b=c=132.762 Å, and contained one TEP1$^{TIR}$ per asymmetric unit. Diffraction data was collected at beamline BL12-2 at the Stanford Synchrotron Radiation Lightsource (SLAC National Accelerator Laboratory, Menlo Park, California, USA). All diffraction data were processed in the program HKL-3000 (*34*). Phases were calculated via molecular replacement using AlphaFold3 output as a search model in Phaser (*35*). Modelling was performed by multiple cycles of manual rebuilding in the program Coot (*29*), ISOLDE (*32*), and refinement in the program BUSTER (*36*). Positional and isotropic atomic displacement parameter (ADP) as well as TLS ADP refinement was performed in the program Phenix with a random 10% of all data set aside for an $R_{free}$ calculation. TLS groups were generated by TLSMD server (https://doi.org/10.1107/S0021889805038987). Data collection and structure refinement statistics are summarized in **Table S2.**

#### Cryo-Electron Tomography Sample Preparation

Reconstituted vault particles containing MVP, MVP–TEP1, or MVP–PARP4 were prepared as described above. Quantifoil R2/2 copper–carbon grids (Electron Microscopy Sciences) were glow discharged at -30 mA for 80 s. A 4 µL aliquot of the reconstituted sample, mixed with 10 nm colloidal gold fiducial markers, was applied to each grid. Excess liquid was removed by double-sided blotting in a Vitrobot (Thermo Fisher Scientific) using a blot force of -10 and a blot time of 3 s. Grids were plunge-frozen rapidly in liquid ethane cooled by liquid nitrogen. Frozen grids were stored and handled under liquid nitrogen throughout all subsequent procedures.

#### Cryo-Electron Tomography Data Acquisition and Processing

Frozen grids were transferred to a Titan Krios transmission electron microscope (Thermo Fisher Scientific) operating at 300 kV for cryo-electron tomography data collection. Tilt-series images were recorded on a Gatan K3 direct electron detector equipped with a BioQuantum energy filter (20 eV slit width) and a Volta phase plate (VPP), using a target defocus range of −0.5 to −0.75 µm.

Data were acquired using a dose-symmetric tilt scheme (*37*)implemented through PaceTOMO (*38*) and SerialEM (*39*). Images were collected every 5° over a ±60° tilt range. The nominal magnification was 26,000×, corresponding to a calibrated pixel size of 3.44 Å/pixel. The total accumulated electron dose per tilt series was approximately 130 $e^-/Å^2$, corresponding to ~4.8 $e^-/Å^2$ per tilt.

Tilt-series alignment was performed in IMOD (v4.11.25) using the fiducial marker-based pipeline, and tomograms were reconstructed by weighted back projection (*40*). Tomograms were denoised using IMOD's non-linear anisotropic diffusion (NAD) filtering.

Initial vault particle positions and orientations were determined in IMOD using the stalkInit procedure. Briefly, two points were placed per particle at the centroid and apex, and stalkInit was used to generate an initial motive list aligning vault particles along a common vertical axis. Angular refinement and subtomogram averaging were performed in PEET (v1.17.0), yielding averaged density maps for MVP, MVP–TEP1, and MVP–PARP4 vault particles (**Table S5)**

Particles contributing to each subtomogram average were split into even and odd half-sets and aligned independently. Resolution was estimated using PEET's calcUnbiasedFSC program applied to the resulting half-maps, and values are reported according to the 0.5 Fourier shell correlation (FSC) criterion. Isosurface renderings were generated in UCSF ChimeraX. Alignment and overlay of MVP–TEP1 and MVP–PARP4 averages were performed using Chimera's "Fit in Volume" function.

### Tomogram Segmentation

To analyze the spatial distribution of interior vault densities, a custom three-class U-Net convolutional neural network was trained in Dragonfly (v2022.2) to classify voxels as background, vault interior, or vault exterior. The network was trained on 26 manually annotated tomogram slices following the neural network segmentation workflow described (*41*).

The trained model was applied to full tomograms to segment vault particles across each field of view. All automated segmentations were subsequently reviewed and manually refined in Dragonfly to correct misclassified voxels. Segmented volumes were rendered in Dragonfly to generate the three-dimensional models shown in the figures.

### Artificial Intelligence (AI)

ChatGPT was occasionally used solely to improve sentence readability and to check grammar and spelling. No content, interpretation, or conclusions were generated by AI.

Table S1. Cryo-EM data collection, refinement and validation statistics for KpSwz apo and portal complex

| **Dataset:** | ***Kp*Swz$^{15\text{-C, E97A}}$** | | **portal$^{Bas14}$ + *Kp*Swz$^{15\text{-C, E97A}}$** |
|---|---|---|---|
| **Data collection and processing** | Tetramer 11CV / EMD-75626 | Hexamer 10WC / EMD-75498 | Portal$^{Bas14}$ + *Kp*Swz$^{AUX}$ 11CQ / EMD-75622 |
| Magnification (x$10^3$) | 105 | | 105 |
| Voltage (kV) | 300 | | 300 |
| Electron exposure (e$^-$/Å$^2$) | 52 | | 50 |
| Defocus range (μm) | 1-2.4 | | 0.9-2.2 |
| Pixel size (Å) | 0.83 | | 0.826 |
| Symmetry imposed | C2 | | C1 (Relaxed C12) |
| Initial particle images (no.) | ~1.3M | | 1.2M |
| Final particle images (no.) | 84,278 | 53,724 | 204,790 |
| Map resolution (Å) | 2.66 | 2.42 | 2.53 |
| FSC threshold: **0.143** | | | |
| **Refinement** | | | |
| Initial model used (PDB code) | AlphaFold | | AlphaFold |
| Model resolution (Å) | 2.8 | 2.7 | 2.9 |
| FSC threshold: **0.5** | | | |
| Map sharpening *B* factor (Å$^2$) | 84.5 | 58.2 | 70 |
| Model composition | | | |
| Non-hydrogen atoms | 5,688 | 8,501 | 46,595 |
| Protein residues | 692 | 1,032 | 5944 |
| *B* factors (mean, Å$^2$) | | | |
| Protein | 68.87 | 83.63 | 96.05 |
| Metal | | | 103.14 |
| R.m.s. deviations | | | |
| Bond lengths (Å) | 0.005 | 0.003 | 0.001 |
| Bond angles (°) | 0.520 | 0.452 | 0.389 |
| Validation | | | |
| MolProbity score | 1.29 | 1.31 | 0.99 |
| Clashscore | 2.77 | 3.83 | 2.18 |
| Poor rotamers (%) | 0 | 0 | 0.24 |
| Ramachandran plot | | | |
| Favored (%) | 96.56 | 97.29 | 98.16 |
| Allowed (%) | 3.44 | 2.71 | 1.84 |
| Disallowed (%) | 0 | 0 | 0 |

Table S2. $TEP1^{TIR}$ X-ray data and model statistics.

| Data collection (TEP1 895-1113 crystal) | |
|---|---|
| Wavelength (Å) | 0.9795 |
| Space group | P 43 3 2 |
| Cell dimensions | |
| *a, b, c* (Å) | 132.762, 132.762, 132.762 |
| *α, β, γ* (°) | 90.00, 90.00, 90.00 |
| Number of reflections measured | 15,346,513 |
| Number of unique reflections | 15058 (981) |
| Resolution (Å) | 40.03 - 2.438 (2.52 - 2.44) |
| $R_{pim}$ | 0.011 |
| $CC_{1/2}$ | 1.00 |
| Overall I/s(I) | 51.8 |
| Completeness (%) | 97.07 (70.68) |
| Redundancy | 58.5 (30.5) |
| Wilson B ($Å^2$) | 35.26 |
| **Refinement** | |
| Resolution (Å) | 40.03 - 2.438 (2.52 - 2.44) |
| Number of reflections<br>Reflection test set | 15058 (981)<br>1505 (98) |
| $R_{work}/R_{free}$ | 0.2372/0.2876 |
| Number of non-hydrogen atoms | |
| All | 1941 |
| Protein | 1791 |
| Non-water solvent | 13 |
| Water | 137 |
| Average *B* factors ($Å^2$) | |
| All | 69.41 |
| Protein | 70.79 |
| Ligand | 70.54 |
| Water | 51.28 |
| RMSD | |
| Bond lengths (Å) | 0.006 |
| Bond angles (°) | 0.67 |
| Ramachandran statistics | |
| Favored regions (%) | 98.17 |
| Allowed regions (%) | 1.83 |
| Outliers (%) | 0.00 |
| Rotamer outliers (%) | 0.52 |
| *MolProbity* overall score | 1.13 |

### Table S3. Cryo-EM data collection, refinement and validation statistics for MVP/TEP1 samples

| **Dataset:** | **MVP/TEP1 + BAD + AMP-PNP** | | |
|---|---|---|---|
| **Data collection and processing** | Overall (BAD) 11EQ EMD-75656 | Shoulder (BAD) 11EE EMD-75647 | Waist (half interface) 11DV EMD-75642 |
| Magnification (x$10^3$) | 165 | | |
| Voltage (kV) | 300 | | |
| Electron exposure ($e^-/Å^2$) | 50 | | |
| Defocus range (μm) | 0.9-2.2 | | |
| Pixel size (Å) | 0.738 (calibrated to 0.716) | | |
| Symmetry imposed | D39 | C1 (symm expand) | C1 (symm expand) |
| Initial particle images (no.) | 25,542 | | |
| Final particle images (no.) | 20,744 | 1,464,614 | 1,016,071 |
| Map resolution (Å) | 2.17 | 1.87 | 2.33 |
| FSC threshold: **0.143** | | | |
| **Refinement** | | | |
| Initial model used (PDB code) | 4HL8 | - | - |
| Model resolution (Å) | 2.5 | 1.9 | 2.7 |
| FSC threshold: **0.5** | | | |
| Map sharpening *B* factor ($Å^2$) | 50.3 | 47.3 | 75 |
| Model composition | | | |
| Non-hydrogen atoms | 476,268 | 6,888 | 9,477 |
| Protein residues | 59,592 | 729 | 1,125 |
| Ligands | 78 | 3 | - |
| *B* factors (mean, $Å^2$) | | | |
| Protein | 41.74 | 12.41 | 60.35 |
| Ligand | 36.17 | 19.96 | - |
| R.m.s. deviations | | | |
| Bond lengths (Å) | 0.002 | 0.003 | 0.005 |
| Bond angles (°) | 0.399 | 0.535 | 0.509 |
| Validation | | | |
| MolProbity score | 1 | 1.19 | 1.31 |
| Clashscore | 2.21 | 4.09 | 3.12 |
| Poor rotamers (%) | 0.15 | 0.48 | 0 |
| Ramachandran plot | | | |
| Favored (%) | 98.27 | 98.45 | 96.68 |
| Allowed (%) | 1.73 | 1.55 | 3.32 |
| Disallowed (%) | 0 | 0 | 0 |

### Table S4. Cryo-EM data collection, refinement and validation statistics for MVP/PARP4/TEP1, and the Human Brain sample.

| **Dataset:** | **MVP/PARP4/TEP1 + NADP⁺** | | | | | | | **Human Brain EMPIAR-10766** |
|---|---|---|---|---|---|---|---|---|
| | Overall (ADPR) 11JB EMD-75731 | intermediate EMD-75744 | Extended EMD-75745 | Cap 11FH EMD-75664 | Shoulder (compact) 11JC EMD-75732 | Shoulder (extended) 11JD EMD-75733 | MINT-MVP 11JF EMD-75735 | Shoulder (brain) 11EE EMD-75647 |
| **Data collection and processing** | | | | | | | | |
| Magnification (x1,000) | 165 | | | | | | | |
| Voltage (kV) | 300 | | | | | | | 300 |
| Electron exposure ($e^{-}/Å^2$) | 50 | | | | | | | 50 |
| Defocus range (μm) | 0.9-2.2 | | | | | | | 1.5-2.5 |
| Pixel size (Å) | 0.743 (calibrated to 0.713) | | | | | | | 1.15 (1.13) |
| Symmetry imposed | D39 | D39 | D39 | C13 | C1 (exp) | C1 (exp) | C1 (exp) | C1 (exp) |
| Initial particle images (no.) | 63,171 | | | | | | | 13,280 |
| Final particle images (no.) | 16,218 | | | 115,486 | 646,476 | 3,024,138 | 649,317 | 464,690 |
| Map resolution (Å) | 2.16 | 2.67 | 2.24 | 1.92 | 1.96 | 2.37 | 2.85 | 2.62 |
| FSC threshold: **0.143** | | | | | | | | |
| **Refinement** | | | | | | | | |
| Initial model used (PDB code) | - | - | - | - | - | - | AlphaFold3 | - |
| Model resolution (Å) | 2.5 | - | - | 2.1 | 2.0 | 2.5 | 3.1 | 2.7 |
| FSC threshold: **0.5** | | | | | | | | |
| Map sharpening *B* factor ($Å^2$) | 48.8 | 56.1 | 57.1 | 43.7 | 47.3 | 89.6 | 109.9 | 83.5 |
| Model composition | | | | | | | | |
| Non-hydrogen atoms | 481,962 | | | 24,427 | 6,895 | 6,324 | 5,717 | 6,123 |
| Protein residues | 60,450 | | | 3185 | 762 | 733 | 683 | 762 |
| Ligands | 78 | | | - | 3 | 3 | - | 3 |
| *B* factors (mean, $Å^2$) | | | | | | | | |
| Protein | 44.61 | | | 32.0 | 24.07 | 34.97 | 76.1 | 50.98 |
| Ligand | 47.3 | | | - | 30.98 | 48.92 | - | 73.3 |
| R.m.s. deviations | | | | | | | | |
| Bond lengths (Å) | 0.002 | | | 0.003 | 0.003 | 0.003 | 0.002 | 0.004 |
| Bond angles (°) | 0.443 | | | 0.436 | 0.486 | 0.447 | 0.407 | 0.498 |
| Validation | | | | | | | | |
| MolProbity score | 1.20 | | | 0.98 | 1.37 | 1.21 | 1.18 | 1.12 |
| Clashscore | 4.20 | | | 2.12 | 6.65 | 4.24 | 1.99 | 2.30 |
| Poor rotamers (%) | 0.15 | | | 0 | 0.61 | 0 | 0 | 0 |
| Ramachandran plot | | | | | | | | |
| Favored (%) | 98.23 | | | 99.58 | 100 | 99.44 | 96.6 | 97.45 |
| Allowed (%) | 1.77 | | | 0.42 | 0 | 0.56 | 3.40 | 2.55 |
| Disallowed (%) | 0 | | | 0 | 0 | 0 | 0 | 0 |

**Table S5. Summary of Cryo-ET data**

| Protein | EMDB Accession No. | Tomograms | Particles | Resolution (nm) 0.5 FSC |
|---|---|---|---|---|
| MVP | D_1000304660 | 6 | 502 | 2.8 |
| MVP-PARP4 | D_1000304609 | 7 | 503 | 3.2 |
| MVP-TEP1 | D_1000304631 | 14 | 518 | 3.2 |

### References


1. S. El-Gebali *et al.*, The Pfam protein families database in 2019. *Nucleic Acids Res* **47**, D427-D432 (2019).
2. D. Xu, L. Jaroszewski, Z. Li, A. Godzik, FFAS-3D: improving fold recognition by including optimized structural features and template re-ranking. *Bioinformatics* **30**, 660-667 (2014).
3. R. D. Schaeffer *et al.*, ECOD: integrating classifications of protein domains from experimental and predicted structures. *Nucleic Acids Res* **53**, D411-D418 (2025).
4. L. Zimmermann *et al.*, A Completely Reimplemented MPI Bioinformatics Toolkit with a New HHpred Server at its Core. *J Mol Biol* **430**, 2237-2243 (2018).
5. K. Essuman *et al.*, TIR Domain Proteins Are an Ancient Family of NAD(+)-Consuming Enzymes. *Curr Biol* **28**, 421-430 e424 (2018).
6. M. Hauser, M. Steinegger, J. Soding, MMseqs software suite for fast and deep clustering and searching of large protein sequence sets. *Bioinformatics* **32**, 1323-1330 (2016).
7. J. Rozewicki, S. Li, K. M. Amada, D. M. Standley, K. Katoh, MAFFT-DASH: integrated protein sequence and structural alignment. *Nucleic Acids Res* **47**, W5-W10 (2019).
8. S. C. Potter *et al.*, HMMER web server: 2018 update. *Nucleic Acids Res* **46**, W200-W204 (2018).
9. T. Frickey, A. Lupas, CLANS: a Java application for visualizing protein families based on pairwise similarity. *Bioinformatics* **20**, 3702-3704 (2004).
10. J. L. Steenwyk, T. J. Buida, 3rd, Y. Li, X. X. Shen, A. Rokas, ClipKIT: A multiple sequence alignment trimming software for accurate phylogenomic inference. *PLoS Biol* **18**, e3001007 (2020).
11. B. Q. Minh *et al.*, IQ-TREE 2: New Models and Efficient Methods for Phylogenetic Inference in the Genomic Era. *Molecular biology and evolution* **37**, 1530-1534 (2020).
12. I. Letunic, P. Bork, Interactive Tree of Life (iTOL) v6: recent updates to the phylogenetic tree display and annotation tool. *Nucleic Acids Res* **52**, W78-W82 (2024).
13. T. Goldfarb *et al.*, NCBI RefSeq: reference sequence standards through 25 years of curation and annotation. *Nucleic Acids Res* **53**, D243-D257 (2025).
14. N. A. O'Leary *et al.*, Exploring and retrieving sequence and metadata for species across the tree of life with NCBI Datasets. *Sci Data* **11**, 732 (2024).
15. F. Tesson *et al.*, Systematic and quantitative view of the antiviral arsenal of prokaryotes. *Nat Commun* **13**, 2561 (2022).
16. A. Beavogui *et al.*, The defensome of complex bacterial communities. *Nat Commun* **15**, 2146 (2024).
17. M. T. Edwards, S. C. Rison, N. G. Stoker, L. Wernisch, A universally applicable method of operon map prediction on minimally annotated genomes using conserved genomic context. *Nucleic Acids Res* **33**, 3253-3262 (2005).

18. B. Taboada, K. Estrada, R. Ciria, E. Merino, Operon-mapper: a web server for precise operon identification in bacterial and archaeal genomes. *Bioinformatics* **34**, 4118-4120 (2018).
19. G. E. Crooks, G. Hon, J. M. Chandonia, S. E. Brenner, WebLogo: a sequence logo generator. *Genome Res* **14**, 1188-1190 (2004).
20. J. Abramson *et al.*, Accurate structure prediction of biomolecular interactions with AlphaFold 3. *Nature*, (2024).
21. C. L. M. Gilchrist, M. Mirdita, M. Steinegger, Multiple Protein Structure Alignment at Scale with FoldMason. *bioRxiv*, 2024.2008.2001.606130 (2024).
22. P. J. Shilling *et al.*, Improved designs for pET expression plasmids increase protein production yield in Escherichia coli. *Commun Biol* **3**, 214 (2020).
23. D. E. Deatherage, J. E. Barrick, Identification of mutations in laboratory-evolved microbes from next-generation sequencing data using breseq. *Methods Mol Biol* **1151**, 165-188 (2014).
24. T. A. Eastwood *et al.*, High-yield vesicle-packaged recombinant protein production from E. coli. *Cell Rep Methods* **3**, 100396 (2023).
25. E. L. Aronovich, R. S. McIvor, P. B. Hackett, The Sleeping Beauty transposon system: a non-viral vector for gene therapy. *Hum Mol Genet* **20**, R14-20 (2011).
26. N. Hananya, X. Ye, S. Koren, T. W. Muir, A genetically encoded photoproximity labeling approach for mapping protein territories. *Proc Natl Acad Sci U S A* **120**, e2219339120 (2023).
27. M. C. Rivera, B. Maguire, J. A. Lake, Dissociation of ribosomes into large and small subunits. *Cold Spring Harb Protoc* **2015**, 363-367 (2015).
28. S. Lovestam, S. H. W. Scheres, Cryo-EM structure of the vault from human brain reveals symmetry mismatch at its caps. *Structure* **33**, 1643-1648 e1641 (2025).
29. P. Emsley, B. Lohkamp, W. G. Scott, K. Cowtan, Features and development of Coot. *Acta Crystallogr D Biol Crystallogr* **66**, 486-501 (2010).
30. P. D. Adams *et al.*, PHENIX: a comprehensive Python-based system for macromolecular structure solution. *Acta Crystallogr D Biol Crystallogr* **66**, 213-221 (2010).
31. R. Sanchez-Garcia *et al.*, DeepEMhancer: a deep learning solution for cryo-EM volume post-processing. *bioRxiv*, 2020.2006.2012.148296 (2020).
32. T. I. Croll, ISOLDE: a physically realistic environment for model building into low-resolution electron-density maps. *Acta Crystallogr D Struct Biol* **74**, 519-530 (2018).
33. D. P. Klebl, Y. Wang, F. Sobott, R. F. Thompson, S. P. Muench, It started with a Cys: Spontaneous cysteine modification during cryo-EM grid preparation. *Front Mol Biosci* **9**, 945772 (2022).
34. W. Minor, M. Cymborowski, Z. Otwinowski, M. Chruszcz, HKL-3000: the integration of data reduction and structure solution--from diffraction images to an initial model in minutes. *Acta Crystallogr D Biol Crystallogr* **62**, 859-866 (2006).
35. A. J. McCoy *et al.*, Phaser crystallographic software. *J Appl Crystallogr* **40**, 658-674 (2007).
36. G. Bricogne *et al.*, BUSTER version X.Y.Z. *Cambridge, United Kingdom: Global Phasing Ltd*, (2017).

37. W. J. H. Hagen, W. Wan, J. A. G. Briggs, Implementation of a cryo-electron tomography tilt-scheme optimized for high resolution subtomogram averaging. *J Struct Biol* **197**, 191-198 (2017).
38. F. Eisenstein *et al.*, Parallel cryo electron tomography on in situ lamellae. *Nat Methods* **20**, 131-138 (2023).
39. D. N. Mastronarde, Automated electron microscope tomography using robust prediction of specimen movements. *J Struct Biol* **152**, 36-51 (2005).
40. J. R. Kremer, D. N. Mastronarde, J. R. McIntosh, Computer visualization of three-dimensional image data using IMOD. *J Struct Biol* **116**, 71-76 (1996).
41. J. E. Heebner *et al.*, Deep Learning-Based Segmentation of Cryo-Electron Tomograms. *J Vis Exp*, (2022).
